# Site-specific free energy surface parameters from single-molecule fluorescence measurements of exciton-coupled (iCy3)_2_ dimer probes positioned at DNA replication fork junctions

**DOI:** 10.1101/2024.10.27.620477

**Authors:** Jack Maurer, Claire S. Albrecht, Peter H. von Hippel, Andrew H. Marcus

## Abstract

Single-stranded (ss) – double-stranded (ds) DNA replication forks and primer-template junctions are important recognition sites for the assembly and function of proteins involved in DNA replication, recombination and repair. DNA ‘breathing’ – i.e., thermally-induced local fluctuations of the sugar-phosphate backbones and bases – can populate metastable conformational macrostates at positions near such junctions and likely play key roles in the functional interactions of the regulatory proteins that bind at these sites. Recently, Maurer *et al*. (1) performed polarization-sweep single-molecule fluorescence (PS-SMF) studies on exciton-coupled (iCy3)_2_ dimer-labeled ss-dsDNA fork constructs, which revealed that the nucleobases and backbones immediately adjacent to the dimer probes undergo conformational fluctuations on time scales ranging from hundreds-of-microseconds to hundreds-of-milliseconds. The local conformations sensed by the dimer probes consist of four quasi-stable macrostates whose populations and dynamics depend on dimer probe position relative to these junctions. Here we present theoretical analyses of these PS-SMF data that quantify the relative stabilities and activation barriers of the free energy surfaces (FESs) of site-specific DNA ‘breathing’ events at key positions within these junctions. Our results suggest a detailed molecular picture for DNA ‘breathing’ at these positions, thus providing insights into understanding the molecular mechanisms of the proteins that operate at these sites.

## 1. Introduction

Determining the relative stabilities and free energies of activation for protein-DNA interactions lies at the heart of understanding the functional mechanisms of biomolecular complexes responsible for gene regulation and expression. For example, during DNA replication, helicase proteins must bind to and form a stable complex with the DNA replication fork to facilitate double-stranded (ds) DNA unwinding, which is required to expose the single-stranded (ss) DNA template sequences used by the replication polymerases to synthesize daughter strands. The assembly mechanisms of such multi-component protein-DNA complexes involve a cooperative sequence of chemical steps in which the protein and nucleic acid components fluctuate between different conformations (or macrostates), ultimately leading to the assembly of functional regulatory macromolecular complexes at defined binding sites at or near ss-dsDNA fork junctions (2,3).

A useful way to understand and parameterize these cooperative assembly mechanisms is via the construction of Gibbs free energy surfaces (FES), which describe variations in the stability of protein and nucleic acid components as a function of the relevant chemical reaction coordinates. Information embedded in such FESs can reveal the role(s) of DNA ‘breathing’ at the specific regulatory sites near the ss-dsDNA fork junctions at which these regulatory protein-DNA complexes bind, assemble and operate. DNA ‘breathing’ in this context refers to the thermal activation of metastable (non-canonical) local conformations of the bases and sugar-phosphate backbones at binding sites for the protein-DNA intermediates that lie along the assembly pathways of higher order complexes (4). Points on the FES corresponding to free energy minima and maxima define the local conformations of stable macrostates and unstable transition states, respectively, while the thermodynamic ‘basins’ that surround each local minimum of the FES are associated with the distributions of quasi-degenerate states, which define the conformational disorder of the various macrostates.

In this work we apply a novel polarization-sweep single-molecule fluorescence (PS-SMF) microscopy and analysis method to determine the local conformational fluctuations – and the associated FESs – of positions within model DNA fork constructs that lie at and near the fork junctions at which replication and transcription proteins function (1). These DNA constructs have been labeled with pairs of cyanine dyes, which are inserted ‘internally’ (termed ‘iCy3’) into the sugar-phosphate backbones at defined positions relative to the ss-dsDNA fork junction (see Fig. 1*A*). By annealing two complementary single strands of DNA with opposed iCy3 labeling positions, an exciton-coupled (iCy3)_2_ dimer-labeled DNA backbone probe can be formed at a predetermined position within a DNA fork construct (Fig. 1*B*) (5-7). The optical transitions of the coupled (iCy3)_2_ dimer are coherent superpositions [symmetric (+) and anti-symmetric (–)] of electronic-vibrational (vibronic) product states. In Fig. 1*C* are illustrated the + / – electric dipole transition moments (EDTMs), 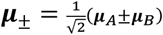,which are orthogonally polarized. In general, the magnitudes of the EDTMs depend sensitively on the local conformations of the dimer probe, which can be approximately characterized by the tilt angle, *θ*_*AB*_, the twist angle, *ϕ*_*AB*_, and the distance, *R*_*AB*_, between the two iCy3 monomers. Such (iCy3)_2_ dimer-labeled ss-dsDNA fork constructs can be used to ‘sense’ the local environment of the DNA bases and backbones within the immediate vicinity of the dimer probe (1,6-9).

**Figure 1.**
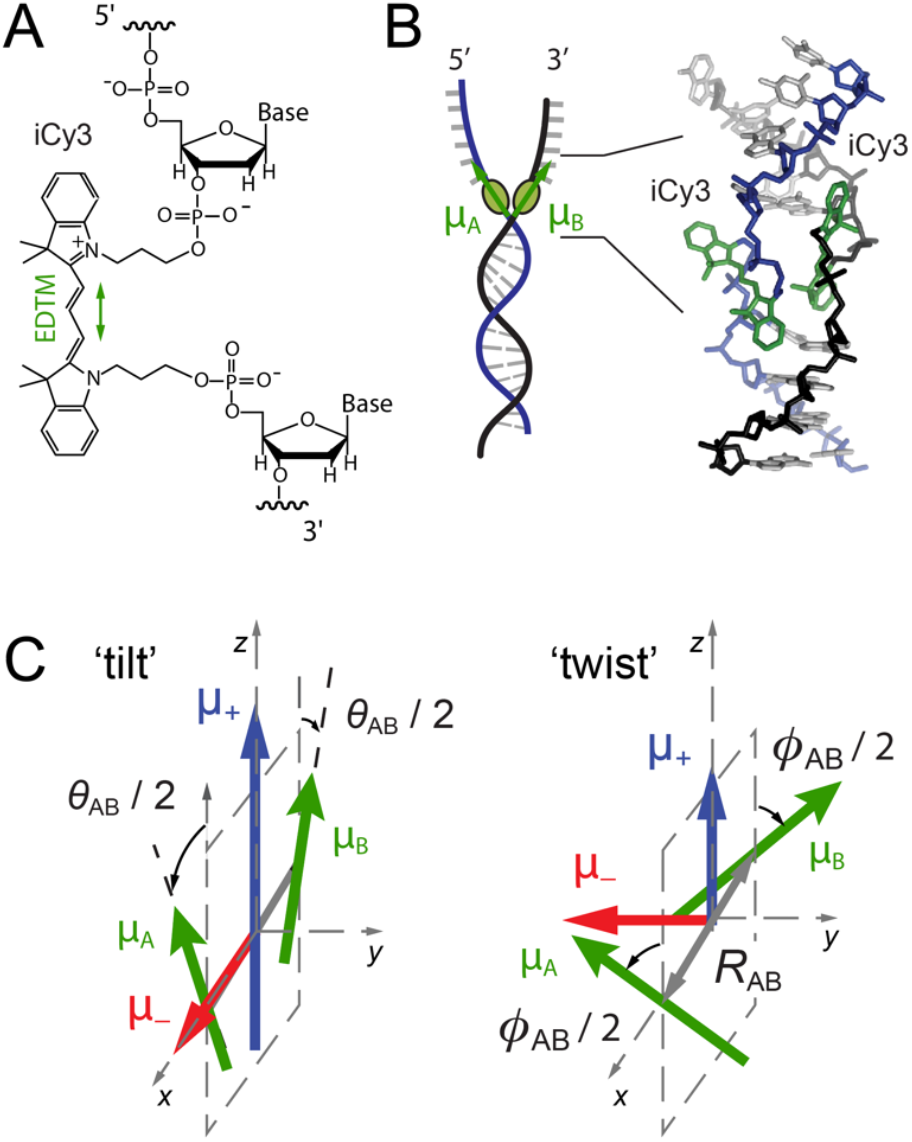
(***A***) Molecular structure and insertion chemistry of the iCy3 chromophore in DNA. The electric dipole transition moment (EDTM) is indicated by the green double-headed arrow. (***B***) Internal (iCy3)_2_ dimer probes are covalently attached within the sugar-phosphate backbones of ss-dsDNA fork constructs. (***C***) Symmetric (+, blue) and anti-symmetric (™, red) transition dipole moments depend on the local conformation of the dimer probe (green). Reproduced with permission from (9). Copyright 2022 American Institute of Physics.

In recent ensemble spectroscopic experiments by Heussman *et al*. (7-9) the *average* local conformations and conformational disorder were determined for (iCy3)_2_ dimer probes at selected positions within ss-dsDNA fork constructs. These studies employed quantum mechanical models to interpret absorbance, CD and two-dimensional fluorescence spectroscopy (2DFS) data. A summary of the ensemble results is presented in Fig. 2, which depicts the mean local conformations of the probes at positions moving across the ss-dsDNA junction. The mean local conformation of the (iCy3)_2_ dimer probe changes systematically as the labeling site is varied across the ss-dsDNA junction from the +2 to the -2 position (Fig. 2*A*). For positive integer positions, the mean local conformation is right-handed (defined as adopting a twist angle *ϕ*_*AB*_ < 90°), characteristic of the right-handed ‘Watson-Crick’ (B-form) structure, and for negative integer positions the mean local conformation is left-handed (*ϕ*_*AB*_ > 90°). Furthermore, 2DFS experiments indicate that these sites exhibit only moderate levels of structural heterogeneity (9), suggesting that the distributions of conformational macrostates are relatively narrow at positions moving across the ss-dsDNA junction towards the single-stranded regions.

**Figure 2.**
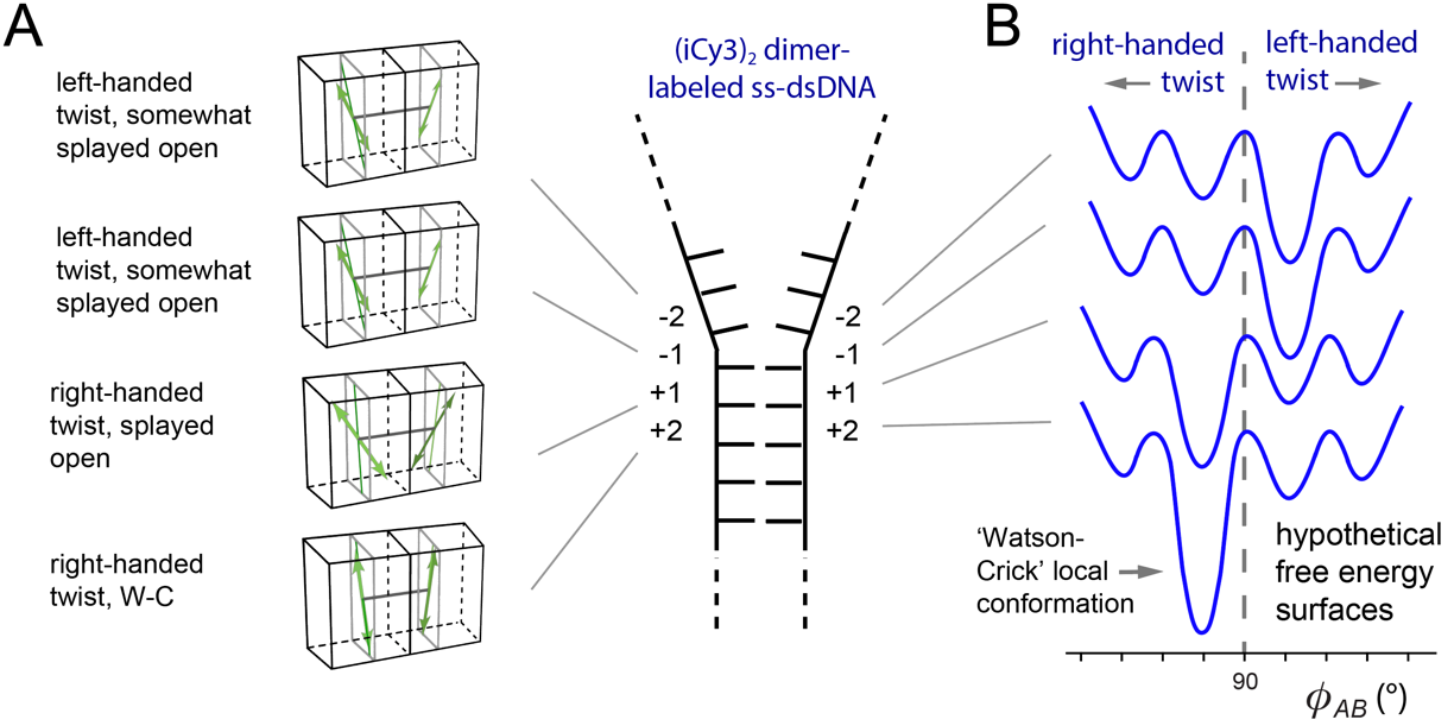
(***A***) Schematic of position-dependent changes of the mean local conformations of (iCy3)_2_ dimer-labeled ss-dsDNA constructs. The insertion site position of the (iCy3)_2_ dimer probe relative to the ss-dsDNA fork junction is indicated using positive and negative integers, as shown. (***B***) Hypothetical free energy surfaces (FESs) specific to probe-labeling positions. Individual surfaces are vertically displaced for clarity. Reproduced from (2).

A likely interpretation of the ensemble results of Heussman *et al*. is that the (iCy3)_2_ dimer probe, which is covalently linked to the sugar-phosphate backbones and bases immediately adjacent to the probe, is limited to a small number of local right- or left-handed conformations with relative stabilities that depend on the probe labeling position relative to the ss-dsDNA fork junction. Figure 2*B* depicts hypothetical FESs to illustrate this point. Each surface contains four possible local conformations of the (iCy3)_2_ dimer probe, which may exhibit either a right-handed or a left-handed twist. In this picture, the right-handed ‘Watson-Crick’ conformation is thermodynamically favored at positive integer positions, while a non-canonical left-handed conformation is favored at negative integer positions.

We here present an FES interpretation of our recent PS-SMF experiments (1), which directly monitor the local conformational fluctuations of the (iCy3)_2_ dimer probe as a function of its insertion positions relative to the ss-dsDNA fork junction. These measurements exploit the sensitivity of the conformation-dependent coherent coupling between the closely spaced iCy3 monomers within (iCy3)_2_ dimer-labeled ss-dsDNA constructs. Our results indicate that the bases and sugar-phosphate backbones ‘sensed’ by the (iCy3)_2_ dimer probes can adopt four quasi-stable macrostates, whose relative stabilities and transition barriers depend on the insertion position at which the dimer probes are placed within the ss-dsDNA fork junction (1). The presence of four quasi-stable macrostates is corroborated by recent MD simulation studies of these systems (10). In addition, the energetics of the (iCy3)_2_ dimer-labeled ss-dsDNA constructs can be systematically varied by increasing or decreasing salt concentrations relative to the ‘physiological’ conditions that we initially employed (11-15).^1^ In the current work we apply a four-state kinetic network model to analyze our PS-SMF data by implementing a multi-parameter optimization calculation using a high-performance computer cluster. We thus quantify the thermodynamic minima and the free energy barriers of relevant conformational macrostates from which we visualize the principal features of the FES. We note that our findings are consistent with those of the early hydrogen exchange experiments from which the structural concepts of DNA ‘breathing’ were originally proposed (16-19). Our results provide a more detailed and specific picture of DNA ‘breathing’ at positions near ss-dsDNA fork junctions, from which we may begin to understand the assembly and functional mechanisms of gene regulatory protein-DNA complexes.

## 2. Materials and Methods

### 2.1 Sample preparation

The DNA sequences and sample preparation methods used to collect the data for these studies were reported previously (1). Briefly, the (iCy3)_2_ dimer-labeled ss-dsDNA fork constructs were prepared taking special care to properly anneal complementary strands in aqueous buffer salt solution (100 mM NaCl, 6 mM MgCl_2_, and 10 mM Tris, pH 8.0). The specific nucleic acid constructs used in the present study are listed in Table I. The annealed ss-dsDNA constructs were diluted to ∼100 pM in solvent environments covering a range of salt concentrations, and then introduced into a custom-built microscope sample chamber constructed from quartz slides and coverslips. The inner surfaces of the sample chamber were coated with a layer of poly(ethylene glycol) (PEG), which was sparsely labeled with biotin (20). Neutravidin was used as a linker to bind the biotin-labeled PEG to the biotin molecules covalently attached to the 5’ ends of the dsDNA regions of the ss-dsDNA constructs, as shown in Table 1 and in Fig. 3*A*.

**Table 1.**
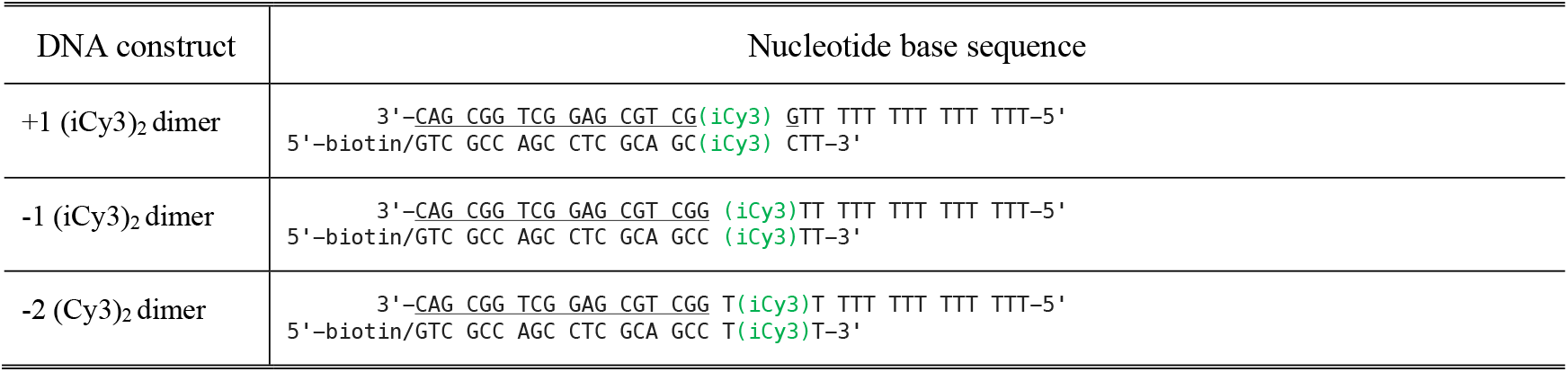
Base sequences and nomenclature for the (iCy3)_2_ dimer-labeled ss-dsDNA fork constructs used in these studies. The horizontal lines indicate regions of complementary base pairing.

**Figure 3.**
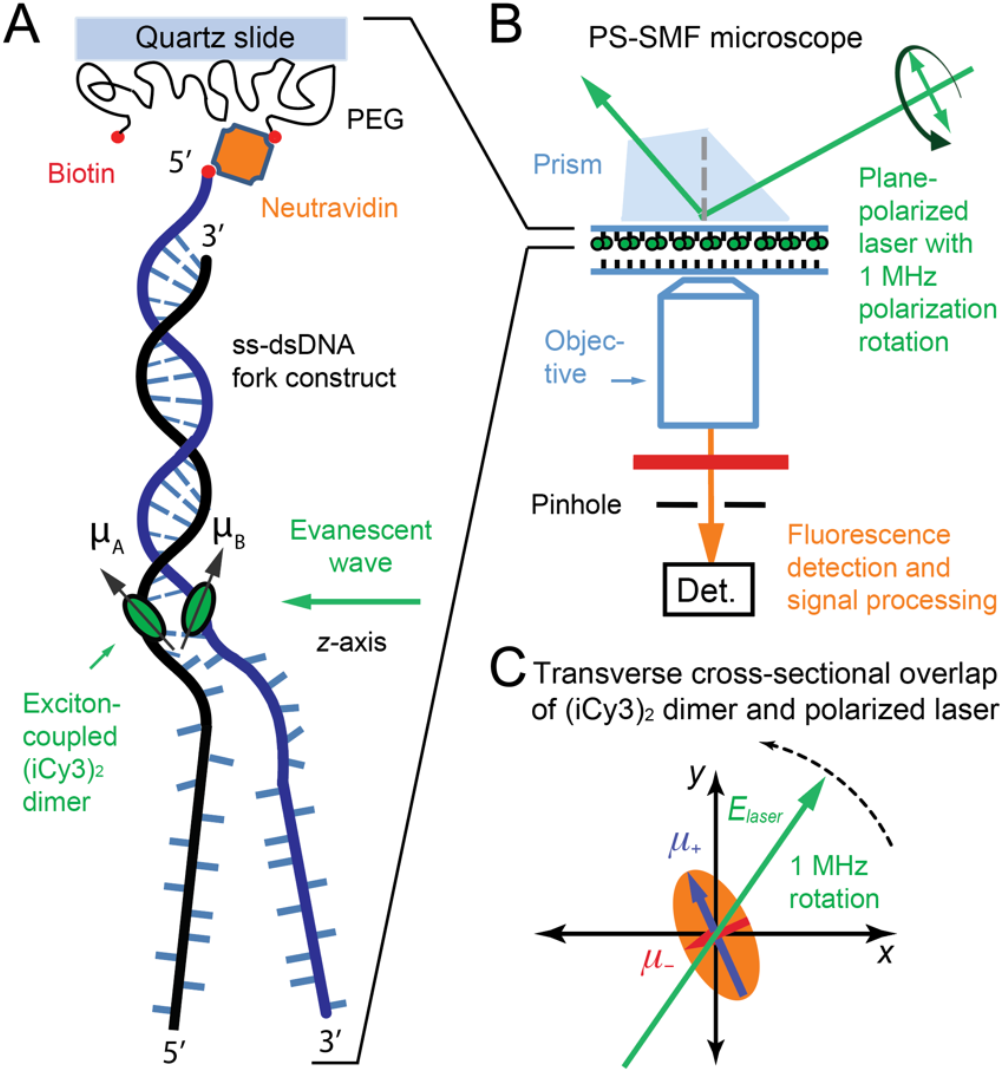
Schematic of PS-SMF microscopy and measurement principles. (***A***) (iCy3)_2_ dimer-labeled ss-dsDNA fork constructs are attached to a microscope slide using biotin/neutravidin linkers. (***B***) A total internal reflection fluorescence (TIRF) illumination geometry is used to preferentially excite molecules attached near the liquid-solid interface. The source laser generates an evanescent wave that propagates parallel to the sample surface with its electric field vector contained in a plane normal to the surface. The electric field vector is rotated continuously at a frequency of 1 MHz by sweeping the phase of an interferometer. (***C***) The symmetric (+) and anti-symmetric (™) EDTMs of the (iCy3)_2_ dimer probe (labeled **μ**_±_, and shown as blue and red vectors, respectively) define major and minor axes of a ‘polarization ellipse’ in the transverse cross-sectional area of the incident laser beam. Reproduced with permission from (1). Copyright 2023 American Chemical Society.

### 2.2 Polarization-sweep single-molecule fluorescence (PS-SMF) microscopy

The PS-SMF method was recently described in detail (1). A schematic of the PS-SMF microscope is shown in Fig. 3. The (iCy3)_2_ dimer-labeled ss-dsDNA fork constructs are attached to the inner surface of the quartz sample chamber (Fig. 3*A*), which is placed on the stage of an inverted total internal reflection fluorescence (TIRF) microscope (Fig. 3*B*). The sample is illuminated using the evanescent field of a continuous wave (cw) 532 nm laser, which propagates parallel to the slide surface with its electric field vector contained within a plane normal to the surface. The plane polarization state of the laser is established using a polarized Mach-Zehnder interferometer and quarter-wave plate, and the electric field vector is rotated continuously at a 1 MHz frequency by sweeping the phase of the interferometer using acousto-optic Bragg cells (1). The symmetric (+) and anti-symmetric (™) excitons of the (iCy3)_2_ dimer probe, which are oriented perpendicular to one another, are alternately excited as the laser polarization direction is rotated across the corresponding EDTMs (see Fig. 3*C*). The modulated fluorescence from a single molecule is isolated through a pinhole and detected using an avalanche photodiode (APD), which is processed using the phase-tagged photon counting (PTPC) method (21). The signal modulation amplitude is called the ‘visibility,’ *v* = [|*μ* |^2^ ™ |*μ* |^2^]*/*[|*μ* |^2^ + |*μ* |^2^], where 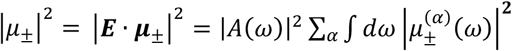 are the intensities associated with the symmetric and anti-symmetric excitons, **E** is the plane-polarized electric field vector of the laser, *A*(*ω*) is the (narrow) spectral envelope of the laser, and 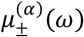 are the symmetric and anti-symmetric EDTMs of the *α*th vibronic transition (1).

As illustrated in Fig. 1*C*, the magnitudes of the **μ**_±_ EDTMs depend sensitively on the local conformation of the (iCy3)_2_ dimer. The orthogonal EDTMs define a ‘polarization ellipse’ in the cross-sectional area in which the laser beam projects onto the molecular frame, as shown in Fig. 3*C*. Therefore, the visibility, *v*, is directly related to the instantaneous local conformation of the (iCy3)_2_ dimer probe within the ss-dsDNA construct, which we monitor on microsecond and longer time scales.

### 2.3 Determination of probability distribution functions (PDFs) and two- and three-point time-correlation functions (TCFs) from PS-SMF experiments

In the experiments of Maurer *et al*. (1), the local fluctuations of individual (iCy3)_2_ dimer-labeled ss-dsDNA fork constructs (see Table 1 for base sequences) were monitored by recording time-dependent trajectories of the PS-SMF signal visibility, *v*(*t*). The average photon detection rate was typically ∼8,000 counts per second (cps), and the total duration of a single scan was *T* = ∼30 s. These time series of measurements were averaged over an adjustable integration (bin) period, *T*_*w*_, to construct ensemble-averaged probability distribution functions (PDFs), *P*(*v*), and the two-point and three-point time-correlation functions (TCFs), 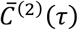 and 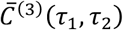,respectively (1,22-24). To compute these functions, we divided the continuous time trajectory into *N* discrete intervals, *T/N* = Δ*t* = *T*_*w*_, such that the *i*th time interval was assigned to the integer index, *i* = *t/*Δ*t*. We also defined the time delay, τ = *n*Δ*t*, with integer index *n*. In a typical procedure, we constructed the ensemble-averaged functions from ∼100 individual PS-SMF visibility trajectories. The resulting functions from each trajectory were in turn averaged together.

The normalized PDF was constructed from the sum of *N* discrete, time-averaged observations of equilibrium microstates with visibility, *v*.

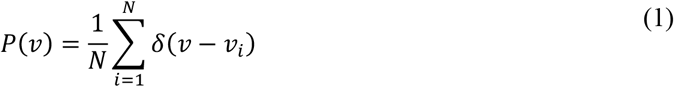

In Eq. (1), *δ*(*x*) is the Dirac delta function and *v*_*i*_ is the value of the visibility determined for the *i*th time interval. The normalized two-point TCF is the time-averaged product of two successive measurements separated by the interval τ

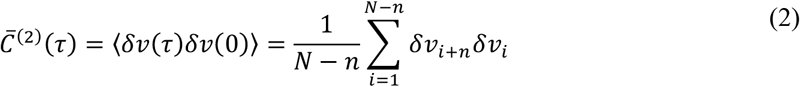

where *δv*(*t*) = *v*(*t*) ™ ⟨*v*⟩ [*δv*_*i*_ = *v*_*i*_ ™ ⟨*v*⟩] is the signal fluctuation about the mean value, ⟨*v*⟩. The two-point TCF,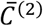, contains information about the number of quasi-stable macrostates of the system, the mean visibility of each macrostate, and the characteristic time scales of interconversion between macrostates (22). The normalized three-point TCF is the time-averaged product of three consecutive measurements separated by the intervals τ_1_ and τ_2_.

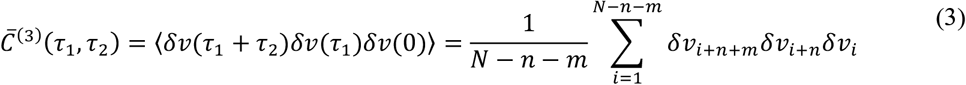

In Eq. (3), we have defined the time intervals τ_1_ = *n*Δ*t* and τ_2_ = *m*Δ*t*. The function 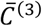 contains detailed kinetic information about the transition pathways that connect the macrostates of the system. This function is sensitive to the roles of intermediates, whose presence can facilitate or hinder successive transitions between macrostates (22).

As we discuss further below, the above functions can be simulated using a kinetic network model, which assumes that the system interconverts between a finite number of Boltzmann-weighted macrostates, and that the rates of interconversion between macrostates are controlled by the Arrhenius activation barriers. The minimum number of macrostates needed to fully describe PS-SMF data is equal to the number of decay components exhibited by the two-point TCF, plus one (22,24). By comparing experimental results to the predictions of the kinetic network model, it is possible to extract the relative stabilities and activation barriers between distinct macrostates, which may be used to parameterize the FESs associated with the local environment of DNA bases and sugar-phosphate backbones ‘sensed’ by the (iCy3)_2_ dimer probes.

In Fig. 4*A* is shown an example of a visibility trajectory, *v*(*t*), that was recorded for the +1 (iCy3)_2_ dimer-labeled ss-dsDNA fork construct and visualized using an integration period of *T*_*w*_ = 100 ms. This trajectory undergoes discontinuous transitions between a small number of discrete values within the range 0 < *v* < 0.4 (dashed lines are guides to the eye). The visibility is a direct measure of the conformational changes of the (iCy3)_2_ dimer probe. Control measurements of iCy3 monomer-labeled DNA constructs do not exhibit discontinuous transitions, in contrast to (iCy3)_2_ dimer-labeled ss-dsDNA constructs (1). From ∼100 individual PS-SMF measurements, each of ∼30 s in duration, the ensemble-averaged functions were constructed. In Fig. 4*B* is shown the PDF (using *T*_*w*_ = 100 ms), which exhibits multiple ‘low visibility’ and ‘high visibility’ states. The major component of ‘low visibility’ features of the PDF lie within the range 0 ≲ *v* ≲ 0.1, while a minor component of ‘high visibility’ features lie within the range 0.1 ≲ *v* ≲ 0.4. The relatively broad and asymmetrically-shaped PDF indicates the presence of both stable and thermally activated local conformational macrostates of the (iCy3)_2_ dimer probes.

**Figure 4.**
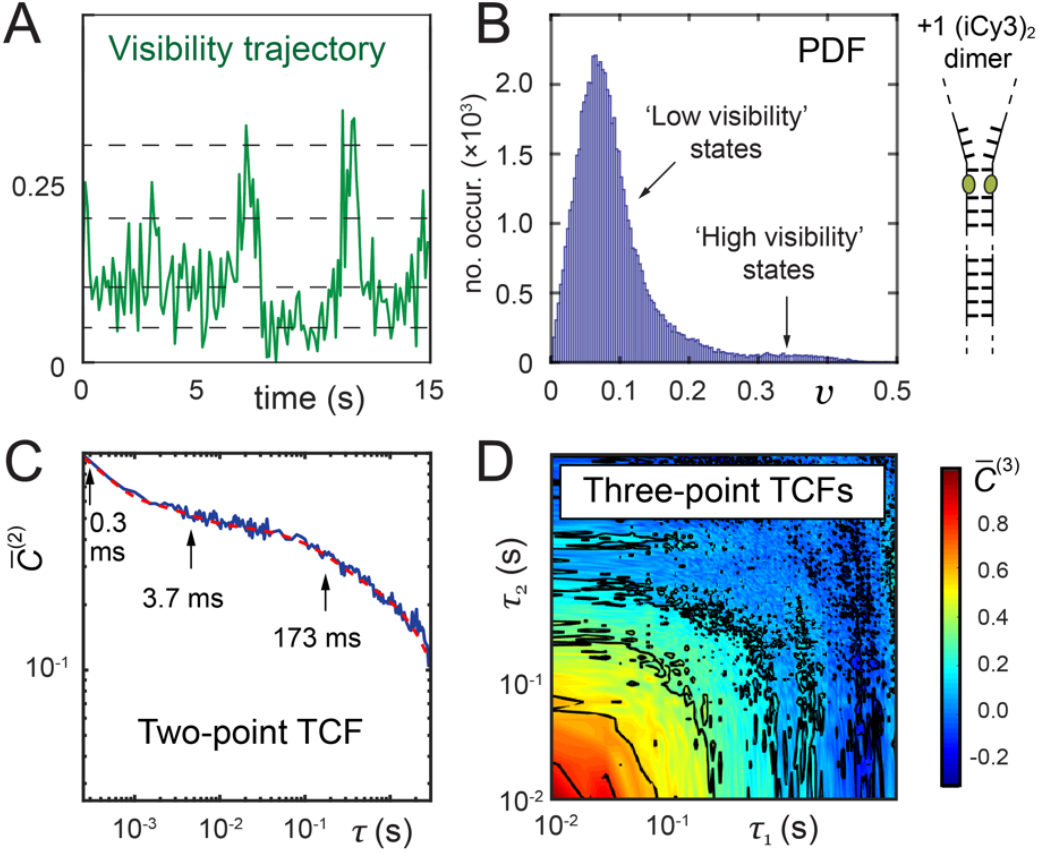
Sample experimental data from PS-SMF measurements of +1 (iCy3)_2_ dimer-labeled ss-dsDNA construct in 100 mM NaCl, 6 mM MgCl_2_, and 10 mM Tris at pH 8.0. (***A***) The first 15 seconds of a PS-SMF visibility trajectory, *v*(*t*), is shown using an integration period of *T*_*w*_ = 100 ms. The mean signal intensity was ∼7,500 s^-1^. Horizontal dashed lines are guides to the eye to indicate the presence of multiple discrete conformational states. (***B***) Normalized probability distribution function (PDF) of visibility values, *P*(*v*), using *T*_*w*_ = 100 ms. (***C***) Two-point time-correlation function (TCF, using *T*_*w*_ = 250 μs) is well modeled as a weighted sum of three exponentially decaying terms (red curve) with decay constants that are well-separated in time. (***D***) Three-point TCF (using *T*_*w*_ = 10 ms) is shown as a two-dimensional contour plot. Figure adapted from Ref. (25).

The dynamics of the +1 (iCy3)_2_ dimer-labeled ss-dsDNA fork construct are reflected by the properties of the two-point and three-point TCFs. In Fig. 4*C* is shown the two-point TCF (using *T*_*w*_ = 250 *μ*s), which is plotted alongside a fit to a model tri-exponential function. There are three well-separated decay components at *t*_1_ = 0.3 ms, *t*_2_ = 3.7 ms and *t*_3_ = 173 ms, which according to simple kinetic theory indicates that the underlying dynamics must involve at least four quasi-stable macrostates in dynamic equilibrium (22,26). In Fig. 4*D* is shown the corresponding three-point TCF (using *T*_*w*_ = 10 ms). The functional form of the three-point TCF is sensitive to the roles of pathway intermediates, whose presence can facilitate or hinder successive transitions between macrostates.

### 2.4 Inference of conformational macrostates of (iCy3)_2_ dimer-labeled ss-dsDNA constructs

As discussed in the previous sections, the ensemble and single-molecule studies of Heussman *et al*. (7-9) and Maurer *et al*. (1) as discussed in the previous sections, indicate that the local conformations of DNA bases and backbones immediately adjacent to the (iCy3)_2_ dimer probes fluctuate in thermal equilibrium between four quasi-stable macrostates. The results of these studies suggest that the FESs that govern the equilibrium distributions and interconversion dynamics are sensitive to position relative to the ss-dsDNA junction, as illustrated in Fig. 2. At positive integer positions within the duplex side of the ss-dsDNA junction the distribution of macrostates is dominated by a right-handed conformation, while at negative integer positions within the single-stranded side of the junction the distribution is dominated by a left-handed conformation. Based on the above observations and considering prior studies of DNA ‘breathing’ using hydrogen-tritium exchange methods (16-19), we may hypothesize structural assignments to the four conformational macrostates of the system.

Hydrogen-tritium exchange experiments performed on samples of reconstituted DNA (16-19) showed that a local ‘cooperatively exchanging’ region of duplex DNA (consisting of a ∼5 – 10 base-pair segment) can undergo thermally activated transitions to populate unstable ‘open’ conformations (4). In Fig. 5 are illustrated four conformational macrostates (labeled S_1_ – S_4_) in order of increasing visibility, which are consistent with the experimental observations of Heussman *et al*. (7-9) and Maurer *et al*. (1). The lowest visibility state, S_1_, represents the so-called ‘propped open’ (PO) macrostate, in which a hydrogen (H) bond within an affected base-pair is disrupted (*i*.*e*., held open) by the gain or loss of an inter-base hydrogen atom, resulting in two H-bond donors or two H-bond acceptors facing one another within a WC base pair. The ‘propped open’ state is right-handed and has stacked flanking bases. Thus, in the S_1_ macrostate, the complementary H-bonds of opposing bases are separated by relatively large distances (indicated by horizontal arrows), such that the twist angle *ϕ*_*AB*_ approaches 90° and the visibility approaches zero. S_2_ represents the canonical ‘Watson-Crick’ (WC) conformation, which is right-handed and has stacked flanking bases. The S_2_ macrostate has a right-handed twist angle *ϕ*_*AB*_ ≲ 90° (7). S_3_ represents a left-handed conformation in which flanking bases are unstacked (LHU) by relatively large distances (indicated by vertical arrows). The S_3_ macrostate has a left-handed twist angle *ϕ*_*AB*_ > 90°. The S_4_ macrostate (RHU), like S_3_, has its flanking bases unstacked, but here with a right-handed twist angle *ϕ*_*AB*_ < 90°. We note that for some of the experiments discussed below that were carried out at elevated and reduced salt concentrations it was necessary to include an additional fifth macrostate, S_5_, to obtain an essentially complete fit to the data.

**Figure 5.**
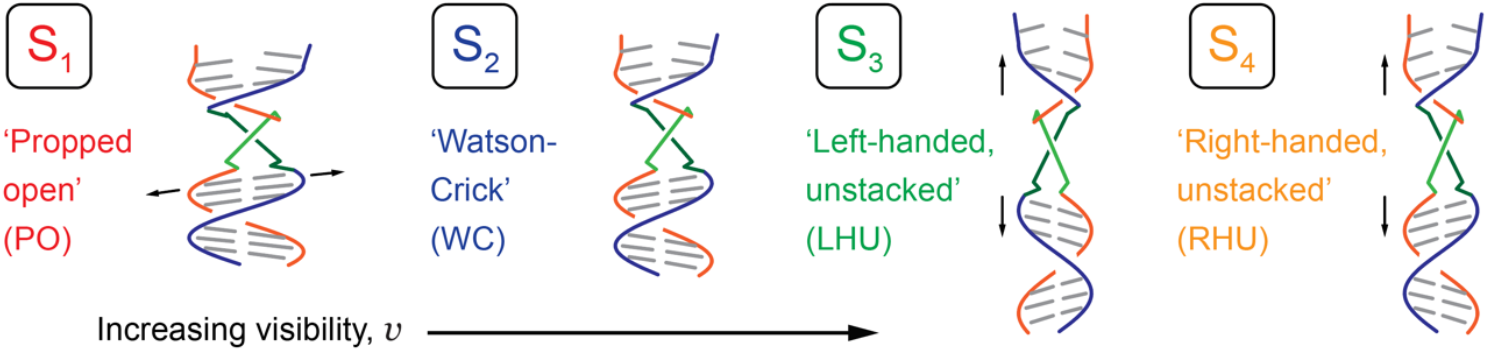
Hypothetical conformational macrostates of (iCy3)_2_ dimer-labeled ss-dsDNA constructs arranged in order of increasing visibility. The (iCy3)_2_ dimer is depicted as light green and dark green line segments attached to the sugar-phosphate backbones (shown in red and blue). The macrostates are color coded: S_1_ (PO, red), S_2_ (WC, blue), S_3_ (LHU, green), and S_4_ (RHU, orange). Adapted from Ref. (25).

### 2.5 Determination of FES parameters from kinetic network model analyses

The combined statistical-dynamical functions of the observed visibility shown in Fig. 4 [i.e., the PDF, *P*(*v*), the two-point TCF, 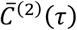,and the three-point TCF, 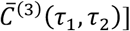,serve to constrain a multiparameter optimization of solutions to a four-state kinetic network model, which assumes that the four macrostates coexist at thermal equilibrium and interconvert stochastically with forward and backward rate constants, *k*_*ij*(*ij*)_. In the following, we briefly summarize this approach, which we have previously implemented in related studies (1,22-24).

The time-dependent population of the *i*th macrostate, *p*_*i*_ (*t*), is modeled using the master equation (26)

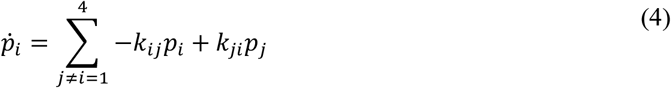

where the *k*_*ij(ij)*_ are the forward (backward) rate constants between macrostates *i* and *j* [*i, j* ∈ {1,2,3,4}]. A solution to Eq. (4) is obtained for an assumed set of input rate constants, which are subject to completeness and detailed balance conditions (22). The output parameters from the solution to Eq. (4) include (24): (*i*) the probabilities of observing the *i*th macrostate at equilibrium, 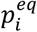;and (*ii*) the time-dependent conditional probabilities, *p*_*ij*_(τ), that the system undergoes a transition from macrostate *i* to macrostate *j* during the interval τ. We note that this includes the survival probabilities, *p*_*ii*_(τ), that the system initially observed in macrostate *i* remains in that macrostate during the interval τ.

Using the output parameters from a solution to Eq. (4), we model the PDF of the signal visibility as a sum of four macrostate contributions

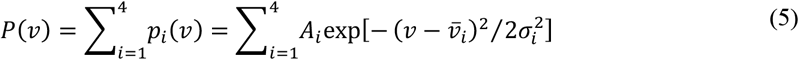

where we have assumed that the probability of the *i*th macrostate, *p*_*i*_(*v*), has a gaussian form with mean visibility 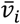,standard deviation σ_*i*_, and amplitude *A*_*i*_. The probability that the *i*th macrostate is present at equilibrium is the integrated area 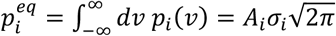.

We simulate the two-point TCF according to

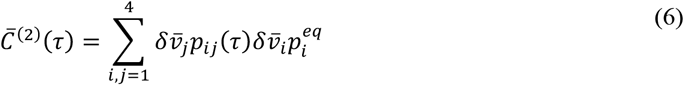

where 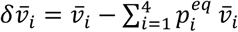 is the visibility fluctuation of the *i*th macrostate. According to Eq. (6),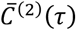 is a sum of 4 × 4 = 16 ‘kinetic pathways’ in which two consecutive fluctuations of the visibility occur, first with 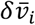 at τ = 0 and second with 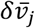 at time τ. Since the signs of the 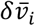 can be positive or negative, the various two-point products, 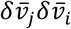 contribute both positive and negative amplitudes to the overall sum. The statistical weights of the various pathways depend on the products of probabilities, 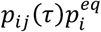,which depend on τ. Thus, the two-point TCF is the statistically-weighted product of two consecutive fluctuations, which occur within the interval τ as the system undergoes spontaneous transitions between the four macrostates.

We simulate the three-point TCF according to

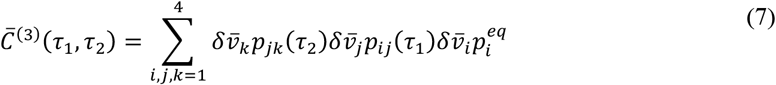

Equation (7) shows that the function 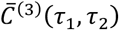 is a sum of (4)^3^ = 64 ‘transition pathways’ in which three consecutive fluctuations of the visibility occur – initially 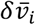,and then with 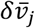 at time τ_1_ and 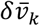 at time τ_2_. The various three-point products, 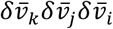 contribute both positive and negative amplitudes to the overall sum, with relative weights given by the product of probabilities, 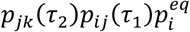 .We thus view the three-point TCF as the statistically-weighted product of three consecutive fluctuations separated by the intervals τ_1_ and τ_2_. The three-point TCF is sensitive to the roles of transient intermediates in the conformational transition pathways that connect the four macrostates at equilibrium.

We performed a multiparameter variational calculation to find ‘optimized’ solutions of Eq. (4) to simulate our PS-SMF data. For each of the (iCy3)_2_ dimer-labeled ss-dsDNA constructs discussed in the following sections, we calculated the statistical functions, 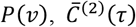 and 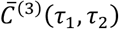, using Eqs. (5) – (7), respectively, for a given set of input rate constants, *k*_*ij*_, mean visibility values, 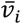,amplitudes, *A*_*i*_, and standard deviations, σ_*i*_. We quantified the agreement between calculated and experimentally-derived functions using a nonlinear least squares target function, χ^2^. We thus obtained ‘optimized’ solutions to the master equation by performing an iterative search of the parameter space to minimize the target function, following a procedure like that described by Phelps *et al*. (22,23) and Israels *et al*. (24). Additional details specific to these studies are provided in the SI. From these calculations, we determined optimized values of the forward and backward time constants, 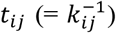,and the equilibrium populations, 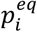.Using these parameters we determined the Boltzmann-weighted relative stabilities 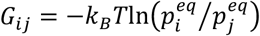, and the relative Arrhenius activation barriers, 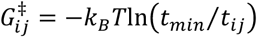,where *k*_*B*_ is Boltzmann’s constant and *t*_*min*_ is a reference time scale equal to that of the fastest process measured in the experiment (typically sub-millisecond).

## 3. Results and Discussion

### 3.1 Local conformational macrostates of the +1, -1 and -2 (iCy3)_2_ dimer-labeled ss-dsDNA fork constructs at ‘physiological’ salt concentrations

In Fig. 6 are shown comparisons between experimental and simulated PS-SMF data for the +1, -1, and -2 (iCy3)_2_ dimer-labeled ss-dsDNA constructs (Fig. 6*A* – 6*C*, respectively), which were obtained by implementing the four-state kinetic network analysis described in the previous sections. These data were obtained under ‘physiological’ salt conditions ([NaCl] = 100 mM, [MgCl_2_] = 6 mM). In all cases, the agreement between simulated and experimental functions is very good. The top row of Fig. 6 shows the PDFs overlaid with the four gaussian functions that represent the conformational macrostates, S_1_ – S_4_ (see Fig. 5 for hypothetical macrostate assignments). The normalized areas of the gaussians are equal to the equilibrium populations of the macrostates. In the second and third rows of Fig. 6, respectively, are shown the unnormalized two- and three-point TCFs, which characterize the conformational dynamics of the (iCy3)_2_ dimer-labeled ss-dsDNA constructs as a function of the probe labeling position relative to the ss-dsDNA fork junction. The two- and three-point TCFs are shown overlaid with their corresponding simulated functions. We emphasize that the final sets of optimized output parameters are obtained from our analyses by simultaneously fitting all three statistical-dynamical functions described by Eqs. (5) – (7).

**Figure 6.**
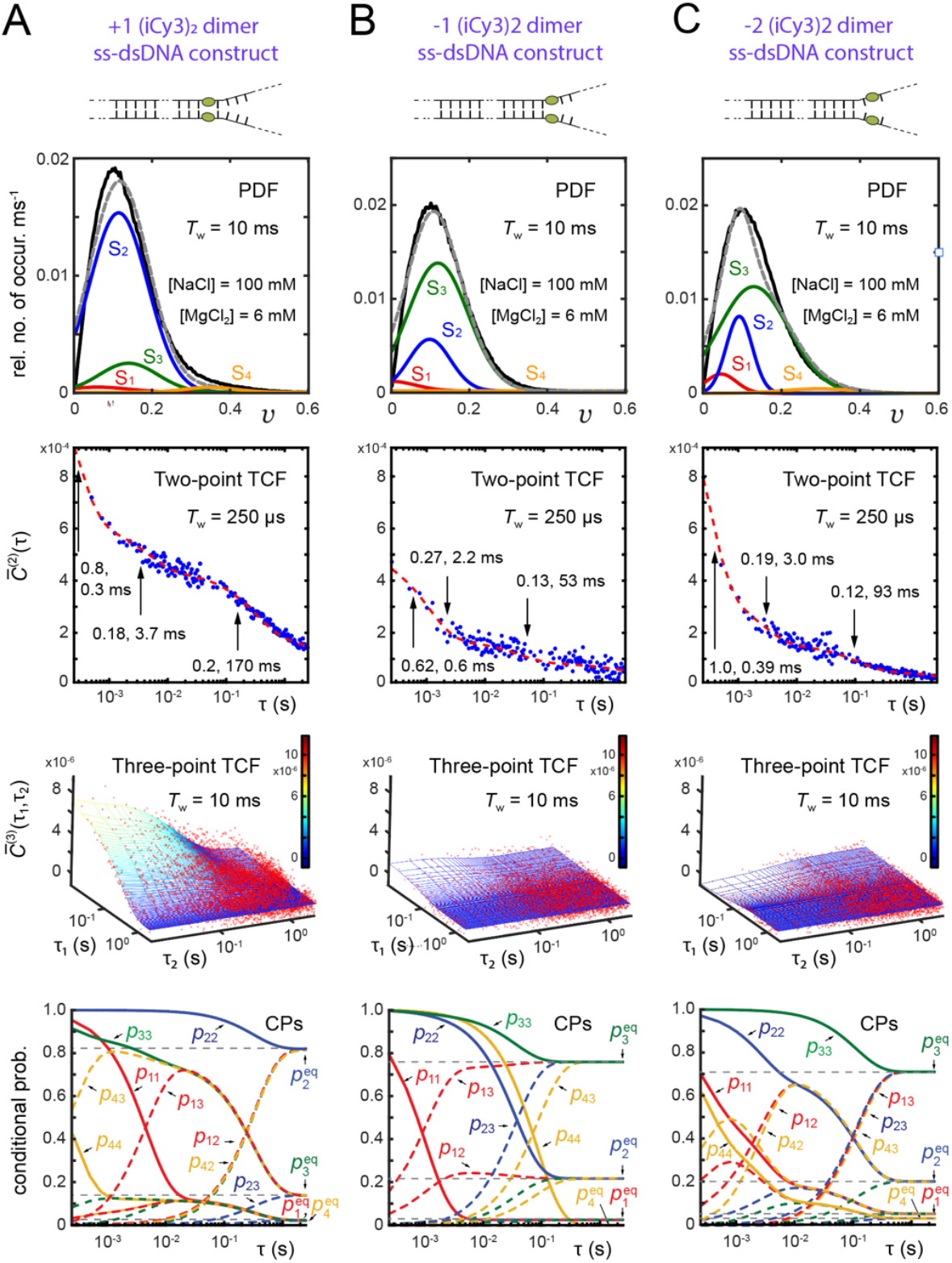
Results of four-state kinetic network analyses applied to PS-SMF measurements of the +1 (***A***), - 1 (***B***) and -2 (***C***) (iCy3)_2_ dimer-labeled ss-dsDNA fork constructs in 100 mM NaCl, 6 mM MgCl_2_, and 10 mM Tris at pH 8.0. Optimized kinetic and equilibrium parameters are obtained from simultaneously fitting the experimental PDF [*P*(*v*), black curves, first row], the two-point 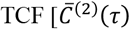 blue dots, second row], the three-point 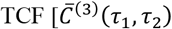,red dots, third row], and the conditional probabilities (CPs) [*p*_*ij*_(τ) with *i, j* ∈ {1,2,3,4}, fourth row]. Integration periods are indicated in the insets. The model PDFs (gray dashed curves) are represented as sums of four gaussian macrostates (red, blue, green and orange curves, labeled S_1_ – S_4_, respectively) with areas equal to the equilibrium probabilities, 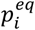.The model two-point TCFs are shown as red curves, and the model three-point TCFs are shown as solid surfaces. For the 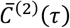 functions, the indicated amplitudes (*α*_*i*_) and time constants (*t*_*i*_) are derived from independent model fits of tri-exponential functions [∑_*i*_ *α*_*i*_*exp*(™ τ*/t*_*i*_)] to the experimental two-point TCFs, as described in (1). The CPs, *p*_*ij*_(τ), are shown as solid curves for *i* = *j* (called ‘survival probabilities’), and as dashed curves for *i* ≠ *j* (‘transition probabilities’). The CPs are color coded as the state labels, according to the value of *i* = 1: red, 2: blue, 3: green, and 4: orange. Figure partially adapted from Ref. (25).

In the fourth row of Fig. 6 are shown the time-dependent conditional probabilities (CPs), *p*_*ij*_(τ) with *i, j* ∈ {1,2,3,4}. The CPs are outputs of the kinetic network analysis, which characterize the kinetic behavior of the solutions used to generate the TCFs [Eqs. (6) and (7)]. The equilibrium populations are the asymptotic values approached by the CPs in the long-time limit. For example, the ‘survival probability’ of macrostate S_*i*_, *p*_*ii*_(τ) (solid curves), decays from unity to the equilibrium population value, 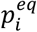 , on the time scale 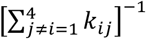,while the ‘transition probability,’ *p*_*ij*_(τ), increases from zero to 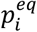 on the time scale 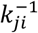.(22). For cases in which macrostate *i* is a transient intermediate species, the transition probability reaches a maximum value at an intermediate time before decaying to the equilibrium value 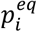.

The PDFs shown in Fig. 6 (top row) are each modeled as a sum of four Gaussian macrostates, S_1_ – S_4_. For each of the three (iCy3)_2_ dimer-labeled ss-dsDNA constructs, most of the population is partitioned between macrostates S_2_ (WC) and S_3_ (LHU), and the residual trace populations are in macrostates S_1_ (PO) and S_4_ (RHU). In Table S1 and Table S2 of the SI, we list the Gaussian parameters for these constructs at ‘physiological’ salt concentrations, as well as those for elevated and reduced salt concentrations discussed in the following sections. For the +1 construct, macrostate S_2_ (WC) is the majority component with 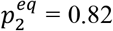 and mean visibility 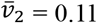 , and macrostate S_3_ (LHU) is the minority component with 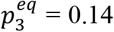 and mean visibility 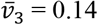. However, for the -1 and -2 constructs, the relative populations of macrostates S_2_ and S_3_ are juxtaposed in comparison to those of the +1 construct. For the -1 (-2) construct, macrostate S_3_ is the major component with 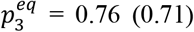 and mean visibility 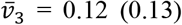,while macrostate S_2_ is the minor component with 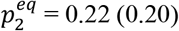 and mean visibility 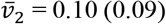.

The two-point and three-point TCFs shown in the second and third rows of Fig. 6, respectively, indicate that the conformational fluctuations of the (iCy3)_2_ dimer probes become abruptly faster as the probe position is varied across the ss-dsDNA junction, from the duplex towards the single-stranded region. The +1 construct exhibits the slowest overall decay with mean relaxation time 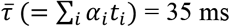,while the -1 and -2 constructs are significantly faster with mean relaxation times 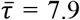 and 12.1 ms, respectively (1). This suggests that the activation barriers that mediate local conformational transitions of the (iCy3)_2_ dimer probes depend on the relative stabilities of the DNA bases and sugar-phosphate backbones immediately adjacent to the probes. The WC base pairs within the duplex side of the ss-dsDNA junction are largely stacked, while the stacking interactions between the bases on the ssDNA side of the junction are much weaker due to the lack of complementary base pairing and the resulting loss of stacking (and melting) cooperativity.

Information about the ‘pathways’ of conformational transitions at short times is contained within the initial amplitudes and signs of the two-point and three-point TCFs. For example, all the two-point TCFs shown in Fig. 6 exhibit positive initial amplitudes at the resolution of *T*_*w*_ = 250 μs. Our optimized solutions show that this follows because the positive pathway terms that contribute to the 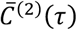 function [see Eq. (6)] are larger than the negative pathway terms.

In Table S3 of the SI, we list (in order of descending importance) the sixteen positive and negative pathway terms contributing to the 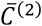 function at τ = 250 μs for each of the (iCy3)_2_ dimer-labeled ss-dsDNA constructs, including at varying salt concentrations studied in this work. We note that for the +1 construct at 300 mM NaCl, and for the -2 construct at 20 mM NaCl and 0 mM MgCl_2_, the inclusion of a fifth macrostate was necessary to obtain an optimized solution, S_5_, leading to additional pathway terms (see Table S1 and Table S2 of the SI). Each term in Table S3 is assigned to a specific two-point pathway (*i* → *j*) and factored into its two-point product of visibility fluctuations, 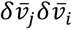,and its assigned statistical weight, 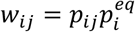.For all the (iCy3)_2_ dimer-labeled ss-dsDNA constructs, the 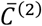 function is most heavily weighted by positive terms like 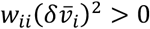, in which the initially measured macrostate, *i*, is measured again within the relatively short period, τ = 250 μs. This is because the survival probabilities, *p*_*ii*_, are generally large for small τ, as shown in the fourth row of Fig. 6 (solid curves).

Indeed, were the 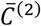 function to be negative, the dominant terms would involve consecutive two-point fluctuations of opposite sign (e.g., 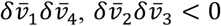etc.), in which transitions occur rapidly between macrostates on opposite sides of the visibility distribution separated by the mean visibility, 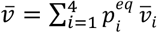. Such terms do not contribute significantly for small τ values, due to the relatively small magnitudes of the ‘transition probabilities,’ *p*_*ij*_ with *i* ≠ *j* (fourth row of Fig. 6, dashed curves). It is interesting to consider a scenario in which the transition probabilities would be large for small τ. In such a situation, the activation barriers mediating transitions between macrostates on opposite sides of 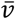 would be low in comparison to the thermal energy, *k*_*B*_*T*. Thus, lowering of the free energy barriers (or increasing temperature) leads to more closely-weighted terms of opposite sign, resulting in diminished amplitude of the 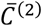 function.

‘Pathway’ considerations like those described for the two-point TCFs also apply to the three-point TCFs. For the three-point TCFs shown in the third row of Fig. 6, the +1 construct exhibits relatively large positive amplitude at the resolution of *T*_*w*_ = 10 ms, while the -1 construct exhibits weak positive amplitude and the -2 construct exhibits weak negative amplitude. In Table S4 of the SI, we list for each of the (iCy3)_2_ dimer-labeled ss-dsDNA constructs the eight largest positive and negative terms (of 64 total) that contribute to the 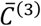 function for τ_1_ = τ_2_ = 10 ms [see Eq. (7)]. Each term is assigned to a specific three-point pathway (*i* → *j* → *k*) and factored into its three-point fluctuation product, 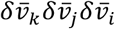,and statistical weight, 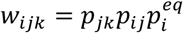.We emphasize that for all the dimer-labeled ss-dsDNA constructs, there are 64 positive and negative terms that contribute to the 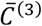 function. For cases in which half of the sampled macrostate fluctuations are positive and the other half are negative (see Table S2 of the SI), there are 32 positive terms contributing to 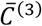,and 32 negative terms. If all the terms were equally weighted at the resolution τ_1_ = τ_2_ = 10 ms, the positive and negative terms would cancel exactly. It is the presence of relatively high activation barriers that leads to the prevalence of pathways in which the same macrostate is measured repeatedly, or in which two macrostates that are connected by a relatively low activation barrier are measured consecutively. As the activation barriers between distinct macrostates on opposite sides of the PDF are lowered, positive and negative terms become more similarly-weighted. Thus, the diminished magnitude of the 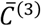 function that we observe as the dimer probe position is varied across the ss-dsDNA junction from the duplex towards the ssDNA regions is due partly to the decreased activation barriers between the four macrostates, in addition to the juxtaposition between the relative stabilities of macrostates S_2_ and S_3_.

For the +1 and -1 constructs at 100 mM NaCl and 6 mM MgCl_2_, the 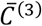 function is dominated by positive terms such as 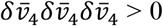 and 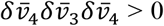 , for which consecutive three-point fluctuations of the visibility occur predominantly between macrostates on the positive side of the mean visibility,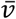 (see Table S4 of the SI). This is because pathway terms involving multiple factors of 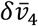 are relatively large, as are the CPs, *p*_43_ (for the +1 construct) and *p*_44_ (for the -1 construct), for τ_1_ = τ_2_ = 10 ms. Negative term contributions generally involve two measurements of macrostates on the positive side of the PDF and one on the negative side (e.g.,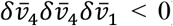),or three consecutive measurements on the negative side of the PDF (e.g.,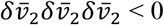). For the +1 construct, negative term contributions are small due to the relatively small magnitudes of the transition probabilities, *p*_*ij*_. However, for the -1 and -2 constructs, the magnitudes of the transition probabilities, *p*_*ij*_, increase for small τ. Thus, negative terms contribute more for the -1 and -2 constructs than for the +1 construct, and the resulting cancellation of terms diminishes the overall magnitudes of the 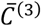 function.

We identify some notable trends in the behavior of the CPs and the three-point TCFs as the dimer probe position is varied across the ss-dsDNA junction. The lifetime of macrostate S_2_ (indicated by the decay time of *p*_22_) decreases while that of macrostate S_3_ increases, thus reflecting the juxtaposition of relative stabilities of macrostates S_2_ and S_3_ (as noted above) and a change in the activation barriers that mediate these states’ respective relaxations. For all the (iCy3)_2_ dimer-labeled ss-dsDNA constructs, the largest positive term contributing to the 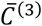 function at short time (τ_1_ = τ_2_ = 10 ms) contains the factor 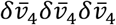 . However, the next-to-largest terms vary depending on the dimer probe position. For the +1 construct, the next-to-largest terms involve transitions between macrostates S_4_ ↔ S_3_, which include factors such as 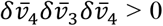. This is due to the relatively large transition probability, *p*_43_, for τ_1_ = τ_2_ = 10 ms. For the -1 construct, the next-to-largest terms contain the factors 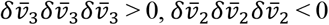 and 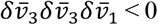, with the latter pathway involving transitions between macrostates S_3_ ↔ S_1_. The increased importance of transitions at short times between macrostates S_3_ ↔ S_1_ for the -1 construct is due to the increased population of macrostate S_3_ and to the increased transition probability, *p*_13_, for τ_1_ = τ_2_ = 10 ms. For the -2 construct, the next-to-largest terms involve transitions between macrostates S_4_ ↔ S_2_ and include factors such as 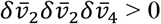 and 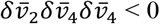 , which are relatively large due to the increased transition probability *p*_13_, for τ_1_= τ_2_ = 10 ms. The -2 construct has most positive and negative terms assigned very similar weights (with the negative terms slightly larger than the positive terms). The near cancellation between positive and negative terms for both the -1 and -2 constructs is due to the lowering of the activation barriers between disparate macrostates, as we next discuss.

In Fig. 7 we present the results of our kinetic network analysis of our PS-SMF experiments on the +1, -1, and -2 (iCy3)_2_ dimer-labeled ss-dsDNA constructs at ‘physiological’ salt concentrations. In the left column of Fig. 7 are shown the optimized kinetic network schemes for each of the three constructs. Macrostates S_1_ – S_4_ are shown interconnected by forward and backward time constants, 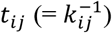,along with their equilibrium populations, 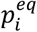.We note that the absence of an interconnection between macrostates indicates that such transitions are not observed in the experiments. For all the (iCy3)_2_ dimer-labeled ss-dsDNA constructs studied in this work, the optimized values for the equilibrium populations are listed in Table S2 of the SI and those for the time constants are listed in Table S5.

**Figure 7.**
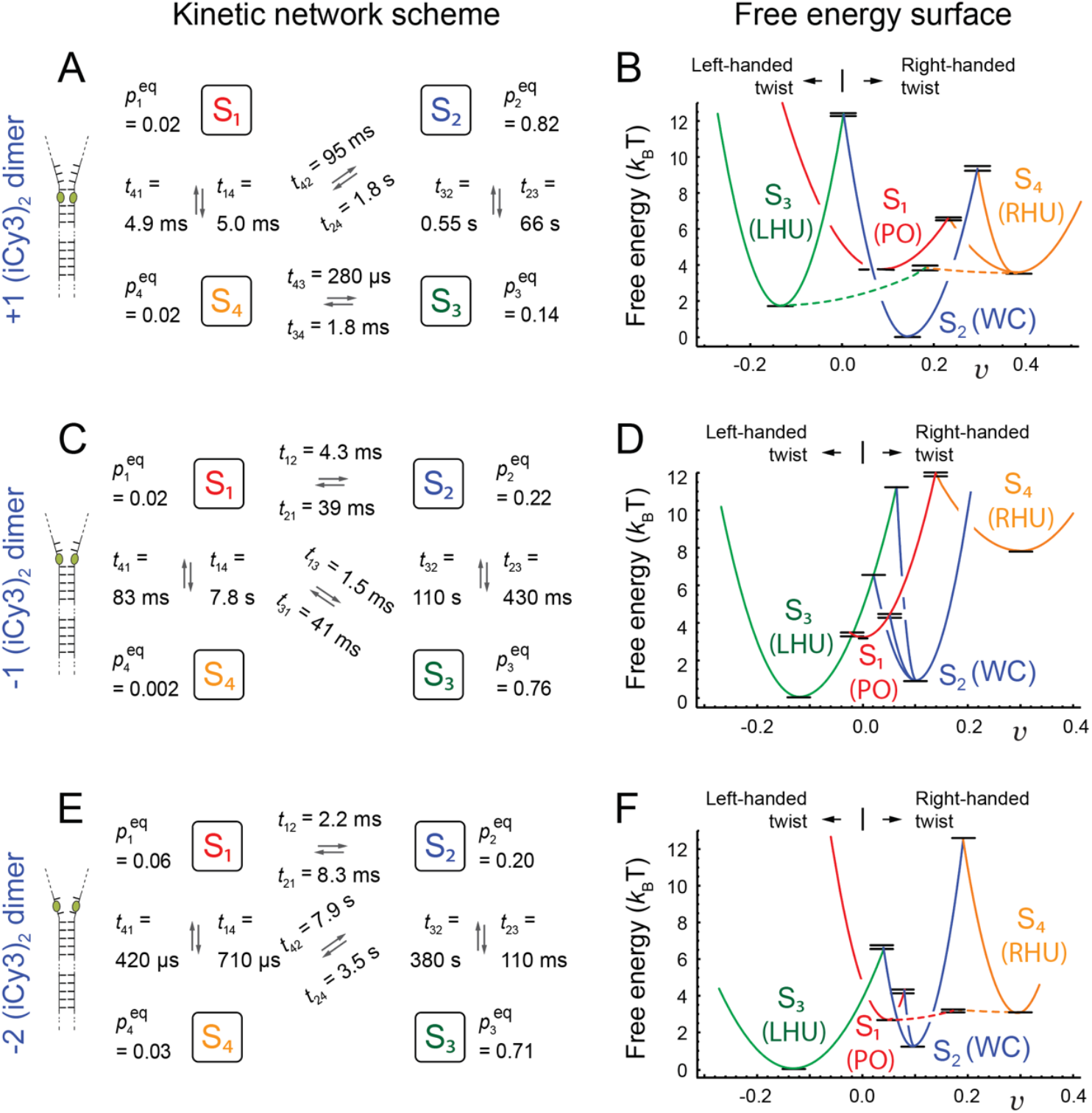
Kinetic network schemes and free energy surfaces (FESs) from the analyses of the (***A, B***) +1, (***C, D***) -1 and (***E, F***) -2 (iCy3)_2_ dimer-labeled ss-dsDNA fork constructs in aqueous buffer with 100 mM NaCl and 6 mM MgCl_2_. Panels ***A, C, E*** show the kinetic network schemes and equilibrium populations, 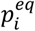 (listed in Table S2 of the SI), and time constants, 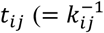 listed in Table S5), which were determined from the analysis of PS-SMF microscopy data. Tentative structural assignments to the four macrostates are illustrated in Fig. 5, with designations S_1_: ‘propped-open’ (PO), S_2_: ‘Watson-Crick’ (WC), S_3_: ‘left-handed unstacked’ (LHU) and S_4_: ‘right-handed unstacked’ (RHU). Panels ***B, D, F*** show the corresponding FESs inferred from the free energy minima (listed in Table S6) and activation barriers (listed in Table S8), which were calculated from the optimized equilibrium and kinetic parameters using the Boltzmann and Arrhenius equations, respectively. Adapted from Ref. (25).

From the equilibrium populations, we determined the relative free energy minima between pairs of macrostates by applying the Boltzmann relation, 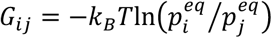.We designate the ‘ground state’ as the macrostate with the lowest free energy, *G*_*min*_ = 0, to which all ‘excited state’ energies are referenced. Similarly, we determined the relative activation barriers from the optimized time constants using the Arrhenius equation 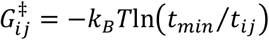,where *t*_*min*_ is the shortest optimized time constant (typically sub-millisecond) for a given sample. Finally, the activation energies shown in Fig. 7 (as black horizontal line segments) are vertically offset relative to their corresponding free energy minima according to 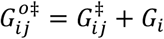.The values of the free energy minima and activation energies are listed in Table S6 and Table S7, respectively. The values of the activation energies obtained by applying the vertical offset, 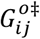,are listed in Table S8 of the SI.

In the right-hand column of Fig. 7 are shown the free energy surfaces (FESs) corresponding to the +1, -1 and -2 (iCy3)_2_ dimer-labeled ss-dsDNA constructs at ‘physiological’ salt concentrations, which we inferred from the free energy minima and activation barriers listed in Table S6 and Table S7, respectively. The FESs show how the relative stabilities and transition barriers of the four macrostates change as the (iCy3)_2_ dimer probe position is varied across the ss-dsDNA junction. We note that the PS-SMF experiment does not distinguish between positive and negative values of the visibility. We have therefore assigned a negative value to the visibility of macrostate S_3_, corresponding to its left-handed twist, and positive values to the visibilities of macrostates S_1_, S_2_ and S_4_, corresponding to their right-handed twists.

The local *thermodynamic* stability of a given (iCy3)_2_ dimer-labeled ss-dsDNA construct depends on the relative free energy minima of macrostates, S_1_ – S_4_. In Fig. 8*A*, we plot the values of the free energy minima as a function of dimer probe position (listed in Table S6). For the +1 construct, macrostate S_2_ (WC) exhibits the lowest free energy (ground state) and S_3_ (LHU) represents the first excited state conformation with energy 1.8 *k*_*B*_*T* (= ∼1.07 kcal mol^-1^). This relatively small difference in free energy may reflect a different stacking conformation of the G:C base pairs immediately adjacent to the (iCy3)_2_ dimer probes. Macrostates S_1_ (PO) and S_4_ (RHU) are degenerate second excited state conformations, each with free energies of 3.7 *k*_*B*_*T*. For the -1 and -2 constructs, the relative stabilities of macrostates S_2_ and S_3_ are reversed relative to that of the +1 construct, a result anticipated by the ensemble studies of Heussman *et al*. (7-9) (see discussion in Sect. 1). For the -1 and the -2 constructs, macrostate S_3_ is the ground state and S_2_ is the first excited state with free energy 1.2 *k*_*B*_*T* (1.3 *k*_*B*_*T*). For the -1 construct, the free energy of macrostate S_1_ (3.6 *k*_*B*_*T*) is very like that of the +1 construct. However, for the -1 construct the free energy of macrostate S_4_ is significantly increased (5.5 *k*_*B*_*T*) relative to that of the +1 construct. For the -2 construct, the free energies of macrostates S_1_ and S_4_ are reduced (2.5 *k*_*B*_*T* and 3.2 *k*_*B*_*T*, respectively) to values slightly lower than those of the +1 construct.

**Figure 8.**
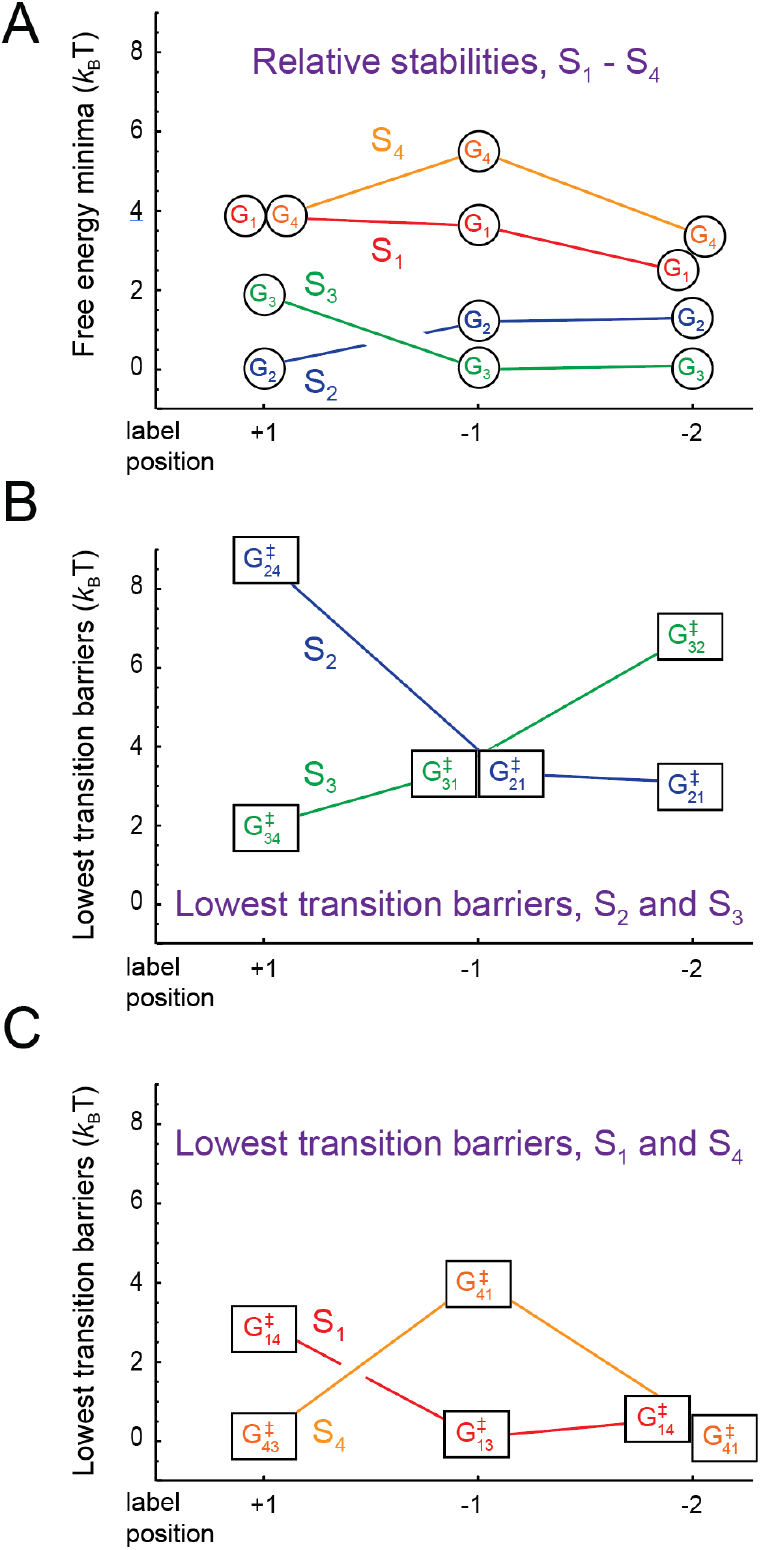
Position-dependent free energy parameters obtained from kinetic network analyses of the +1, -1 and -2 (iCy3)_2_ dimer-labeled ss-dsDNA constructs in aqueous buffer with 100 mM NaCl and 6 mM MgCl_2_. (***A***) Free energy minima of macrostates S_1_ – S_4_ (values listed in Table S6 and lowest lying transition barriers of (***B***) macrostates S_2_ and S_3_ and (***C***) macrostates S_1_ and S_4_ (values listed in Table S7).

The local *mechanical* stability of a given (iCy3)_2_ dimer-labeled ss-dsDNA construct depends on the magnitudes of transition barriers that mediate interconversion between the four macrostates, S_1_ – S_4_. For a given macrostate, the smallest transition barrier determines the primary relaxation pathway of that macrostate. In Fig. 8*B*, we plot the values of the lowest transition barriers for the two most stable of the four macrostates, S_2_ and S_3_, as a function of dimer probe position (see Table S7). For the +1 construct, the lowest transition barrier of macrostate S_2_ (WC) is significantly high (8.8 *k*_*B*_*T*), indicating that the ground state conformation for this construct has a relatively long lifetime 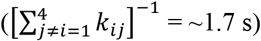.

In contrast, the lowest transition barrier of the first excited state conformation S_3_ (LHU) is relatively low (1.9 *k*_*B*_*T*) indicating that this macrostate has a moderate lifetime (∼81 ms). As the probe position is varied across the ss-dsDNA junction, from +1 to -1 to -2, the lowest transition barrier of macrostate S_2_ decreases (from 8.8 *k*_*B*_*T* to 3.3 *k*_*B*_*T* to 3.0 *k*_*B*_*T* with lifetimes 1.7 s to 36 ms to 7.7 ms, respectively) and that of macrostate S_3_ increases (from 1.9 *k*_*B*_*T* to 3.3 *k*_*B*_*T* to 6.8 *k*_*B*_*T* with lifetimes 1.8 ms to 41 ms to 380 ms). Thus, for the +1 construct, macrostate S_2_ is the thermodynamically favored conformation and the most mechanically stable. However, the S_2_ macrostate becomes thermodynamically and mechanically unstable (relative to S_3_) as the probe position is varied across the ss-dsDNA junction towards the ssDNA sequences. In Fig. 8*C*, we plot the position-dependent values of the lowest transition barriers for S_1_ (PO) and S_4_ (RHU), the least stable two of the four macrostates. For the +1 construct, the lowest transition barrier of macrostate S_4_ is immeasurably small (assigned to the reference value of zero) and that of S_1_ is low (2.9 *k*_*B*_*T*). As the probe position is varied across the ss-dsDNA junction, the transition barrier of the S_4_ macrostate initially increases and then decreases (from 0 to 4.0 *k*_*B*_*T* to 0) and that of macrostate S_1_ decreases (from 2.9 *k*_*B*_*T* to 0 to 0.5 *k*_*B*_*T*). Thus, both the thermodynamic and mechanical stabilities of macrostates S_1_, S_2_ and S_4_ decrease as the dimer probe position is moved across the ss-dsDNA junction towards the single-stranded sequences, while those of macrostate S_3_ increase, although not to the level of S_2_ at the +1 position.

The results summarized in Fig. 8 show why the local conformations adopted by the bases and sugar-phosphate backbones immediately adjacent to the (iCy3)_2_ dimer probes exhibit relative stabilities and transition barriers that depend sensitively on the position of each pair of probes within the ss-dsDNA fork junction. The conformational macrostates visible in the PS-SMF experiment, S_1_ – S_4_, likely reflect different local base stacking and backbone conformations within the duplex portion of the ss-dsDNA junction. Under ‘physiological’ buffer salt conditions, the local conformations of the +1, -1 and -2 (iCy3)_2_ dimer-labeled ss-dsDNA constructs exist primarily as Boltzmann-weighted mixtures of the right-handed macrostate S_2_ (WC) and the left-handed macrostate S_3_ (LHU). The S_2_ macrostate of the +1 construct is the thermodynamically favored ground state and exhibits the highest transition barrier of the three positions investigated, indicating that the thermodynamic ground state conformation within the duplex side of the fork junction, S_2_, is also mechanically stable with a relatively long population lifetime (∼300 ms).

In contrast, the left-handed S_3_ macrostate of the -2 construct is the thermodynamically favored conformation and exhibits a considerably smaller transition barrier, indicating that the ground state conformation within the ssDNA region of the fork junction, S_3_, is mechanically less stable, with a much shorter population lifetime (∼70 ms). The behavior of the -1 construct is somewhat unique and possibly reflects the complexity of the transition between the duplex and the ssDNA portions of the fork junction. While the S_3_ macrostate is thermodynamically favored relative to S_2_ at the -1 position, the transition barriers for both S_2_ and S_3_ are equal and slightly smaller than that of S_4_. Thus, the -1 position is the least mechanically stable of the three positions investigated. These observations are reminiscent of previous ensemble studies of local base stacking conformations at and near ss-dsDNA junctions using fluorescent base analogues, which suggested that the degree of base stacking is greatly diminished at the -1 construct position, relative to all the neighboring positions (27).

### 3.2 Salt concentration dependence of the local conformational macrostates of the +1 and -2 (iCy3)_2_ dimer-labeled ss-dsDNA fork constructs

More detailed information about the conformational macrostates discussed in the previous sections can be obtained by examining how the FES parameters of the (iCy3)_2_ dimer-labeled ss-dsDNA constructs depend on the salt concentrations of the solvent environment. For this purpose, we performed salt concentration-dependent studies of the +1 and -2 constructs to compare the different salt concentration-dependent behaviors of the duplex versus the single-stranded regions of the ss-dsDNA fork junction. We first consider how the equilibrium and dynamic properties of the (iCy3)_2_ dimer probes at positions near the ss-dsDNA fork junction can be influenced by the forces that govern DNA duplex stability. The results presented in Sect. 3.1 demonstrate that at ‘physiological’ salt concentrations (100 mM NaCl, 6 mM MgCl_2_), the +1 position within the duplex side of the junction favors the right-handed macrostate S_2_ (WC), and the -1 and -2 positions within the ssDNA side of the junction favor the left-handed unstacked macrostate S_3_ (LHU). The decline in stability of the S_2_ macrostate relative to the S_3_ macrostate at the -1 and -2 positions is likely related to the absence of complementary base-pairing at the negative integer positions and the resulting abrupt loss of cooperative base-stacking.

In duplex DNA the WC macrostate is the most stable, with adjacent base pairs optimally stacked (inter-base pair spacing ∼3.4 Å) and the sugar-phosphate backbone optimally extended. (28). However, thermal fluctuations can induce population of the energetically low-lying macrostates, including S_1_ (PO, in which the WC hydrogen bonds are disrupted while maintaining the fully stacked structure of flanking bases), S_3_ (LHU, in which the sugar-phosphate backbones adopt a left-handed twist and the flanking bases are somewhat unstacked) and S_4_ (RHU, in which the sugar-phosphate backbones adopt a right-handed twist, and flanking bases are fully unstacked). The marginal stability of the WC macrostate in duplex DNA can be understood in terms of opposing thermodynamic forces that influence base stacking (11,29-34). Forces that favor base stacking are primarily entropic – i.e., dependent on the expulsion of water molecules that are otherwise confined between flat base surfaces – and thus have some similarity to the hydrophobic bonding interactions that drive protein folding (35-38). The forces that oppose base stacking are primarily enthalpic – i.e., the electrostatic repulsion between negatively charged backbone phosphate groups, which at physiological salt concentrations are approximately 70% screened on average by positive ions within the condensation layer immediately surrounding the negatively charged DNA-water interface (14).

We note that the hydrogen-tritium exchange experiments of McConnell *et al*. and Printz *et al*. (16-19) showed that the exchange rate of complementary WC hydrogens is minimized at ‘physiological’ salt concentrations but rises precipitously when the salt concentration is either elevated or reduced. Decreasing salt concentrations from physiological levels is thought to destabilize the DNA duplex by reducing the electrostatic screening between negatively charged phosphate groups (15). However, increasing salt concentrations to moderate levels above physiological (>100 mM NaCl) can also destabilize the DNA duplex, although the molecular origin of this destabilization is not well understood. A possible explanation for duplex destabilization at moderate salt concentrations is a reduction of water entropy associated with the formation of structured solvation shells around the increasing concentrations of ions and counterions in bulk solution. Nevertheless, the specific roles played by monovalent sodium and divalent magnesium ions in stabilizing (or destabilizing) conformational macrostates are unclear. The following salt-dependent studies examine how sodium and magnesium cation concentrations affect the equilibrium distributions and transition barriers of the conformational macrostates at the +1 and -2 positions relative to the ss-dsDNA fork junction.

Our salt-concentration-dependent results for the +1 (iCy3)_2_ dimer-labeled ss-dsDNA construct are presented in Fig. S1 of the SI. The experimental functions are quite sensitive to variations in salt concentrations, thus demonstrating the sensitivity of our PS-SMF measurements to changes in nucleic acid stability and dynamics. For all the salt concentrations that we studied, the PDFs (top row) exhibit a pattern of low- and high-visibility features like those discussed in the previous section, and most of the two-point and three-point TCFs (second and third rows, respectively) exhibit three well-separated decay components (well modeled using four macrostates). An exception is the TCF of the highest salt concentration (Fig. S1*A*), which exhibits four well-separated decay components (well modeled using five macrostates) (1). The experimental data are shown overlaid with the simulated functions based on the kinetic network model analyses described in previous sections. In all cases, the agreement between simulated and experimental functions is very good. The Gaussian parameters for all constructs and salt concentrations (including equilibrium probabilities) are listed in Table S1 and Table S2 of the SI.

In Fig. S2 of the SI, we present the optimized kinetic network schemes for each set of salt concentrations (left column), and the associated free energy surfaces (FESs, right column). The values of the free energy minima and activation energies are listed in Table S6 and Table S7, respectively.

In Fig. 9, we summarize our salt concentration-dependent measurements of the FES parameters of the +1 (iCy3)_2_ dimer-labeled ss-dsDNA construct. As discussed in previous sections, the local thermodynamic stability of the +1 construct depends on the relative free energy minima of the underlying macrostates. We first focus on how the relative stabilities of the macrostates depend on monovalent sodium and divalent magnesium ion concentrations. In Figs. 9*A* and 9*B* are shown the free energy minima of the macrostates plotted – from left to right – as a function of decreasing sodium ion concentration while holding the magnesium ion concentration fixed ([MgCl_2_] = 6 mM). We note that this is equivalent to increasing the ratio: [MgCl_2_]/[NaCl]. At the highest salt concentrations (300 mM NaCl, 6 mM MgCl_2_), macrostate S_3_ (LHU) is the ground state, which is significantly more stable than the first excited state S_2_ (WC, Fig. 9*A*).

**Figure 9.**
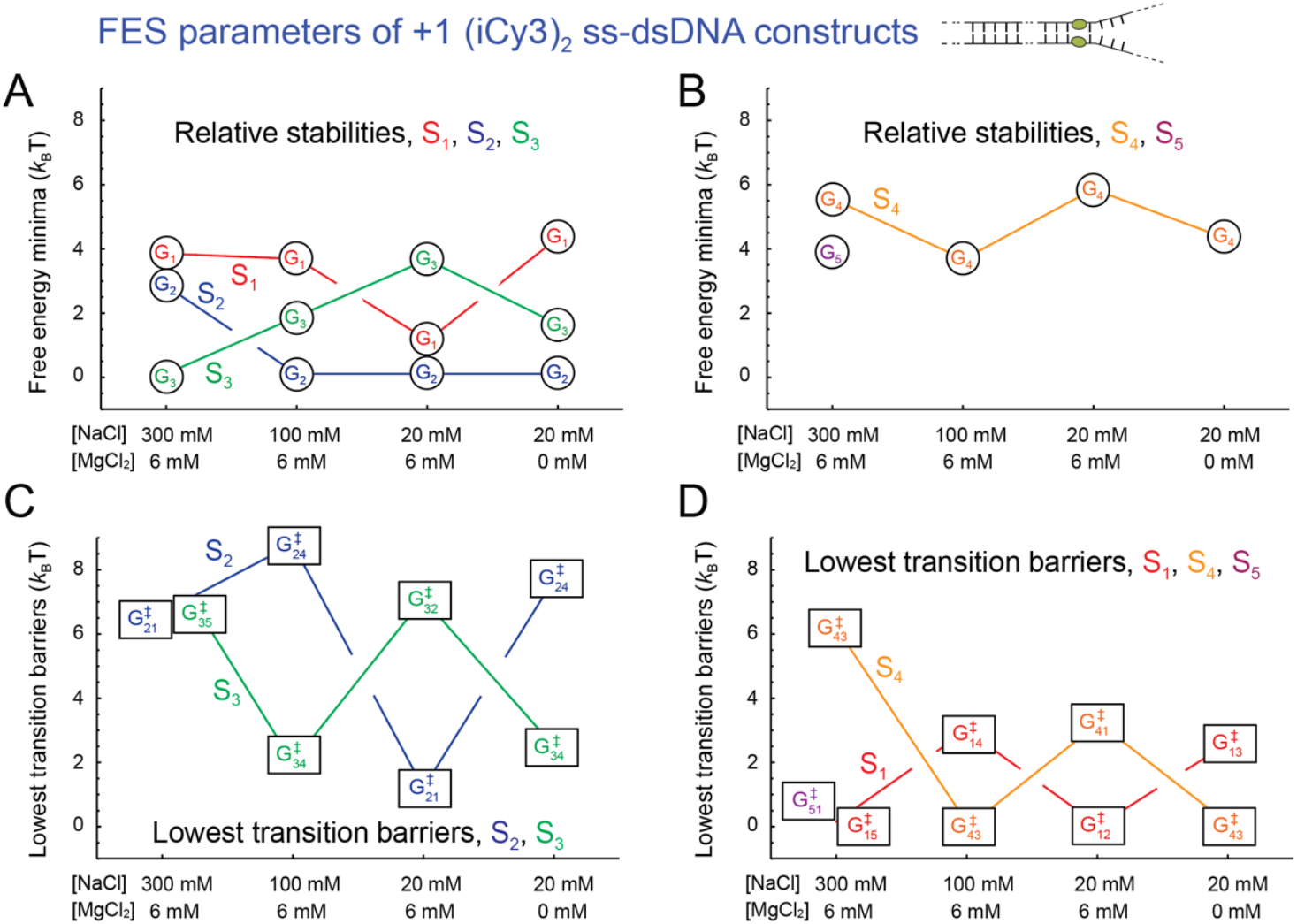
Salt concentration-dependent free energy parameters obtained from kinetic network analyses of the +1 (iCy3)_2_ dimer-labeled ss-dsDNA constructs in aqueous buffer and varying salt concentrations. Formatting and color schemes are the same as described in Fig. 8. Values for the free energy minima are listed in Table S6 of the SI and those for the transition barriers are listed in Table S7.

For all the other salt concentrations that we studied (including ‘physiological’: 100 mM NaCl, 6 mM MgCl_2_), the S_2_ macrostate is the ground state. As the sodium ion concentration is decreased from 300 mM to 100 mM, and then from 100 mM to 20 mM, the S_3_ macrostate becomes progressively less stable. Over the same range of decreasing sodium ion concentration (or increasing [MgCl_2_]/[NaCl] ratio) the S_1_ macrostate becomes progressively more stable. Elimination of magnesium at low sodium ion concentration (equivalent to [MgCl_2_]/[NaCl] = 0, far right column) destabilizes the S_1_ macrostate and stabilizes the S_3_ macrostate, resulting in a situation resembling the physiological condition. In contrast, the stability of the S_4_ (RHU) macrostate does not exhibit a consistent dependence on salt ion concentrations. At the highest salt concentrations (300 mM NaCl, 6 mM MgCl_2_), the S_1_ and S_5_ macrostates exhibit multiple (degenerate) free energy minima. The optimized kinetic scheme (see Fig. S2*A*) shows that the S_5_ macrostate is directly connected to S_1_ and S_3_.

The local mechanical stability of the +1 construct depends on the magnitudes of the transition barriers of the various macrostates, and the smallest transition barrier for a given macrostate determines its primary relaxation pathway. In Figs. 9*C* and 9*D*, we plot as a function of salt concentrations the lowest transition barriers of macrostates S_1_ – S_5_ for the +1 (iCy3)_2_ dimer-labeled ss-dsDNA construct. As discussed in previous sections, at ‘physiological’ salt concentrations (100 mM NaCl, 6 mM MgCl_2_) the S_1_ and S_2_ macrostates – for which the flanking bases are stacked – exhibit relatively high transition barriers, and the S_3_ and S_4_ macrostates – for which the flanking bases are unstacked – exhibit relatively low transition barriers. When the monovalent sodium ion concentration is elevated or reduced at fixed magnesium ion concentration (6 mM) relative to ‘physiological’ concentrations, the transition barriers for the base-stacked macrostates (S_1_ and S_2_) are decreased, while those for the base-unstacked macrostates (S_3_ and S_4_) are increased. Elimination of divalent magnesium at low sodium ion concentration (20 mM) restores the transition barrier values close to those observed at physiological salt concentrations. It is interesting that at the highest salt concentrations (300 mM NaCl, 6 mM MgCl_2_), the transition barrier of the S_5_ macrostate is nearly equal to that of macrostate S_1_.

We next consider our salt concentration-dependent results for the -2 (iCy3)_2_ dimer-labeled ss-dsDNA construct, which we present in Fig. S3 of the SI. For all the salt concentrations that we studied, the PDFs (top row) exhibit a pattern of low- and high-visibility features like those discussed in previous sections. For most of the salt concentrations, the two-point and three-point TCFs (second and third rows, respectively) exhibit three well-separated decay components. However, for the lowest salt concentrations (20 mM NaCl and 0 mM MgCl_2_, Fig. S3*D*), the TCFs exhibit four well-separated decay components (1). The experimental data are shown overlaid with the simulated functions based on the kinetic network model analyses. In all cases, the agreement between simulated and experimental functions is very good. The corresponding kinetic network schemes for each set of salt concentrations and the associated FESs are presented in Fig. S4 of the SI.

In Fig. 10, we summarize our salt concentration-dependent measurements of the FES parameters for the -2 (iCy3)_2_ dimer-labeled ss-dsDNA constructs. In contrast to the +1 construct, in which the dimer probe is positioned within the double-stranded portion of the ss-dsDNA fork junction, the probes in the -2 ss-dsDNA construct are in the single-stranded arms of the fork junction. The salt-dependent changes for the -2 position shown in Fig. 10 permit comparison with those for the +1 position shown in Fig. 9. In Figs. 10*A* and 10*B* we show the free energy minima for the -2-construct plotted – from left to right – as a function of decreasing sodium ion concentration while holding the magnesium ion concentration fixed ([MgCl_2_] = 6 mM). At physiological salt concentrations (100 mM NaCl, 6 mM MgCl_2_), the most stable (ground state) macrostate at the -2 position is S_3_ (LHU) and the next most stable macrostate is S_2_ (WC). This contrasts with the situation seen for +1 position, in which the relative stabilities of macrostates S_2_ and S_3_ are reversed (see Fig. 9). When the monovalent sodium ion concentration is elevated or reduced at fixed magnesium ion concentration relative to ‘physiological’ concentrations, the S_3_ and S_4_ (RHU) macrostates at the -2 position are destabilized while the S_2_ macrostate is stabilized. In contrast, as the sodium ion concentration is decreased from its highest to its lowest value at fixed magnesium ion concentration, the S_1_ macrostate is destabilized. These salt-dependent variations in stability at the -2 position are opposite (or nearly opposite) to those we observe at the +1 position. When the concentration of divalent magnesium ions is reduced to zero at low sodium ion concentration (20 mM NaCl, 0 MgCl_2_, far right column), the relative stabilities of the macrostates undergo a significant change. In this low sodium ion concentration in the absence of magnesium, macrostates S_2_, S_1_, S_3_ and S_4_ are all stabilized and populated at significant levels. An additional unstable macrostate S_5_ is also present, which connects directly to macrostate S_1_ (see Fig. S4*G*).

**Figure 10.**
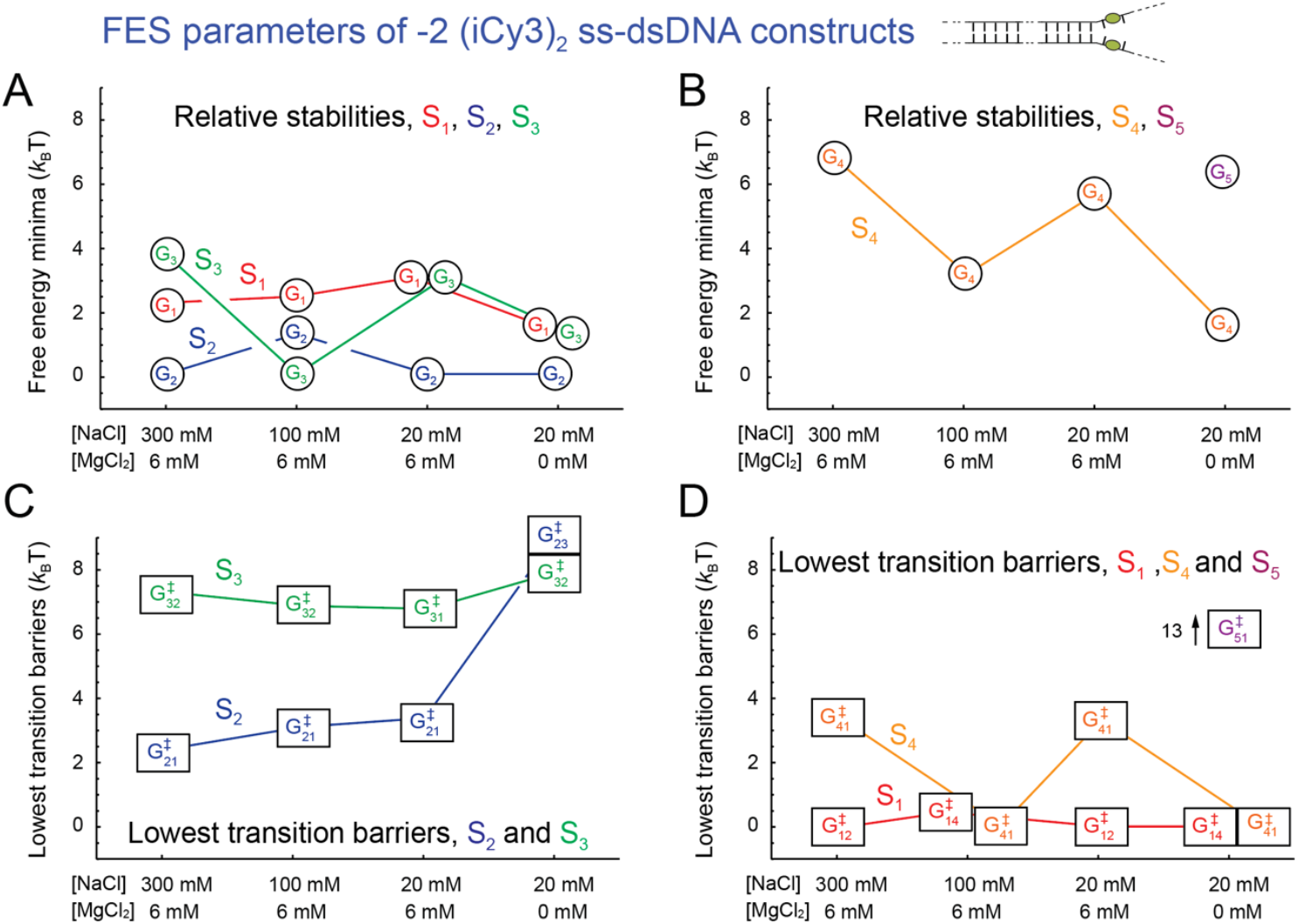
Salt concentration-dependent free energy parameters obtained from kinetic network analyses of the -2 (iCy3)_2_ dimer-labeled ss-dsDNA constructs in aqueous buffer and varying salt concentrations. Formatting and color schemes are the same as described in Fig. 8. Values for the free energy minima are listed in Table S6 and those for the transition barriers are listed in Table S7.

In Figs. 10*C* and 10*D*, we plot the lowest transition barriers of macrostates S_1_ – S_5_ for the -2 (iCy3)_2_ dimer-labeled ss-dsDNA construct as a function of salt concentration. We find that the transition barriers at the -2 position are much less sensitive to changes in salt concentration than are the barriers at the +1 position, as anticipated from the relatively fast decays of the two-point and three-point TCFs (see Fig. S3 of the SI). In addition, when the monovalent sodium ion concentration is decreased at constant magnesium ion concentration, the transition barriers of macrostates S_2_ and S_3_ change very little. In contrast, the transition barrier of macrostate S_4_ is immeasurably small at ‘physiological’ salt concentrations and increases to levels comparable to those seen for the S_2_ macrostate when the sodium ion concentration is elevated or reduced from ‘physiological’ values. Eliminating magnesium ions at low sodium ion concentration (20 mM NaCl, 0 MgCl_2_) results in significant increases in the transition barriers of macrostates S_2_ and S_3_ (9.1 *k*_*B*_*T* and 7.8 *k*_*B*_*T*, respectively), and macrostate S_5_ exhibits a uniquely high transition barrier (13.2 *k*_*B*_*T*). The high transition barriers observed in the absence of divalent magnesium ion concentration give rise to the relatively large positive initial amplitude of the three-point TCF at resolution *T*_*w*_ = 10 ms.

## 4. Conclusions and Overview

### 4.1. Kinetic network model analysis of PS-SMF experiments on (iCy3)_2_ dimer-labeled ss-dsDNA fork constructs

In this work, we have applied a kinetic network model analysis to interpret – in structural and dynamical terms – PS-SMF measurements of the local conformational fluctuations of exciton-coupled (iCy3)_2_ dimer-labeled ss-dsDNA fork constructs. The PS-SMF method directly monitors the structural fluctuations of (iCy3)_2_ dimer probes that are site-specifically positioned relative to the ss-dsDNA fork junction and report on the local conformational changes of the sugar-phosphate backbones and bases immediately adjacent to the probes (1). This approach provides detailed information about the relative stabilities and transition barriers associated with the conformational macrostates that characterize the local free energy surfaces (FESs). Our results corroborate previous ensemble spectroscopic studies (6,8) that had determined that the *average* local conformation of the probes exhibits a right-handed twist at the +1 position and a left-handed twist at the -1 and -2 positions (see Fig. 2). Moreover, prior studies of structural disorder in these systems (9) concluded that only a small number of macrostates coexist at equilibrium, that a right-handed macrostate is favored at positive integer positions and that a left-handed macrostate is favored at negative integer positions.

### 4.2. Position-dependent macrostate distributions and free energy surfaces (FESs) of (iCy3)_2_ dimer-labeled ss-dsDNA fork constructs at ‘physiological’ salt concentrations

We find that at ‘physiological’ salt conditions the local conformation at and near the ss-dsDNA fork junction interconverts primarily between four distinct macrostates (S_1_ – S_4_, as illustrated in Fig. 5), each corresponding to a defined local configuration of bases, base-pairs and sugar-phosphate backbones. For all the construct positions that we investigated, the ‘right-handed Watson-Crick’ S_2_ (WC) macrostate and the ‘left-handed unstacked’ S_3_ (LHU) macrostate were the major DNA components present, while the ‘propped open’ S_1_ (PO) and ‘right-handed unstacked’ S_4_ (RHU) macrostates were populated only at trace levels (see Fig. 8). At the +1 position (within the duplex portion of the fork junction), the S_2_ (WC) macrostate is thermodynamically favored and mechanically stable, with a population lifetime ∼1.7 s. However, at the -1 and -2 positions (within the single-stranded portions of the junction), the S_3_ (PO) macrostate is thermodynamically favored and less mechanically stable than it is when present within the duplex positions, with population lifetimes of ∼41 ms (-1) and ∼380 ms (-2), respectively.

### 4.3. Effects of salt concentration changes on macrostate stabilities and transition barriers

More detailed information about the nature of the macrostates near fork junctions is revealed by our salt concentration-dependent studies of the +1 and -2 (iCy3)_2_ dimer-labeled ss-dsDNA constructs. For the +1 construct, the S_2_ (WC) macrostate is the thermodynamically stable ground state at ‘physiological’ (100 mM NaCl, 6 mM MgCl_2_) and reduced (20 mM NaCl, 6 mM MgCl_2_ and 20 mM NaCl, 0 mM MgCl_2_) salt concentrations (see Fig. 9*A*). However, at elevated salt concentrations (300 mM NaCl, 6 mM MgCl_2_), the S_3_ (LHU) macrostate is most stable. This may be due to a reduction of favorable base-stacking interactions at moderately high sodium ion concentrations (> 100 mM), which may coincide with the formation of structured ion solvation shells within the aqueous environment of the dsDNA portions of this construct. Another important factor is the screening of the negatively charged backbone phosphates by the monovalent sodium ions within the ion condensation layer surrounding the DNA cylinders. As noted above, we also observed that the stability of the S_3_ (LHU) macrostate scales with increasing sodium ion concentration at constant magnesium ion concentration (6 mM MgCl_2_). A possible explanation for this behavior is that the repulsive interactions between the backbone phosphates in the S_3_ macrostate are more effectively screened by the monovalent sodium ions within the condensation layer, so that the S_3_ macrostate is preferentially stabilized relative to the other macrostates at the elevated sodium ion concentration.

To further explore this possibility, we note that the left-handed twist of the S_3_ macrostate, which is local to the ss-dsDNA fork junction, is reminiscent of the left-handed helical turn found in segments of double-stranded Z-DNA (28,39). Z-DNA exhibits a secondary structure pattern of nucleotide residues in which purine bases adopt *syn* rather than *anti* conformations within this dsDNA form. Conversion from the *anti* to the *syn* form of the purine bases occurs by rotating the purine residue about its glycosidic bond, which is accompanied by the transformation of the purine sugar pucker from C2’ endo to C3’ endo. The complementary nucleotide residue within the base pair involved has its pyrimidine base and sugar moieties rotated relative to the helical axis to maintain the canonical WC H-bonding structure. For comparison, in B-DNA all base orientations are *anti*, and all sugar pucker conformations are C2’ endo. In Z-DNA, the degree of base stacking is diminished relative to B-DNA (the rise per DNA base-pair along the dsDNA axis is 3.7 Å versus 3.4 Å) and the inter-base H bonds are displaced radially from the longitudinal symmetry axis (28,39). We note that consequently Z-DNA is also more stabilized by elevated monovalent ion concentrations than B-DNA, due to the smaller intra-strand phosphate-phosphate distance of the C3’ endo (5.9 Å) compared to the C2’ endo (7.0 Å) sugar pucker forms (28,39-41), which results in a higher negative charge density and thus more ion condensation in the Z-form than in the B- (or WC) form of duplex DNA

We note also that the stability of the S_1_ (PO) macrostate scales with the increasing ratio of divalent magnesium to monovalent sodium ion ([MgCl_2_]/[NaCl]) in the solvent environment (Fig. 9*A*). When the sodium ion concentration in solution is reduced while the magnesium ion concentration is held constant, the divalent magnesium plays a more prominent role in the interactions between positive solution ions and negatively charged backbone phosphates. In contrast to (non-site-specifically) condensed monovalent sodium ions, divalent magnesium binds strongly (and specifically) to individual backbone phosphates, and thus a single magnesium cation can, in effect, reverse the effective charge of individual backbone phosphates from negative to positive (42,43).

As discussed in the previous section, the S_1_ (PO) macrostate is most effectively stabilized in the +1 construct at the same reduced salt concentrations for which the S_3_ macrostate is least stable (20 mM NaCl, 6 mM MgCl_2_, Fig. 9*A*). However, elimination of magnesium at this low sodium ion concentration (20 mM NaCl, 0 mM MgCl_2_) reverses the relative stabilities of macrostates S_1_ and S_3_, resulting in a distribution of macrostate stabilities resembling that observed at ‘physiological’ salt concentrations. A possible explanation for this behavior is that the S_1_ macrostate is preferentially stabilized by divalent magnesium ions, but not by monovalent sodium ions. This contrasts with the behavior of the S_3_ macrostate, which is preferentially stabilized by monovalent sodium ions, but not by divalent magnesium ions.

### 4.4. Proposed structural and dynamical models for DNA macrostates positioned within the single-stranded and the double-stranded regions of the DNA fork junction

In Fig. 11*A*, we illustrate our proposed structural assignments for the macrostates of the +1 (iCy3)_2_ dimer-labeled ss-dsDNA construct. We assume that the possible local conformations of the (iCy3)_2_ dimer probe, which each gives rise to a unique value of the PS-SMF signal visibility, *v*, are directly related to the local conformations of the bases and backbones immediately adjacent to the probes. Each macrostate is thus depicted with complementary base-pair and sugar residues nearest to the probes shown atomistically. As we discuss further below, these assignments provide additional secondary structure details that are suggested by the salt concentration-dependences of the relative macrostate stabilities.

**Figure 11.**
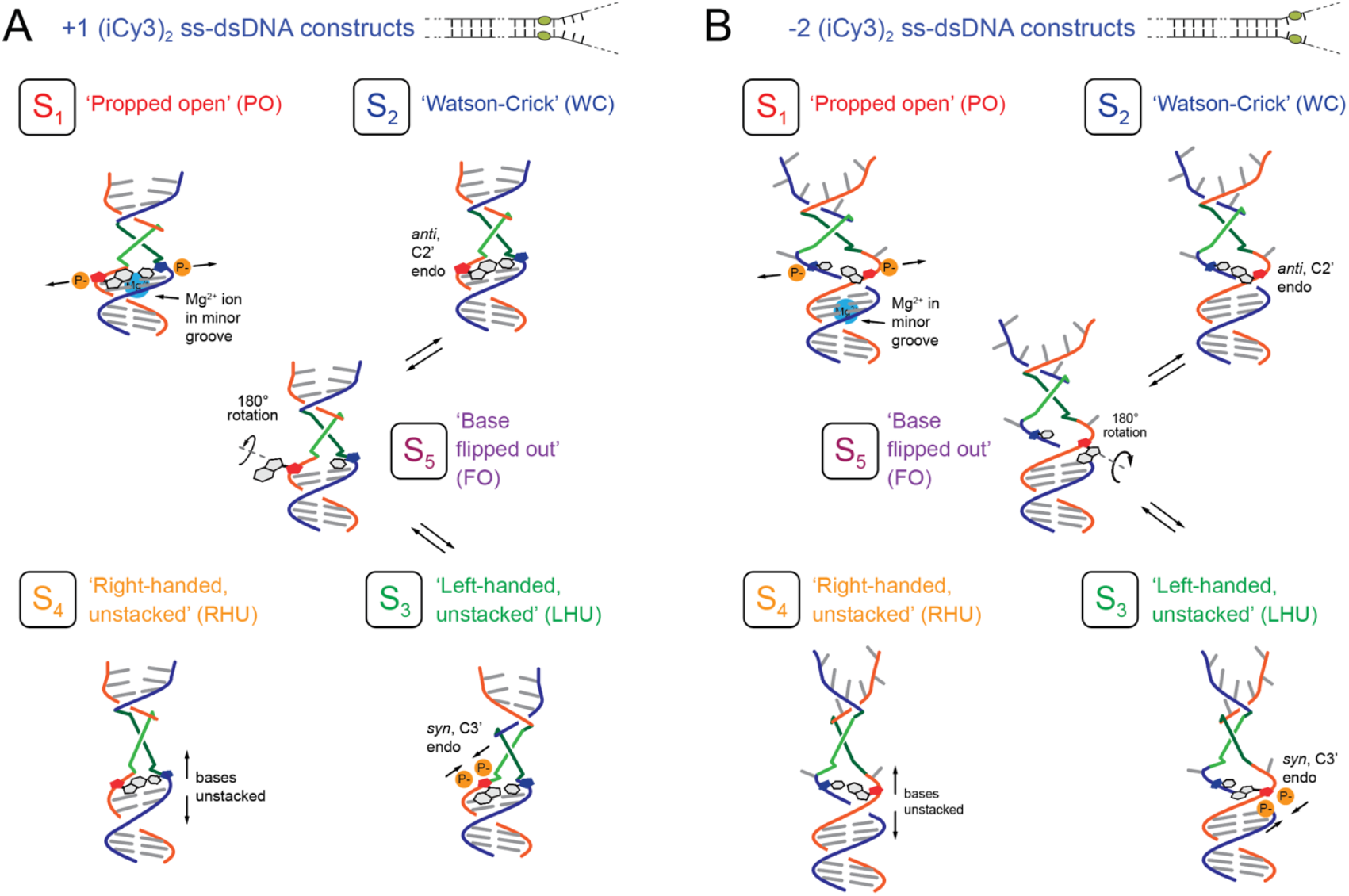
Proposed structural models for macrostates S_1_ – S_5_ of the (***A***) +1 and (***B***) -2 (iCy3)_2_ dimer-labeled ss-dsDNA fork constructs that are suggested by the results of the salt-concentration dependence studies. Complementary DNA backbones are colored blue and red, bases are gray, and the iCy3 chromophores are shown as green and blue line segments.

Thus, we note that the complementary inter-base H bonds of a single-base pair of the S_1_ (PO) macrostate are disrupted, resulting in an increased inter-strand phosphate-phosphate distance for that base-pair accompanied by reduced base stacking interactions compared to the S_2_ (WC) macrostate. The pH and salt concentration-dependences of such ‘propped-open’ states were originally studied using hydrogen-tritium exchange experiments (16-19). Here we propose that the S_1_ macrostate is preferentially stabilized by the presence of divalent magnesium ions that can bind relatively tightly within the minor groove between opposite backbone phosphates near the +1 position of the ss-dsDNA junction. The attractive Coulombic interactions between the divalent magnesium ion and the backbone phosphates of complementary nucleotide residues serve to stabilize the S_1_ macrostate. Because the minor groove is defined within the duplex portion of the ss-dsDNA junction, but not within the single-stranded sequences, the effects of magnesium cations are more pronounced in the +1 construct than in the -2 construct.

The S_2_ (WC) macrostate is depicted in Fig. 11*A*, with flanking bases adjacent to the dimer probe stacked in the *anti*-form and complementary WC base-pairing. The sugar pucker of the purine nucleotide residue closest to the probes is C2’ endo (red pentagon) such that the helical backbone twist at the +1 position of the ss-dsDNA fork junction is right-handed. In contrast, the S_3_ (LHU) macrostate is depicted with flanking bases adjacent to the dimer probe somewhat unstacked in the *syn*-form with (possibly) non-WC base-pairing. The sugar pucker of the purine nucleotide residue is C3’ endo, and the local helical backbone twist at the +1 position is left-handed. The C3’ endo sugar pucker of the S_3_ macrostate results in a smaller intra-strand phosphate-phosphate distance than the C2’ endo sugar pucker within the S_2_ macrostate. The increased negative charge density of the S_3_ macrostate (relative to the S_2_ macrostate) is thus more effectively screened and stabilized at the elevated concentration of monovalent sodium ions that are present within the ion condensation layer.

The S_4_ (RHU) macrostate is depicted in Fig. 11*A* with flanking bases adjacent to the dimer probe significantly lifted off one another (unstacked) while retaining complementary WC base pairing. Small angle X-ray scattering (SAXS) studies (44) showed that the intra-strand phosphate-phosphate distances are about the same for unstacked bases as for stacked bases. The S_4_ macrostate is thus not expected to interact preferentially with monovalent or divalent counterions, apart from its indirect coupling to macrostates that do interact strongly with ions, which is consistent with our experimental observations.

The structural assignments depicted in Fig. 11 for macrostates S_1_ – S_4_ of the +1 construct provide additional insights into the identity of macrostate S_5_, which is observed at elevated salt concentrations (300 mM NaCl, 6 mM MgCl_2_). To populate the S_3_ macrostate, the purine base must ‘flip out’ between flanking bases of the DNA framework and rotate about the glycosidic bond to transform from *anti* to *syn*. The ‘flipped out’ conformation can be viewed as a short-lived intermediate that lies on the pathway between the S_3_ macrostate and other macrostate conformations, with lifetimes too short to observe at ‘physiological’ and low salt concentrations. However, at the elevated salt concentrations for which the S_3_ macrostate is the ground state, the ‘flipped out’ state is also stabilized and thus exhibits a sufficiently long population lifetime (∼1.5 ms) to be detected as macrostate S_5_.

In Fig. 11*B*, we illustrate the structural assignments we propose for the macrostates of the -2 (iCy3)_2_ dimer-labeled ss-dsDNA construct. The structural details depicted for the -2 construct resemble those suggested for the +1 construct. Because the dimer probes of the -2 construct are positioned within the single-stranded region of the ss-dsDNA fork junction, the conformations of the bases and backbones immediately adjacent to the probes are displaced relative to those of the +1 construct. Thus, the relative stabilities of the macrostates for the -2 construct resemble, but are not identical to, those of the +1 construct. As discussed in the previous sections, the effects of varying salt concentrations on the macrostate stabilities of the -2 construct are nearly opposite to those of the +1 construct. At ‘physiological’ salt concentrations, the S_3_ (LHU) macrostate of the - 2 construct is the ground state, which might be due to the weakening of the stacking interactions within the single-stranded region of the junction. This occurs at the same ‘physiological’ salt concentrations for which the ground state conformation of the +1 construct is the S_2_ (WC) macrostate. However, at elevated salt concentration (the same elevated salt concentration for which the ground state conformation of the +1 construct is the S_3_ macrostate), the ground state conformation of the -2 construct is the S_2_ macrostate.

It is interesting to note that for the -2 construct, the stability of the S_1_ (PO) macrostate gradually decreases with increasing [MgCl_2_]/[NaCl] ratio, which is opposite to the behavior we observed for the +1 construct. At the lowest salt concentrations (20 mM NaCl, 6 mM MgCl_2_), the S_1_ macrostate is significantly destabilized in the -2 construct, while it is stabilized in the +1 construct. Elimination of magnesium ions at the lowest salt concentrations (20 mM NaCl, 0 mM MgCl_2_) affects the relative stability of S_1_ at the -2 and +1 positions in opposite directions. For the -2 construct, macrostate S_1_ is stabilized by the elimination of the magnesium counterion, while elimination of magnesium destabilizes the +1 construct. The differential stabilities of the S_1_ macrostate at the -2 and +1 positions, and their behaviors at low salt concentrations, support the notion that magnesium ions are site-specifically associated with the duplex region of the ss-dsDNA junction, in contrast to their interactions within single-stranded sequences. It is important to note the very different behaviors of the -2 and +1 constructs at the lowest salt concentrations tested (20 mM NaCl, 0 mM MgCl_2_). While the relative stabilities and transition barriers of the macrostates of the +1 construct resemble those observed under ‘physiological’ salt concentrations, in the -2 construct macrostates S_1_, S_3_ and S_4_ exhibit nearly equal stabilities and are all present at significant levels. It is under this low salt condition for the -2 construct that macrostate S_5_ is present at trace levels with a moderate population lifetime (∼290 ms).

### 4.5. Final thoughts

The PS-SMF method can sensitively monitor local fluctuations of the DNA bases and the sugar-phosphate backbones on length scales of a few angstroms and biologically relevant time scales spanning hundreds-of-microseconds to seconds. The method is thus well suited to study DNA ‘breathing,’ which is critical to understanding the mechanisms of protein-DNA interactions. The local structural and kinetic properties of ss-dsDNA fork junctions and the associated FESs revealed in the current studies can provide new insights into how the DNA replication and repair proteins that assemble at these sites carry out their biological functions. These protein complexes are critically involved in gene regulation within the biologically central processes of DNA synthesis and repair, where they recognize and interact with fluctuating local DNA secondary structures that are essentially functionally independent of specific DNA base composition and sequence. In addition, we note that the PS-SMF method described in this paper and its predecessor (1) can also be applied more generally to reveal detailed structural and dynamic information about a large variety of other protein-DNA interaction systems.

## Acknowledgements

The authors are grateful to their laboratory colleagues for many helpful discussions. This work was supported by NIH NIGMS Grant GM-15792 to A.H.M. and P.H.v.H. P.H.v.H. is an American Cancer Society Research Professor of Chemistry.

## Supporting Information

**Table S1.** Optimized parameters for the Gaussian components of the PS-SMF probability distribution functions (PDFs).

| Sample | $A_1$ | $\bar{\nu}_1$ | $\sigma_1$ | $A_2$ | $\bar{\nu}_2$ | $\sigma_2$ | $A_3$ | $\bar{\nu}_3$ | $\sigma_3$ | $A_4$ | $\bar{\nu}_4$ | $\sigma_4$ | $A_5$ | $\bar{\nu}_5$ | $\sigma_5$ |
| --- | --- | --- | --- | --- | --- | --- | --- | --- | --- | --- | --- | --- | --- | --- | --- |
| +1, 300 mM NaCl | 0.08 | 0.003 | 0.037 | 0.59 | 0.11 | 0.049 | 4.84 | 0.15 | 0.075 | 0.02 | 0.32 | 0.072 | 0.12 | 0.39 | 0.054 |
| +1, 100 mM NaCl | 0.15 | 0.06 | 0.059 | 6.11 | 0.11 | 0.053 | 1.02 | 0.14 | 0.053 | 0.18 | 0.35 | 0.048 | - | - | - |
| +1, 20mM NaCl | 1.79 | 0.06 | 0.055 | 6.07 | 0.11 | 0.048 | 0.32 | 0.17 | 0.032 | 0.02 | 0.30 | 0.053 | - | - | - |
| +1, 20mM NaCl, 0 mM MgCl<sub>2</sub> | 0.12 | 0.0001 | 0.019 | 5.31 | 0.12 | 0.062 | 1.01 | 0.14 | 0.060 | 0.19 | 0.37 | 0.028 | - | - | - |
| -1, 100mM NaCl | 0.22 | 0.0001 | 0.045 | 2.29 | 0.10 | 0.037 | 5.50 | 0.12 | 0.054 | 0.002 | 0.30 | 0.049 | - | - | - |
| -2, 300mM NaCl | 0.97 | 0.04 | 0.039 | 6.81 | 0.12 | 0.051 | 0.37 | 0.15 | 0.031 | 0.02 | 0.30 | 0.039 | - | - | - |
| -2, 100mM NaCl | 0.78 | 0.04 | 0.027 | 3.46 | 0.09 | 0.023 | 4.48 | 0.13 | 0.063 | 0.20 | 0.30 | 0.063 | - | - | - |
| -2, 20mM NaCl | 0.40 | 0.0001 | 0.038 | 6.71 | 0.12 | 0.054 | 0.39 | 0.17 | 0.044 | 0.07 | 0.30 | 0.02 | - | - | - |
| -2, 20mM NaCl, 0 mM MgCl<sub>2</sub> | 1.43 | 0.06 | 0.034 | 6.32 | 0.11 | 0.037 | 1.31 | 0.19 | 0.045 | 1.09 | 0.24 | 0.045 | 0.01 | 0.32 | 0.046 |

Parameters defined in Eq. (5). Error bars for the mean visibilities are $< 0.01$ (see Statistical Uncertainty Analysis below). PDFs were constructed using the integration period $T_w = 10$ ms. All samples prepared in 10 mM Tris at pH 8.0 and 6 mM MgCl<sub>2</sub>, unless otherwise specified.

**Table S2.** Optimized values of the equilibrium probabilities and the visibility fluctuations.

| Sample | $p_1^{eq}$ | $p_2^{eq}$ | $p_3^{eq}$ | $p_4^{eq}$ | $p_5^{eq}$ | $\delta\bar{v}_1$ | $\delta\bar{v}_2$ | $\delta\bar{v}_3$ | $\delta\bar{v}_4$ | $\delta\bar{v}_5$ | $\bar{v}$ |
| --- | --- | --- | --- | --- | --- | --- | --- | --- | --- | --- | --- |
| +1, 300 mM NaCl | 0.01 | 0.07 | 0.91 | 0.003 | 0.02 | -0.15 | -0.04 | 0.00 | 0.17 | 0.24 | 0.15 |
| +1, 100 mM NaCl | 0.02 | 0.82 | 0.14 | 0.02 | - | -0.06 | -0.01 | 0.02 | 0.23 | - | 0.12 |
| +1, 20mM NaCl | 0.25 | 0.74 | 0.02 | 0.002 | - | -0.04 | 0.01 | 0.07 | 0.20 | - | 0.10 |
| +1, 20mM NaCl, 0 mM MgCl<sub>2</sub> | 0.01 | 0.82 | 0.15 | 0.01 | - | -0.12 | 0.00 | 0.02 | 0.25 | - | 0.12 |
| -1, 100mM NaCl | 0.02 | 0.22 | 0.76 | 0.0003 | - | -0.11 | -0.01 | 0.01 | 0.19 | - | 0.11 |
| -2, 300mM NaCl | 0.10 | 0.88 | 0.03 | 0.001 | - | -0.07 | 0.01 | 0.04 | 0.19 | - | 0.11 |
| -2, 100mM NaCl | 0.05 | 0.20 | 0.71 | 0.03 | - | -0.08 | -0.03 | 0.01 | 0.18 | - | 0.12 |
| -2, 20mM NaCl | 0.04 | 0.91 | 0.04 | 0.003 | - | -0.12 | 0.00 | 0.05 | 0.18 | - | 0.12 |
| -2, 20mM NaCl, 0 mM MgCl<sub>2</sub> | 0.12 | 0.60 | 0.15 | 0.12 | 0.001 | -0.07 | -0.02 | 0.06 | 0.11 | 0.19 | 0.13 |

**Table S3.** Decomposition of largest pathway terms for the two-point TCFs 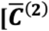function, Eq. (6) of main text] for τ = 250 *μ* s.

| +1, 300 mM NaCl, 6 mM MgCl<sub>2</sub> |  |  |  |  |  |  |  |  |  |
| --- | --- | --- | --- | --- | --- | --- | --- | --- | --- |
| Positive terms |  |  |  |  | Negative terms |  |  |  |  |
| $i$ | $j$ | $\delta\bar{v}_i \delta\bar{v}_j$ | $w_{ij}$ | $w_{ij} \delta\bar{v}_i \delta\bar{v}_j$ | $i$ | $j$ | $\delta\bar{v}_i \delta\bar{v}_j$ | $w_{ij}$ | $w_{ij} \delta\bar{v}_i \delta\bar{v}_j$ |
| 5 | 5 | $5.70 \times 10^{-2}$ | $1.40 \times 10^{-2}$ | $7.98 \times 10^{-4}$ | 1 | 5 | $-3.58 \times 10^{-2}$ | $1.84 \times 10^{-3}$ | $-6.60 \times 10^{-5}$ |
| 1 | 1 | $2.24 \times 10^{-2}$ | $5.94 \times 10^{-3}$ | $1.33 \times 10^{-4}$ | 5 | 1 | $-3.58 \times 10^{-2}$ | $1.84 \times 10^{-3}$ | $-6.60 \times 10^{-5}$ |
| 4 | 4 | $2.82 \times 10^{-2}$ | $3.96 \times 10^{-3}$ | $1.12 \times 10^{-4}$ | 3 | 5 | $-6.45 \times 10^{-4}$ | $3.79 \times 10^{-4}$ | $-2.44 \times 10^{-7}$ |
| 2 | 2 | $1.47 \times 10^{-3}$ | $7.31 \times 10^{-2}$ | $1.07 \times 10^{-4}$ | 5 | 3 | $-6.45 \times 10^{-4}$ | $3.79 \times 10^{-4}$ | $-2.44 \times 10^{-7}$ |
| 3 | 3 | $7.29 \times 10^{-6}$ | $9.10 \times 10^{-1}$ | $6.64 \times 10^{-6}$ | 2 | 5 | $-9.15 \times 10^{-3}$ | $4.35 \times 10^{-6}$ | $-3.98 \times 10^{-8}$ |
| 2 | 1 | $5.74 \times 10^{-3}$ | $2.98 \times 10^{-5}$ | $1.71 \times 10^{-7}$ | 5 | 2 | $-9.15 \times 10^{-3}$ | $3.75 \times 10^{-6}$ | $-3.43 \times 10^{-8}$ |
| 1 | 2 | $5.74 \times 10^{-3}$ | $2.57 \times 10^{-5}$ | $1.47 \times 10^{-7}$ | 4 | 3 | $-4.54 \times 10^{-4}$ | $2.72 \times 10^{-6}$ | $-1.24 \times 10^{-9}$ |
| 3 | 1 | $4.04 \times 10^{-4}$ | $2.51 \times 10^{-5}$ | $1.02 \times 10^{-8}$ | 3 | 4 | $-4.54 \times 10^{-4}$ | $1.92 \times 10^{-6}$ | $-8.73 \times 10^{-10}$ |

| +1, 100 mM NaCl, 6 mM MgCl<sub>2</sub> |  |  |  |  |  |  |  |  |  |
| --- | --- | --- | --- | --- | --- | --- | --- | --- | --- |
| Positive terms |  |  |  |  | Negative terms |  |  |  |  |
| *i* | *j* | $\delta\bar{v}_i\delta\bar{v}_j$ | $w_{ij}$ | $w_{ij}\delta\bar{v}_i\delta\bar{v}_j$ | *i* | *j* | $\delta\bar{v}_i\delta\bar{v}_j$ | $w_{ij}$ | $w_{ij}\delta\bar{v}_i\delta\bar{v}_j$ |
| 4 | 4 | $5.28 \times 10^{-2}$ | $9.07 \times 10^{-3}$ | $4.79 \times 10^{-4}$ | 1 | 4 | $-1.44 \times 10^{-2}$ | $7.06 \times 10^{-4}$ | $-1.02 \times 10^{-5}$ |
| 1 | 1 | $3.91 \times 10^{-3}$ | $2.08 \times 10^{-2}$ | $8.12 \times 10^{-5}$ | 4 | 1 | $-1.44 \times 10^{-2}$ | $7.06 \times 10^{-4}$ | $-1.01 \times 10^{-5}$ |
| 4 | 3 | $4.25 \times 10^{-3}$ | $1.15 \times 10^{-2}$ | $4.90 \times 10^{-5}$ | 1 | 3 | $-1.16 \times 10^{-3}$ | $3.42 \times 10^{-4}$ | $-3.96 \times 10^{-7}$ |
| 3 | 4 | $4.25 \times 10^{-3}$ | $1.15 \times 10^{-2}$ | $4.88 \times 10^{-5}$ | 3 | 1 | $-1.16 \times 10^{-3}$ | $3.41 \times 10^{-4}$ | $-3.95 \times 10^{-7}$ |
| 2 | 2 | $5.62 \times 10^{-5}$ | $8.19 \times 10^{-1}$ | $4.60 \times 10^{-5}$ | 2 | 4 | $-1.72 \times 10^{-3}$ | $7.67 \times 10^{-5}$ | $-1.32 \times 10^{-7}$ |
| 3 | 3 | $3.43 \times 10^{-4}$ | $1.26 \times 10^{-1}$ | $4.32 \times 10^{-5}$ | 4 | 2 | $-1.72 \times 10^{-3}$ | $4.01 \times 10^{-5}$ | $-6.91 \times 10^{-8}$ |
| 2 | 1 | $4.69 \times 10^{-4}$ | $2.21 \times 10^{-6}$ | $1.04 \times 10^{-9}$ | 3 | 2 | $-1.39 \times 10^{-4}$ | $7.74 \times 10^{-5}$ | $-1.08 \times 10^{-8}$ |
| 1 | 2 | $4.69 \times 10^{-4}$ | $1.12 \times 10^{-6}$ | $5.28 \times 10^{-10}$ | 2 | 3 | $-1.39 \times 10^{-4}$ | $3.96 \times 10^{-5}$ | $-5.50 \times 10^{-9}$ |

| +1, 20 mM NaCl, 6 mM MgCl<sub>2</sub> |  |  |  |  |  |  |  |  |  |
| --- | --- | --- | --- | --- | --- | --- | --- | --- | --- |
| Positive terms |  |  |  |  | Negative terms |  |  |  |  |
| *i* | *j* | $\delta\bar{v}_i\delta\bar{v}_j$ | $w_{ij}$ | $w_{ij}\delta\bar{v}_i\delta\bar{v}_j$ | *i* | *j* | $\delta\bar{v}_i\delta\bar{v}_j$ | $w_{ij}$ | $w_{ij}\delta\bar{v}_i\delta\bar{v}_j$ |
| 1 | 1 | $1.76 \times 10^{-3}$ | $1.64 \times 10^{-3}$ | $2.89 \times 10^{-4}$ | 1 | 2 | $-3.38 \times 10^{-4}$ | $8.48 \times 10^{-2}$ | $-2.87 \times 10^{-5}$ |
| 4 | 4 | $3.53 \times 10^{-2}$ | $2.66 \times 10^{-3}$ | $9.42 \times 10^{-5}$ | 2 | 1 | $-3.38 \times 10^{-4}$ | $8.48 \times 10^{-2}$ | $-2.87 \times 10^{-5}$ |
| 3 | 3 | $3.52 \times 10^{-3}$ | $2.56 \times 10^{-2}$ | $9.02 \times 10^{-5}$ | 4 | 1 | $-7.89 \times 10^{-3}$ | $3.84 \times 10^{-5}$ | $-3.03 \times 10^{-7}$ |
| 2 | 2 | $6.50 \times 10^{-5}$ | $6.60 \times 10^{-1}$ | $4.28 \times 10^{-5}$ | 1 | 4 | $-7.89 \times 10^{-3}$ | $3.84 \times 10^{-4}$ | $-3.03 \times 10^{-7}$ |
| 4 | 2 | $1.51 \times 10^{-3}$ | $8.88 \times 10^{-6}$ | $1.34 \times 10^{-8}$ | 3 | 1 | $-2.49 \times 10^{-3}$ | $6.24 \times 10^{-7}$ | $-1.56 \times 10^{-9}$ |
| 2 | 4 | $1.51 \times 10^{-3}$ | $8.88 \times 10^{-6}$ | $1.34 \times 10^{-8}$ | 1 | 3 | $-2.49 \times 10^{-3}$ | $9.72 \times 10^{-8}$ | $-2.42 \times 10^{-10}$ |
| 3 | 2 | $4.78 \times 10^{-5}$ | $9.33 \times 10^{-6}$ | $4.47 \times 10^{-9}$ | | | | | |
| 2 | 3 | $4.78 \times 10^{-4}$ | $1.45 \times 10^{-6}$ | $6.95 \times 10^{-10}$ | | | | | |

| +1, 20 mM NaCl, 0 mM MgCl<sub>2</sub> |  |  |  |  |  |  |  |  |  |
| --- | --- | --- | --- | --- | --- | --- | --- | --- | --- |
| Positive terms |  |  |  |  | Negative terms |  |  |  |  |
| *i* | *j* | $\delta\bar{v}_i\delta\bar{v}_j$ | $w_{ij}$ | $w_{ij}\delta\bar{v}_i\delta\bar{v}_j$ | *i* | *j* | $\delta\bar{v}_i\delta\bar{v}_j$ | $w_{ij}$ | $w_{ij}\delta\bar{v}_i\delta\bar{v}_j$ |
| 4 | 4 | $5.80 \times 10^{-2}$ | $7.17 \times 10^{-3}$ | $4.16 \times 10^{-4}$ | 1 | 3 | $-1.75 \times 10^{-3}$ | $3.03 \times 10^{-4}$ | $-5.30 \times 10^{-7}$ |
| 1 | 1 | $1.64 \times 10^{-2}$ | $5.61 \times 10^{-3}$ | $9.27 \times 10^{-5}$ | 3 | 1 | $-1.75 \times 10^{-3}$ | $3.03 \times 10^{-4}$ | $-5.30 \times 10^{-7}$ |
| 3 | 3 | $1.85 \times 10^{-4}$ | $1.48 \times 10^{-1}$ | $2.74 \times 10^{-5}$ | 2 | 4 | $-1.36 \times 10^{-3}$ | $2.09 \times 10^{-4}$ | $-2.83 \times 10^{-7}$ |
| 2 | 2 | $3.18 \times 10^{-5}$ | $8.25 \times 10^{-1}$ | $2.62 \times 10^{-5}$ | 4 | 1 | $-3.10 \times 10^{-2}$ | $7.43 \times 10^{-6}$ | $-2.30 \times 10^{-7}$ |
| 4 | 3 | $3.28 \times 10^{-3}$ | $6.54 \times 10^{-3}$ | $2.14 \times 10^{-5}$ | 1 | 4 | $-3.10 \times 10^{-2}$ | $7.21 \times 10^{-6}$ | $-2.23 \times 10^{-7}$ |
| 3 | 4 | $3.28 \times 10^{-3}$ | $6.35 \times 10^{-3}$ | $2.08 \times 10^{-5}$ | 3 | 2 | $-7.67 \times 10^{-5}$ | $3.49 \times 10^{-4}$ | $-2.68 \times 10^{-8}$ |
| 1 | 2 | $7.24 \times 10^{-4}$ | $3.55 \times 10^{-7}$ | $2.57 \times 10^{-10}$ | 4 | 2 | $-1.35 \times 10^{-3}$ | $1.53 \times 10^{-5}$ | $-2.07 \times 10^{-8}$ |
| 2 | 1 | $7.24 \times 10^{-4}$ | $1.36 \times 10^{-7}$ | $9.86 \times 10^{-11}$ | 2 | 3 | $-7.67 \times 10^{-5}$ | $1.56 \times 10^{-4}$ | $-1.19 \times 10^{-8}$ |

| -1, 100 mM NaCl, 6 mM MgCl<sub>2</sub> |  |  |  |  |  |  |  |  |  |
| --- | --- | --- | --- | --- | --- | --- | --- | --- | --- |
| Positive terms |  |  |  |  | Negative terms |  |  |  |  |
| *i* | *j* | $\delta\bar{v}_i\delta\bar{v}_j$ | $w_{ij}$ | $w_{ij}\delta\bar{v}_i\delta\bar{v}_j$ | *i* | *j* | $\delta\bar{v}_i\delta\bar{v}_j$ | $w_{ij}$ | $w_{ij}\delta\bar{v}_i\delta\bar{v}_j$ |
| 1 | 1 | $1.31 \times 10^{-2}$ | $2.06 \times 10^{-2}$ | $2.70 \times 10^{-4}$ | 3 | 1 | $-8.88 \times 10^{-4}$ | $4.08 \times 10^{-3}$ | $-3.63 \times 10^{-6}$ |
| 3 | 3 | $6.02 \times 10^{-5}$ | $7.52 \times 10^{-1}$ | $4.53 \times 10^{-5}$ | 1 | 3 | $-8.88 \times 10^{-4}$ | $3.97 \times 10^{-3}$ | $-3.53 \times 10^{-6}$ |
| 2 | 2 | $1.85 \times 10^{-4}$ | $2.16 \times 10^{-1}$ | $4.00 \times 10^{-5}$ | 2 | 3 | $-1.06 \times 10^{-4}$ | $2.36 \times 10^{-4}$ | $-2.49 \times 10^{-8}$ |
| 4 | 4 | $3.44 \times 10^{-2}$ | $2.75 \times 10^{-4}$ | $9.47 \times 10^{-6}$ | 4 | 1 | $-2.12 \times 10^{-2}$ | $7.44 \times 10^{-7}$ | $-1.58 \times 10^{-8}$ |
| 1 | 2 | $1.56 \times 10^{-3}$ | $1.35 \times 10^{-3}$ | $2.10 \times 10^{-6}$ | 1 | 4 | $-2.12 \times 10^{-2}$ | $7.44 \times 10^{-7}$ | $-1.58 \times 10^{-8}$ |
| 2 | 1 | $1.56 \times 10^{-3}$ | $1.24 \times 10^{-3}$ | $1.93 \times 10^{-6}$ | 3 | 2 | $-1.06 \times 10^{-4}$ | $1.26 \times 10^{-4}$ | $-1.33 \times 10^{-8}$ |
| 3 | 4 | $1.44 \times 10^{-3}$ | $6.83 \times 10^{-8}$ | $9.84 \times 10^{-11}$ | 4 | 2 | $-2.52 \times 10^{-3}$ | $2.26 \times 10^{-8}$ | $-5.70 \times 10^{-11}$ |
| 4 | 3 | $1.44 \times 10^{-3}$ | $6.65 \times 10^{-8}$ | $9.57 \times 10^{-11}$ | 2 | 4 | $-2.52 \times 10^{-3}$ | $2.07 \times 10^{-8}$ | $-5.24 \times 10^{-11}$ |

| -2, 300 mM NaCl, 6 mM MgCl<sub>2</sub> |  |  |  |  |  |  |  |  |  |
| --- | --- | --- | --- | --- | --- | --- | --- | --- | --- |
| Positive terms |  |  |  |  | Negative terms |  |  |  |  |
| *i* | *j* | $\delta \bar{v}_i \delta \bar{v}_j$ | $w_{ij}$ | $w_{ij} \delta \bar{v}_i \delta \bar{v}_j$ | *i* | *j* | $\delta \bar{v}_i \delta \bar{v}_j$ | $w_{ij}$ | $w_{ij} \delta \bar{v}_i \delta \bar{v}_j$ |
| 1 | 1 | $5.52 \times 10^{-3}$ | $5.27 \times 10^{-2}$ | $2.91 \times 10^{-4}$ | 1 | 2 | $-3.89 \times 10^{-4}$ | $4.27 \times 10^{-2}$ | $-1.66 \times 10^{-5}$ |
| 4 | 4 | $3.44 \times 10^{-2}$ | $1.88 \times 10^{-3}$ | $6.46 \times 10^{-5}$ | 2 | 1 | $-3.89 \times 10^{-4}$ | $4.27 \times 10^{-2}$ | $-1.66 \times 10^{-5}$ |
| 3 | 3 | $1.74 \times 10^{-3}$ | $2.88 \times 10^{-2}$ | $5.02 \times 10^{-5}$ | 4 | 1 | $-1.38 \times 10^{-2}$ | $3.28 \times 10^{-5}$ | $-4.52 \times 10^{-7}$ |
| 2 | 2 | $2.74 \times 10^{-5}$ | $8.39 \times 10^{-1}$ | $2.30 \times 10^{-5}$ | 1 | 4 | $-1.38 \times 10^{-2}$ | $3.28 \times 10^{-5}$ | $-4.52 \times 10^{-7}$ |
| 4 | 2 | $9.71 \times 10^{-4}$ | $1.09 \times 10^{-5}$ | $1.06 \times 10^{-8}$ | 3 | 1 | $-3.10 \times 10^{-3}$ | $3.74 \times 10^{-7}$ | $-1.16 \times 10^{-9}$ |
| 2 | 4 | $9.71 \times 10^{-4}$ | $1.09 \times 10^{-5}$ | $1.06 \times 10^{-8}$ | 1 | 3 | $-3.10 \times 10^{-3}$ | $2.72 \times 10^{-7}$ | $-8.46 \times 10^{-10}$ |
| 3 | 2 | $2.18 \times 10^{-4}$ | $1.35 \times 10^{-5}$ | $2.95 \times 10^{-9}$ | | | | | |
| 2 | 3 | $2.18 \times 10^{-4}$ | $9.84 \times 10^{-6}$ | $2.15 \times 10^{-9}$ | | | | | |

| -2, 100 mM NaCl, 6 mM MgCl<sub>2</sub> |  |  |  |  |  |  |  |  |  |
| --- | --- | --- | --- | --- | --- | --- | --- | --- | --- |
| Positive terms |  |  |  |  | Negative terms |  |  |  |  |
| *i* | *j* | $\delta \bar{v}_i \delta \bar{v}_j$ | $w_{ij}$ | $w_{ij} \delta \bar{v}_i \delta \bar{v}_j$ | *i* | *j* | $\delta \bar{v}_i \delta \bar{v}_j$ | $w_{ij}$ | $w_{ij} \delta \bar{v}_i \delta \bar{v}_j$ |
| 4 | 4 | $3.02 \times 10^{-2}$ | $1.98 \times 10^{-2}$ | $5.98 \times 10^{-4}$ | 4 | 1 | $-1.33 \times 10^{-2}$ | $1.15 \times 10^{-2}$ | $-1.54 \times 10^{-5}$ |
| 1 | 1 | $5.86 \times 10^{-3}$ | $3.74 \times 10^{-2}$ | $2.19 \times 10^{-4}$ | 1 | 4 | $-1.33 \times 10^{-2}$ | $1.15 \times 10^{-2}$ | $-1.54 \times 10^{-5}$ |
| 2 | 2 | $9.22 \times 10^{-4}$ | $1.97 \times 10^{-1}$ | $1.82 \times 10^{-4}$ | 2 | 4 | $-5.28 \times 10^{-3}$ | $7.63 \times 10^{-4}$ | $-4.03 \times 10^{-6}$ |
| 3 | 3 | $4.41 \times 10^{-5}$ | $7.10 \times 10^{-1}$ | $3.13 \times 10^{-5}$ | 4 | 2 | $-5.28 \times 10^{-3}$ | $7.62 \times 10^{-4}$ | $-4.02 \times 10^{-6}$ |
| 1 | 2 | $2.32 \times 10^{-3}$ | $4.95 \times 10^{-3}$ | $1.15 \times 10^{-5}$ | 2 | 3 | $-2.02 \times 10^{-4}$ | $4.57 \times 10^{-4}$ | $-9.22 \times 10^{-8}$ |
| 2 | 1 | $2.32 \times 10^{-3}$ | $4.95 \times 10^{-3}$ | $1.15 \times 10^{-5}$ | 3 | 2 | $-2.02 \times 10^{-4}$ | $4.57 \times 10^{-4}$ | $-9.22 \times 10^{-8}$ |
| 3 | 4 | $1.15 \times 10^{-3}$ | $6.32 \times 10^{-7}$ | $7.30 \times 10^{-10}$ | 1 | 3 | $-5.08 \times 10^{-4}$ | $6.04 \times 10^{-6}$ | $-3.07 \times 10^{-9}$ |
| 4 | 3 | $1.15 \times 10^{-3}$ | $6.30 \times 10^{-7}$ | $7.28 \times 10^{-10}$ | 3 | 1 | $-5.08 \times 10^{-4}$ | $6.04 \times 10^{-6}$ | $-3.07 \times 10^{-9}$ |

| -2, 20 mM NaCl, 6 mM MgCl<sub>2</sub> |  |  |  |  |  |  |  |  |  |
| --- | --- | --- | --- | --- | --- | --- | --- | --- | --- |
| Positive terms |  |  |  |  | Negative terms |  |  |  |  |
| *i* | *j* | $\delta \bar{v}_i \delta \bar{v}_j$ | $w_{ij}$ | $w_{ij} \delta \bar{v}_i \delta \bar{v}_j$ | *i* | *j* | $\delta \bar{v}_i \delta \bar{v}_j$ | $w_{ij}$ | $w_{ij} \delta \bar{v}_i \delta \bar{v}_j$ |
| 1 | 1 | $1.36 \times 10^{-2}$ | $2.51 \times 10^{-2}$ | $3.40 \times 10^{-4}$ | 1 | 2 | $-1.95 \times 10^{-4}$ | $1.41 \times 10^{-2}$ | $-2.75 \times 10^{-6}$ |
| 3 | 3 | $2.62 \times 10^{-3}$ | $4.38 \times 10^{-2}$ | $1.15 \times 10^{-4}$ | 2 | 1 | $-1.95 \times 10^{-4}$ | $1.41 \times 10^{-2}$ | $-2.74 \times 10^{-6}$ |
| 4 | 4 | $3.35 \times 10^{-2}$ | $3.36 \times 10^{-3}$ | $1.12 \times 10^{-4}$ | 4 | 1 | $-2.13 \times 10^{-3}$ | $4.36 \times 10^{-5}$ | $-9.29 \times 10^{-7}$ |
| 2 | 2 | $2.80 \times 10^{-6}$ | $9.01 \times 10^{-1}$ | $2.52 \times 10^{-6}$ | 1 | 4 | $-2.13 \times 10^{-3}$ | $4.36 \times 10^{-5}$ | $-9.29 \times 10^{-7}$ |
| 4 | 2 | $3.06 \times 10^{-4}$ | $1.05 \times 10^{-5}$ | $3.22 \times 10^{-9}$ | 3 | 1 | $-5.96 \times 10^{-3}$ | $1.79 \times 10^{-5}$ | $-1.07 \times 10^{-7}$ |
| 2 | 4 | $3.06 \times 10^{-4}$ | $1.05 \times 10^{-5}$ | $3.22 \times 10^{-9}$ | 1 | 3 | $-5.96 \times 10^{-3}$ | $3.76 \times 10^{-7}$ | $-2.24 \times 10^{-9}$ |
| 2 | 3 | $8.56 \times 10^{-5}$ | $2.68 \times 10^{-5}$ | $2.30 \times 10^{-9}$ | | | | | |
| 3 | 2 | $8.56 \times 10^{-5}$ | $1.02 \times 10^{-5}$ | $8.70 \times 10^{-10}$ | | | | | |

| -2, 20 mM NaCl, 0 mM MgCl<sub>2</sub> |  |  |  |  |  |  |  |  |  |
| --- | --- | --- | --- | --- | --- | --- | --- | --- | --- |
| Positive terms |  |  |  |  | Negative terms |  |  |  |  |
| *i* | *j* | $\delta \bar{v}_i \delta \bar{v}_j$ | $w_{ij}$ | $w_{ij} \delta \bar{v}_i \delta \bar{v}_j$ | *i* | *j* | $\delta \bar{v}_i \delta \bar{v}_j$ | $w_{ij}$ | $w_{ij} \delta \bar{v}_i \delta \bar{v}_j$ |
| 4 | 4 | $1.24 \times 10^{-2}$ | $6.11 \times 10^{-2}$ | $7.61 \times 10^{-4}$ | 4 | 1 | $-8.86 \times 10^{-3}$ | $6.11 \times 10^{-2}$ | $-5.41 \times 10^{-4}$ |
| 1 | 1 | $6.30 \times 10^{-3}$ | $6.11 \times 10^{-2}$ | $3.85 \times 10^{-4}$ | 1 | 4 | $-8.86 \times 10^{-3}$ | $6.11 \times 10^{-2}$ | $-5.41 \times 10^{-4}$ |
| 3 | 3 | $2.79 \times 10^{-3}$ | $1.20 \times 10^{-1}$ | $3.35 \times 10^{-4}$ | 2 | 3 | $-1.06 \times 10^{-3}$ | $2.81 \times 10^{-2}$ | $-2.98 \times 10^{-5}$ |
| 2 | 2 | $4.05 \times 10^{-4}$ | $5.73 \times 10^{-1}$ | $2.32 \times 10^{-4}$ | 3 | 2 | $-1.06 \times 10^{-3}$ | $2.81 \times 10^{-2}$ | $-2.98 \times 10^{-5}$ |
| 5 | 5 | $3.29 \times 10^{-2}$ | $5.90 \times 10^{-4}$ | $1.94 \times 10^{-5}$ | 1 | 3 | $-4.19 \times 10^{-3}$ | $1.32 \times 10^{-3}$ | $-5.53 \times 10^{-6}$ |
| 3 | 4 | $5.89 \times 10^{-3}$ | $1.32 \times 10^{-3}$ | $7.76 \times 10^{-6}$ | 3 | 1 | $-4.19 \times 10^{-3}$ | $1.32 \times 10^{-3}$ | $-5.53 \times 10^{-6}$ |
| 4 | 3 | $5.89 \times 10^{-3}$ | $1.32 \times 10^{-3}$ | $7.76 \times 10^{-6}$ | 2 | 4 | $-2.24 \times 10^{-3}$ | $1.44 \times 10^{-4}$ | $-3.23 \times 10^{-7}$ |
| 1 | 2 | $1.60 \times 10^{-3}$ | $1.44 \times 10^{-4}$ | $2.30 \times 10^{-7}$ | 4 | 2 | $-2.24 \times 10^{-3}$ | $1.44 \times 10^{-4}$ | $-3.23 \times 10^{-7}$ |

**Table S4.** Decomposition of largest pathway terms for the three-point TCFs 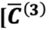function, Eq. (7) of main text] for τ _1_ = τ _2_ = 10 ms.

| +1, 300 mM NaCl, 6 mM MgCl<sub>2</sub> |  |  |  |  |  |  |  |  |  |  |  |
| --- | --- | --- | --- | --- | --- | --- | --- | --- | --- | --- | --- |
| Positive terms |  |  |  |  |  | Negative terms |  |  |  |  |  |
| $i$ | $j$ | $k$ | $\delta\bar{v}_i\delta\bar{v}_j\delta\bar{v}_k$ | $w_{ijk}$ | $w_{ijk}\delta\bar{v}_i\delta\bar{v}_j\delta\bar{v}_k$ | $i$ | $j$ | $k$ | $\delta\bar{v}_i\delta\bar{v}_j\delta\bar{v}_k$ | $w_{ijk}$ | $w_{ijk}\delta\bar{v}_i\delta\bar{v}_j\delta\bar{v}_k$ |
| 5 | 5 | 5 | $1.36 \times 10^{-2}$ | $1.66 \times 10^{-3}$ | $2.66 \times 10^{-5}$ | 5 | 1 | 5 | $-8.54 \times 10^{-3}$ | $8.86 \times 10^{-4}$ | $-7.56 \times 10^{-6}$ |
| 4 | 4 | 4 | $4.73 \times 10^{-3}$ | $3.78 \times 10^{-3}$ | $1.79 \times 10^{-5}$ | 1 | 5 | 5 | $-8.54 \times 10^{-3}$ | $8.44 \times 10^{-4}$ | $-7.21 \times 10^{-6}$ |
| 1 | 1 | 5 | $5.36 \times 10^{-3}$ | $4.49 \times 10^{-4}$ | $2.40 \times 10^{-6}$ | 5 | 5 | 1 | $-8.54 \times 10^{-3}$ | $8.44 \times 10^{-4}$ | $-7.21 \times 10^{-6}$ |
| 5 | 1 | 1 | $5.36 \times 10^{-3}$ | $4.49 \times 10^{-4}$ | $2.40 \times 10^{-6}$ | 2 | 2 | 2 | $-5.64 \times 10^{-5}$ | $7.06 \times 10^{-2}$ | $-3.98 \times 10^{-6}$ |
| 1 | 5 | 1 | $5.36 \times 10^{-3}$ | $4.28 \times 10^{-4}$ | $2.29 \times 10^{-6}$ | 1 | 1 | 1 | $-3.36 \times 10^{-3}$ | $2.28 \times 10^{-4}$ | $-7.65 \times 10^{-7}$ |
| 2 | 2 | 5 | $3.51 \times 10^{-4}$ | $6.03 \times 10^{-4}$ | $2.12 \times 10^{-7}$ | 2 | 5 | 5 | $-2.19 \times 10^{-3}$ | $1.97 \times 10^{-4}$ | $-4.30 \times 10^{-7}$ |
| 5 | 2 | 2 | $3.51 \times 10^{-4}$ | $5.18 \times 10^{-4}$ | $1.82 \times 10^{-7}$ | 3 | 5 | 5 | $-1.56 \times 10^{-4}$ | $2.52 \times 10^{-3}$ | $-3.93 \times 10^{-7}$ |
| 2 | 1 | 5 | $1.37 \times 10^{-3}$ | $1.30 \times 10^{-4}$ | $1.78 \times 10^{-7}$ | 5 | 5 | 3 | $-1.56 \times 10^{-4}$ | $2.52 \times 10^{-3}$ | $-3.93 \times 10^{-7}$ |

| +1, 100 mM NaCl, 6 mM MgCl<sub>2</sub> |  |  |  |  |  |  |  |  |  |  |  |
| --- | --- | --- | --- | --- | --- | --- | --- | --- | --- | --- | --- |
| Positive terms |  |  |  |  |  | Negative terms |  |  |  |  |  |
| $i$ | $j$ | $k$ | $\delta\bar{v}_i\delta\bar{v}_j\delta\bar{v}_k$ | $w_{ijk}$ | $w_{ijk}\delta\bar{v}_i\delta\bar{v}_j\delta\bar{v}_k$ | $i$ | $j$ | $k$ | $\delta\bar{v}_i\delta\bar{v}_j\delta\bar{v}_k$ | $w_{ijk}$ | $w_{ijk}\delta\bar{v}_i\delta\bar{v}_j\delta\bar{v}_k$ |
| 4 | 4 | 4 | $1.21 \times 10^{-2}$ | $2.80 \times 10^{-4}$ | $3.40 \times 10^{-6}$ | 1 | 4 | 4 | $-3.30 \times 10^{-3}$ | $2.68 \times 10^{-4}$ | $-8.83 \times 10^{-7}$ |
| 4 | 3 | 4 | $9.77 \times 10^{-4}$ | $1.84 \times 10^{-3}$ | $1.80 \times 10^{-6}$ | 4 | 4 | 1 | $-3.30 \times 10^{-3}$ | $2.68 \times 10^{-4}$ | $-8.83 \times 10^{-7}$ |
| 4 | 4 | 3 | $9.77 \times 10^{-4}$ | $1.83 \times 10^{-3}$ | $1.79 \times 10^{-6}$ | 4 | 1 | 4 | $-3.30 \times 10^{-3}$ | $2.50 \times 10^{-4}$ | $-8.24 \times 10^{-7}$ |
| 3 | 4 | 4 | $9.77 \times 10^{-4}$ | $1.82 \times 10^{-3}$ | $1.78 \times 10^{-6}$ | 1 | 4 | 3 | $-2.66 \times 10^{-4}$ | $1.74 \times 10^{-3}$ | $-4.64 \times 10^{-7}$ |
| 4 | 3 | 3 | $7.88 \times 10^{-5}$ | $1.20 \times 10^{-2}$ | $9.47 \times 10^{-7}$ | 3 | 4 | 1 | $-2.66 \times 10^{-4}$ | $1.74 \times 10^{-3}$ | $-4.63 \times 10^{-7}$ |
| 3 | 3 | 4 | $7.88 \times 10^{-5}$ | $1.20 \times 10^{-2}$ | $9.44 \times 10^{-7}$ | 1 | 3 | 4 | $-2.66 \times 10^{-4}$ | $1.66 \times 10^{-3}$ | $-4.41 \times 10^{-7}$ |
| 3 | 4 | 3 | $7.88 \times 10^{-5}$ | $1.19 \times 10^{-2}$ | $9.36 \times 10^{-7}$ | 4 | 3 | 1 | $-2.66 \times 10^{-4}$ | $1.66 \times 10^{-3}$ | $-4.41 \times 10^{-7}$ |
| 3 | 3 | 3 | $6.34 \times 10^{-6}$ | $7.84 \times 10^{-2}$ | $4.898 \times 10^{-7}$ | 4 | 1 | 3 | $-2.66 \times 10^{-4}$ | $1.54 \times 10^{-3}$ | $-4.09 \times 10^{-7}$ |

| +1, 20 mM NaCl, 6 mM MgCl<sub>2</sub> |  |  |  |  |  |  |  |  |  |  |  |
| --- | --- | --- | --- | --- | --- | --- | --- | --- | --- | --- | --- |
| Positive terms |  |  |  |  |  | Negative terms |  |  |  |  |  |
| *i* | *j* | *k* | $\delta\bar{v}_i\delta\bar{v}_j\delta\bar{v}_k$ | $w_{ijk}$ | $w_{ijk}\delta\bar{v}_i\delta\bar{v}_j\delta\bar{v}_k$ | *i* | *j* | *k* | $\delta\bar{v}_i\delta\bar{v}_j\delta\bar{v}_k$ | $w_{ijk}$ | $w_{ijk}\delta\bar{v}_i\delta\bar{v}_j\delta\bar{v}_k$ |
| 3 | 3 | 3 | $2.09 \times 10^{-4}$ | $2.51 \times 10^{-2}$ | $5.24 \times 10^{-6}$ | 1 | 1 | 1 | $-7.41 \times 10^{-5}$ | $1.56 \times 10^{-2}$ | $-1.16 \times 10^{-6}$ |
| 4 | 4 | 4 | $6.64 \times 10^{-3}$ | $6.65 \times 10^{-4}$ | $4.41 \times 10^{-6}$ | 1 | 2 | 2 | $-2.69 \times 10^{-6}$ | $1.39 \times 10^{-1}$ | $-3.75 \times 10^{-7}$ |
| 1 | 2 | 1 | $1.41 \times 10^{-5}$ | $4.67 \times 10^{-2}$ | $6.59 \times 10^{-7}$ | 2 | 2 | 1 | $-2.69 \times 10^{-6}$ | $1.39 \times 10^{-1}$ | $-3.75 \times 10^{-7}$ |
| 1 | 1 | 2 | $1.41 \times 10^{-5}$ | $4.67 \times 10^{-2}$ | $6.59 \times 10^{-7}$ | 2 | 1 | 2 | $-2.69 \times 10^{-6}$ | $1.39 \times 10^{-1}$ | $-3.75 \times 10^{-7}$ |
| 2 | 1 | 1 | $1.41 \times 10^{-5}$ | $4.67 \times 10^{-2}$ | $6.59 \times 10^{-7}$ | 4 | 4 | 1 | $-1.48 \times 10^{-3}$ | $1.85 \times 10^{-4}$ | $-2.75 \times 10^{-7}$ |
| 2 | 2 | 2 | $5.12 \times 10^{-7}$ | $4.16 \times 10^{-1}$ | $2.13 \times 10^{-7}$ | 1 | 4 | 4 | $-1.48 \times 10^{-3}$ | $1.85 \times 10^{-4}$ | $-2.75 \times 10^{-7}$ |
| 4 | 4 | 2 | $2.83 \times 10^{-4}$ | $4.93 \times 10^{-4}$ | $1.39 \times 10^{-7}$ | 4 | 1 | 2 | $-6.32 \times 10^{-5}$ | $2.79 \times 10^{-4}$ | $-1.76 \times 10^{-8}$ |
| 2 | 4 | 4 | $2.83 \times 10^{-4}$ | $4.93 \times 10^{-4}$ | $1.39 \times 10^{-7}$ | 2 | 1 | 4 | $-6.32 \times 10^{-5}$ | $2.79 \times 10^{-4}$ | $-1.76 \times 10^{-8}$ |

| +1, 20 mM NaCl, 0 mM MgCl<sub>2</sub> |  |  |  |  |  |  |  |  |  |  |  |
| --- | --- | --- | --- | --- | --- | --- | --- | --- | --- | --- | --- |
| Positive terms |  |  |  |  |  | Negative terms |  |  |  |  |  |
| *i* | *j* | *k* | $\delta\bar{v}_i\delta\bar{v}_j\delta\bar{v}_k$ | $w_{ijk}$ | $w_{ijk}\delta\bar{v}_i\delta\bar{v}_j\delta\bar{v}_k$ | *i* | *j* | *k* | $\delta\bar{v}_i\delta\bar{v}_j\delta\bar{v}_k$ | $w_{ijk}$ | $w_{ijk}\delta\bar{v}_i\delta\bar{v}_j\delta\bar{v}_k$ |
| 4 | 4 | 4 | $1.40 \times 10^{-2}$ | $6.93 \times 10^{-5}$ | $9.69 \times 10^{-7}$ | 1 | 1 | 1 | $-2.12 \times 10^{-3}$ | $1.11 \times 10^{-4}$ | $-2.36 \times 10^{-7}$ |
| 4 | 4 | 3 | $7.90 \times 10^{-4}$ | $8.02 \times 10^{-4}$ | $6.34 \times 10^{-7}$ | 4 | 4 | 1 | $-7.46 \times 10^{-3}$ | $2.78 \times 10^{-5}$ | $-2.07 \times 10^{-7}$ |
| 4 | 3 | 4 | $7.90 \times 10^{-4}$ | $8.00 \times 10^{-4}$ | $6.32 \times 10^{-7}$ | 1 | 4 | 4 | $-7.46 \times 10^{-3}$ | $2.70 \times 10^{-5}$ | $-2.01 \times 10^{-7}$ |
| 3 | 4 | 4 | $7.90 \times 10^{-4}$ | $7.78 \times 10^{-4}$ | $6.15 \times 10^{-7}$ | 4 | 1 | 4 | $-7.46 \times 10^{-3}$ | $2.51 \times 10^{-5}$ | $-1.87 \times 10^{-7}$ |
| 4 | 3 | 3 | $4.47 \times 10^{-5}$ | $9.25 \times 10^{-3}$ | $4.13 \times 10^{-7}$ | 2 | 2 | 2 | $-1.79 \times 10^{-7}$ | $7.98 \times 10^{-1}$ | $-1.43 \times 10^{-7}$ |
| 3 | 4 | 3 | $4.47 \times 10^{-5}$ | $9.00 \times 10^{-3}$ | $4.02 \times 10^{-7}$ | 4 | 3 | 1 | $-4.22 \times 10^{-4}$ | $3.24 \times 10^{-4}$ | $-1.37 \times 10^{-7}$ |
| 3 | 3 | 4 | $4.47 \times 10^{-5}$ | $8.98 \times 10^{-3}$ | $4.01 \times 10^{-7}$ | 1 | 3 | 4 | $-4.22 \times 10^{-4}$ | $3.15 \times 10^{-4}$ | $-1.33 \times 10^{-7}$ |
| 3 | 3 | 3 | $2.52 \times 10^{-6}$ | $1.04 \times 10^{-1}$ | $2.62 \times 10^{-7}$ | 3 | 4 | 1 | $-4.22 \times 10^{-4}$ | $3.12 \times 10^{-4}$ | $-1.32 \times 10^{-7}$ |

| -1, 100 mM NaCl, 6 mM MgCl<sub>2</sub> |  |  |  |  |  |  |  |  |  |  |  |
| --- | --- | --- | --- | --- | --- | --- | --- | --- | --- | --- | --- |
| Positive terms |  |  |  |  |  | Negative terms |  |  |  |  |  |
| $i$ | $j$ | $k$ | $\delta \bar{v}_i \delta \bar{v}_j \delta \bar{v}_k$ | $w_{ijk}$ | $w_{ijk} \delta \bar{v}_i \delta \bar{v}_j \delta \bar{v}_k$ | $i$ | $j$ | $k$ | $\delta \bar{v}_i \delta \bar{v}_j \delta \bar{v}_k$ | $w_{ijk}$ | $w_{ijk} \delta \bar{v}_i \delta \bar{v}_j \delta \bar{v}_k$ |
| 4 | 4 | 4 | $6.39 \times 10^{-3}$ | $2.17 \times 10^{-4}$ | $1.38 \times 10^{-6}$ | 2 | 2 | 2 | $-2.52 \times 10^{-6}$ | $1.41 \times 10^{-1}$ | $-3.56 \times 10^{-7}$ |
| 3 | 3 | 3 | $4.67 \times 10^{-7}$ | $6.50 \times 10^{-1}$ | $3.04 \times 10^{-7}$ | 3 | 3 | 1 | $-6.89 \times 10^{-6}$ | $1.80 \times 10^{-2}$ | $-1.24 \times 10^{-7}$ |
| 3 | 1 | 2 | $1.21 \times 10^{-5}$ | $4.66 \times 10^{-3}$ | $5.64 \times 10^{-8}$ | 1 | 3 | 3 | $-6.89 \times 10^{-6}$ | $1.76 \times 10^{-2}$ | $-1.21 \times 10^{-7}$ |
| 2 | 1 | 3 | $1.21 \times 10^{-5}$ | $4.67 \times 10^{-3}$ | $5.16 \times 10^{-8}$ | 1 | 2 | 2 | $-2.12 \times 10^{-5}$ | $5.02 \times 10^{-3}$ | $-1.06 \times 10^{-7}$ |
| 3 | 1 | 1 | $1.02 \times 10^{-4}$ | $5.04 \times 10^{-4}$ | $5.13 \times 10^{-8}$ | 2 | 2 | 1 | $-2.12 \times 10^{-5}$ | $4.69 \times 10^{-3}$ | $-9.94 \times 10^{-8}$ |
| 1 | 1 | 3 | $1.02 \times 10^{-4}$ | $4.93 \times 10^{-4}$ | $5.02 \times 10^{-8}$ | 3 | 1 | 3 | $-6.89 \times 10^{-6}$ | $1.42 \times 10^{-2}$ | $-9.80 \times 10^{-8}$ |
| 1 | 3 | 1 | $1.02 \times 10^{-4}$ | $4.86 \times 10^{-4}$ | $4.95 \times 10^{-8}$ | 1 | 2 | 1 | $-1.78 \times 10^{-4}$ | $1.67 \times 10^{-4}$ | $-2.97 \times 10^{-8}$ |
| 2 | 2 | 3 | $1.44 \times 10^{-6}$ | $2.93 \times 10^{-2}$ | $4.21 \times 10^{-8}$ | 2 | 1 | 2 | $-2.12 \times 10^{-5}$ | $1.40 \times 10^{-3}$ | $-2.97 \times 10^{-8}$ |

| -2, 300 mM NaCl, 6 mM MgCl<sub>2</sub> |  |  |  |  |  |  |  |  |  |  |  |
| --- | --- | --- | --- | --- | --- | --- | --- | --- | --- | --- | --- |
| Positive terms |  |  |  |  |  | Negative terms |  |  |  |  |  |
| $i$ | $j$ | $k$ | $\delta \bar{v}_i \delta \bar{v}_j \delta \bar{v}_k$ | $w_{ijk}$ | $w_{ijk} \delta \bar{v}_i \delta \bar{v}_j \delta \bar{v}_k$ | $i$ | $j$ | $k$ | $\delta \bar{v}_i \delta \bar{v}_j \delta \bar{v}_k$ | $w_{ijk}$ | $w_{ijk} \delta \bar{v}_i \delta \bar{v}_j \delta \bar{v}_k$ |
| 3 | 3 | 3 | $7.26 \times 10^{-5}$ | $2.79 \times 10^{-2}$ | $2.02 \times 10^{-6}$ | 1 | 1 | 1 | $-4.11 \times 10^{-4}$ | $9.08 \times 10^{-4}$ | $-3.73 \times 10^{-7}$ |
| 4 | 4 | 4 | $6.37 \times 10^{-3}$ | $3.06 \times 10^{-4}$ | $1.95 \times 10^{-6}$ | 1 | 2 | 2 | $-2.02 \times 10^{-6}$ | $7.75 \times 10^{-2}$ | $-1.56 \times 10^{-7}$ |
| 1 | 2 | 1 | $2.88 \times 10^{-5}$ | $8.39 \times 10^{-3}$ | $2.42 \times 10^{-7}$ | 2 | 2 | 1 | $-2.02 \times 10^{-6}$ | $7.75 \times 10^{-2}$ | $-1.56 \times 10^{-7}$ |
| 1 | 1 | 2 | $2.88 \times 10^{-5}$ | $8.38 \times 10^{-3}$ | $2.42 \times 10^{-7}$ | 2 | 1 | 2 | $-2.02 \times 10^{-6}$ | $7.74 \times 10^{-2}$ | $-1.56 \times 10^{-7}$ |
| 2 | 1 | 1 | $2.88 \times 10^{-5}$ | $8.38 \times 10^{-3}$ | $2.42 \times 10^{-7}$ | 4 | 4 | 1 | $-2.56 \times 10^{-3}$ | $5.46 \times 10^{-5}$ | $-1.40 \times 10^{-7}$ |
| 2 | 2 | 2 | $1.42 \times 10^{-7}$ | $7.16 \times 10^{-1}$ | $1.01 \times 10^{-7}$ | 1 | 4 | 4 | $-2.56 \times 10^{-3}$ | $5.46 \times 10^{-5}$ | $-1.40 \times 10^{-7}$ |
| 4 | 4 | 2 | $1.79 \times 10^{-4}$ | $4.06 \times 10^{-4}$ | $7.28 \times 10^{-8}$ | 4 | 1 | 2 | $-7.18 \times 10^{-5}$ | $1.23 \times 10^{-4}$ | $-8.86 \times 10^{-9}$ |
| 2 | 4 | 4 | $1.79 \times 10^{-4}$ | $4.06 \times 10^{-4}$ | $7.28 \times 10^{-8}$ | 2 | 1 | 4 | $-7.18 \times 10^{-5}$ | $1.23 \times 10^{-4}$ | $-8.86 \times 10^{-9}$ |

| -2, 100 mM NaCl, 6 mM MgCl<sub>2</sub> |  |  |  |  |  |  |  |  |  |  |  |
| --- | --- | --- | --- | --- | --- | --- | --- | --- | --- | --- | --- |
| Positive terms |  |  |  |  |  | Negative terms |  |  |  |  |  |
| $i$ | $j$ | $k$ | $\delta \bar{v}_i \delta \bar{v}_j \delta \bar{v}_k$ | $w_{ijk}$ | $w_{ijk} \delta \bar{v}_i \delta \bar{v}_j \delta \bar{v}_k$ | $i$ | $j$ | $k$ | $\delta \bar{v}_i \delta \bar{v}_j \delta \bar{v}_k$ | $w_{ijk}$ | $w_{ijk} \delta \bar{v}_i \delta \bar{v}_j \delta \bar{v}_k$ |
| 4 | 4 | 4 | $5.26 \times 10^{-3}$ | $4.21 \times 10^{-4}$ | $2.21 \times 10^{-6}$ | 2 | 2 | 2 | $-2.80 \times 10^{-5}$ | $8.77 \times 10^{-2}$ | $-2.46 \times 10^{-6}$ |
| 2 | 2 | 4 | $1.60 \times 10^{-4}$ | $1.37 \times 10^{-2}$ | $2.20 \times 10^{-6}$ | 2 | 4 | 4 | $-9.18 \times 10^{-4}$ | $2.39 \times 10^{-3}$ | $-2.19 \times 10^{-6}$ |
| 4 | 2 | 2 | $1.60 \times 10^{-4}$ | $1.37 \times 10^{-2}$ | $2.20 \times 10^{-6}$ | 4 | 4 | 2 | $-9.18 \times 10^{-4}$ | $2.39 \times 10^{-3}$ | $-2.19 \times 10^{-6}$ |
| 2 | 4 | 2 | $1.60 \times 10^{-4}$ | $1.36 \times 10^{-2}$ | $2.18 \times 10^{-6}$ | 4 | 2 | 4 | $-9.18 \times 10^{-4}$ | $2.14 \times 10^{-3}$ | $-1.97 \times 10^{-6}$ |
| 4 | 1 | 2 | $4.04 \times 10^{-4}$ | $3.97 \times 10^{-3}$ | $1.61 \times 10^{-6}$ | 1 | 2 | 2 | $-7.06 \times 10^{-5}$ | $2.30 \times 10^{-2}$ | $-1.63 \times 10^{-6}$ |
| 2 | 1 | 4 | $4.04 \times 10^{-4}$ | $3.97 \times 10^{-3}$ | $1.61 \times 10^{-6}$ | 2 | 2 | 1 | $-7.06 \times 10^{-5}$ | $2.30 \times 10^{-2}$ | $-1.63 \times 10^{-6}$ |
| 2 | 4 | 1 | $4.04 \times 10^{-4}$ | $3.97 \times 10^{-3}$ | $1.60 \times 10^{-6}$ | 4 | 4 | 1 | $-2.32 \times 10^{-3}$ | $6.98 \times 10^{-4}$ | $-1.62 \times 10^{-6}$ |
| 1 | 4 | 2 | $4.04 \times 10^{-4}$ | $3.97 \times 10^{-3}$ | $1.60 \times 10^{-6}$ | 1 | 4 | 4 | $-2.32 \times 10^{-3}$ | $6.98 \times 10^{-4}$ | $-1.62 \times 10^{-6}$ |

| -2, 20 mM NaCl, 6 mM MgCl<sub>2</sub> |  |  |  |  |  |  |  |  |  |  |  |
| --- | --- | --- | --- | --- | --- | --- | --- | --- | --- | --- | --- |
| Positive terms |  |  |  |  |  | Negative terms |  |  |  |  |  |
| $i$ | $j$ | $k$ | $\delta \bar{v}_i \delta \bar{v}_j \delta \bar{v}_k$ | $w_{ijk}$ | $w_{ijk} \delta \bar{v}_i \delta \bar{v}_j \delta \bar{v}_k$ | $i$ | $j$ | $k$ | $\delta \bar{v}_i \delta \bar{v}_j \delta \bar{v}_k$ | $w_{ijk}$ | $w_{ijk} \delta \bar{v}_i \delta \bar{v}_j \delta \bar{v}_k$ |
| 4 | 4 | 4 | $6.13 \times 10^{-3}$ | $9.54 \times 10^{-4}$ | $5.85 \times 10^{-6}$ | 4 | 4 | 1 | $-3.90 \times 10^{-3}$ | $6.69 \times 10^{-5}$ | $-2.61 \times 10^{-7}$ |
| 3 | 3 | 3 | $1.34 \times 10^{-4}$ | $4.17 \times 10^{-2}$ | $5.59 \times 10^{-6}$ | 1 | 4 | 4 | $-3.90 \times 10^{-3}$ | $6.69 \times 10^{-5}$ | $-2.61 \times 10^{-7}$ |
| 4 | 4 | 2 | $5.58 \times 10^{-5}$ | $7.83 \times 10^{-4}$ | $4.37 \times 10^{-8}$ | 1 | 1 | 1 | $-1.58 \times 10^{-3}$ | $6.56 \times 10^{-5}$ | $-1.04 \times 10^{-7}$ |
| 2 | 4 | 4 | $5.58 \times 10^{-5}$ | $7.82 \times 10^{-4}$ | $4.36 \times 10^{-8}$ | 3 | 3 | 1 | $-3.05 \times 10^{-4}$ | $8.68 \times 10^{-5}$ | $-2.65 \times 10^{-8}$ |
| 1 | 2 | 1 | $2.26 \times 10^{-5}$ | $1.53 \times 10^{-3}$ | $3.46 \times 10^{-8}$ | 1 | 3 | 3 | $-3.05 \times 10^{-4}$ | $4.11 \times 10^{-5}$ | $-1.25 \times 10^{-8}$ |
| 1 | 1 | 2 | $2.26 \times 10^{-5}$ | $1.53 \times 10^{-3}$ | $3.46 \times 10^{-8}$ | 1 | 2 | 2 | $-3.23 \times 10^{-7}$ | $3.58 \times 10^{-2}$ | $-1.16 \times 10^{-8}$ |
| 2 | 1 | 1 | $2.26 \times 10^{-5}$ | $1.53 \times 10^{-3}$ | $3.45 \times 10^{-8}$ | 2 | 2 | 1 | $-3.23 \times 10^{-7}$ | $3.58 \times 10^{-2}$ | $-1.16 \times 10^{-8}$ |
| 4 | 1 | 1 | $2.48 \times 10^{-3}$ | $5.17 \times 10^{-6}$ | $1.28 \times 10^{-8}$ | 2 | 1 | 2 | $-3.23 \times 10^{-7}$ | $3.58 \times 10^{-2}$ | $-1.15 \times 10^{-8}$ |

| -2, 20 mM NaCl, 0 mM MgCl<sub>2</sub> |  |  |  |  |  |  |  |  |  |  |  |
| --- | --- | --- | --- | --- | --- | --- | --- | --- | --- | --- | --- |
| Positive terms |  |  |  |  |  | Negative terms |  |  |  |  |  |
| $i$ | $j$ | $k$ | $\delta\bar{v}_i\delta\bar{v}_j\delta\bar{v}_k$ | $w_{ijk}$ | $w_{ijk}\delta\bar{v}_i\delta\bar{v}_j\delta\bar{v}_k$ | $i$ | $j$ | $k$ | $\delta\bar{v}_i\delta\bar{v}_j\delta\bar{v}_k$ | $w_{ijk}$ | $w_{ijk}\delta\bar{v}_i\delta\bar{v}_j\delta\bar{v}_k$ |
| 4 | 4 | 4 | $1.39 \times 10^{-3}$ | $1.34 \times 10^{-2}$ | $1.86 \times 10^{-5}$ | 4 | 4 | 1 | $-9.88 \times 10^{-4}$ | $1.34 \times 10^{-2}$ | $-1.32 \times 10^{-5}$ |
| 1 | 4 | 1 | $7.03 \times 10^{-4}$ | $1.34 \times 10^{-2}$ | $9.42 \times 10^{-6}$ | 1 | 4 | 4 | $-9.88 \times 10^{-4}$ | $1.34 \times 10^{-2}$ | $-1.32 \times 10^{-5}$ |
| 4 | 1 | 1 | $7.03 \times 10^{-4}$ | $1.34 \times 10^{-2}$ | $9.42 \times 10^{-6}$ | 4 | 1 | 4 | $-9.88 \times 10^{-4}$ | $1.34 \times 10^{-2}$ | $-1.32 \times 10^{-5}$ |
| 1 | 1 | 4 | $7.03 \times 10^{-4}$ | $1.34 \times 10^{-2}$ | $9.42 \times 10^{-6}$ | 1 | 1 | 1 | $-5.00 \times 10^{-4}$ | $1.34 \times 10^{-2}$ | $-6.70 \times 10^{-6}$ |
| 5 | 5 | 5 | $5.96 \times 10^{-3}$ | $5.56 \times 10^{-4}$ | $3.31 \times 10^{-6}$ | 2 | 4 | 4 | $-2.51 \times 10^{-4}$ | $1.03 \times 10^{-2}$ | $-2.58 \times 10^{-6}$ |
| 3 | 4 | 4 | $6.57 \times 10^{-4}$ | $3.59 \times 10^{-3}$ | $2.36 \times 10^{-6}$ | 4 | 4 | 2 | $-2.51 \times 10^{-4}$ | $1.03 \times 10^{-2}$ | $-2.58 \times 10^{-6}$ |
| 4 | 4 | 3 | $6.57 \times 10^{-4}$ | $3.59 \times 10^{-3}$ | $2.36 \times 10^{-6}$ | 2 | 2 | 2 | $-8.15 \times 10^{-6}$ | $3.14 \times 10^{-1}$ | $-2.56 \times 10^{-6}$ |
| 2 | 1 | 4 | $1.78 \times 10^{-4}$ | $1.03 \times 10^{-2}$ | $1.84 \times 10^{-6}$ | 3 | 1 | 4 | $-4.67 \times 10^{-4}$ | $3.59 \times 10^{-3}$ | $-1.68 \times 10^{-6}$ |

**Table S5.** Optimized forward and backward inverse rate constants corresponding to the elementary reactive steps for the samples studied.

| Sample | $k_{12}^{-1}$ | $k_{21}^{-1}$ | $k_{13}^{-1}$ | $k_{31}^{-1}$ | $k_{14}^{-1}$ | $k_{41}^{-1}$ | $k_{23}^{-1}$ | $k_{32}^{-1}$ | $k_{24}^{-1}$ | $k_{42}^{-1}$ | $k_{34}^{-1}$ | $k_{43}^{-1}$ | $k_{15}^{-1}$ | $k_{51}^{-1}$ | $k_{25}^{-1}$ | $k_{52}^{-1}$ | $k_{35}^{-1}$ | $k_{53}^{-1}$ | $k_{45}^{-1}$ | $k_{54}^{-1}$ |
| --- | --- | --- | --- | --- | --- | --- | --- | --- | --- | --- | --- | --- | --- | --- | --- | --- | --- | --- | --- | --- |
| +1, 300 mM NaCl | 66.3 | 533 | - | - | - | - | - | - | - | - | 11.8e4 | 363 | 0.85 | 1.75 | - | - | 556 | 9.91 | - | - |
| +1, 100 mM NaCl | - | - | - | - | 4.97 | 4.86 | 66.1e3 | 548 | 17.7e2 | 95.4 | 1.84 | 0.28 | - | - | - | - | - | - | - | - |
| +1, 20 mM NaCl | 0.55 | 1.64 | - | - | 16.4e2 | 15.4 | 27.1e3 | 627 | - | - | - | - | - | - | - | - | - | - | - | - |
| +1, 20 mM NaCl, 0 mM MgCl<sub>2</sub> | - | - | 4.64 | 121 | - | - | 25.1e2 | 109 | 731 | 368 | 4.31 | 0.37 | - | - | - | - | - | - | - | - |
| -1, 100 mM NaCl | 4.27 | 39.0 | 1.45 | 41.2 | 77.6e2 | 82.7 | 433 | 11.2e4 | - | - | - | - | - | - | - | - | - | - | - | - |
| -2, 300 mM NaCl | 0.40 | 3.74 | - | - | 539 | 10.8 | 21.8e3 | 518 | - | - | - | - | - | - | - | - | - | - | - | - |
| -2, 100 mM NaCl | 2.20 | 8.32 | - | - | 0.71 | 0.42 | 109 | 382 | 35.0e3 | 78.6e3 | - | - | - | - | - | - | - | - | - | - |
| -2, 20 mM NaCl | 0.55 | 12.9 | 52.5e3 | 494 | 180 | 15.6 | 84.5e2 | 18.6e2 | - | - | - | - | - | - | - | - | - | - | - | - |
| -2, 20 mM NaCl, 0 mM MgCl<sub>2</sub> | - | - | 10.4 | 12.7 | 0.0005 | 0.0005 | 4.65 | 1.17 | - | - | 75.0e3 | 91.7e3 | 87.2e3 | 285 | - | - | - | - | - | - |

Inverse rate constants defined in Eq. (4), listed in units of milliseconds. Error bars are < 1% of reported values (see Statistical Uncertainty Analysis below).

**Table S6.**
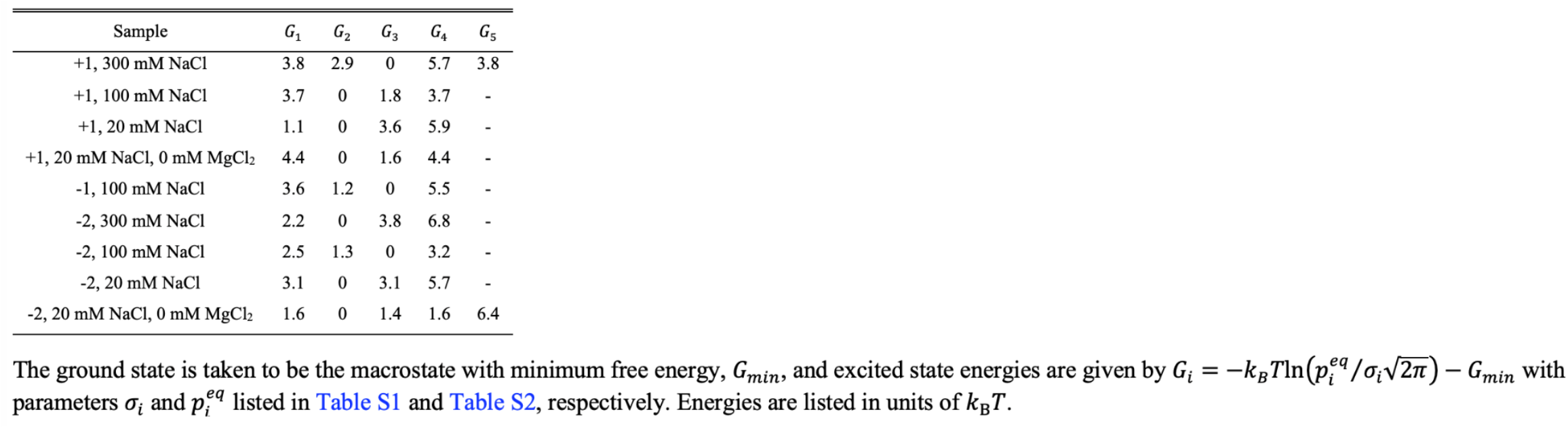
Relative free energy minima.

| Sample | $G_1$ | $G_2$ | $G_3$ | $G_4$ | $G_5$ |
| --- | --- | --- | --- | --- | --- |
| +1, 300 mM NaCl | 3.8 | 2.9 | 0 | 5.7 | 3.8 |
| +1, 100 mM NaCl | 3.7 | 0 | 1.8 | 3.7 | - |
| +1, 20 mM NaCl | 1.1 | 0 | 3.6 | 5.9 | - |
| +1, 20 mM NaCl, 0 mM MgCl<sub>2</sub> | 4.4 | 0 | 1.6 | 4.4 | - |
| -1, 100 mM NaCl | 3.6 | 1.2 | 0 | 5.5 | - |
| -2, 300 mM NaCl | 2.2 | 0 | 3.8 | 6.8 | - |
| -2, 100 mM NaCl | 2.5 | 1.3 | 0 | 3.2 | - |
| -2, 20 mM NaCl | 3.1 | 0 | 3.1 | 5.7 | - |
| -2, 20 mM NaCl, 0 mM MgCl<sub>2</sub> | 1.6 | 0 | 1.4 | 1.6 | 6.4 |

**Table S7.** Free energies of activation corresponding to the elementary reactive steps.

| Sample | $G_{12}^\ddagger$ | $G_{21}^\ddagger$ | $G_{13}^\ddagger$ | $G_{31}^\ddagger$ | $G_{14}^\ddagger$ | $G_{41}^\ddagger$ | $G_{23}^\ddagger$ | $G_{32}^\ddagger$ | $G_{24}^\ddagger$ | $G_{42}^\ddagger$ | $G_{34}^\ddagger$ | $G_{43}^\ddagger$ | $G_{15}^\ddagger$ | $G_{51}^\ddagger$ | $G_{25}^\ddagger$ | $G_{52}^\ddagger$ | $G_{35}^\ddagger$ | $G_{53}^\ddagger$ | $G_{45}^\ddagger$ | $G_{54}^\ddagger$ |
| --- | --- | --- | --- | --- | --- | --- | --- | --- | --- | --- | --- | --- | --- | --- | --- | --- | --- | --- | --- | --- |
| +1, 300 mM NaCl | 4.4 | 6.4 | - | - | - | - | - | - | - | - | 11.8 | 6.1 | 0 | 0.7 | - | - | 6.5 | 6.2 | - | - |
| +1, 100 mM NaCl | - | - | - | - | 2.9 | 2.9 | 12.4 | 7.6 | 8.8 | 1.9 | 1.9 | 0 | - | - | - | - | - | - | - | - |
| +1, 20 mM NaCl | 0 | 1.1 | - | - | 8.0 | 3.2 | 10.8 | 7.0 | - | - | - | - | - | - | - | - | - | - | - | - |
| +1, 20 mM NaCl, 0 mM MgCl<sub>2</sub> | - | - | 2.5 | 5.8 | - | - | 8.8 | 5.7 | 7.6 | 6.9 | 2.4 | 0 | - | - | - | - | - | - | - | - |
| -1, 100 mM NaCl | 1.1 | 3.3 | 0 | 3.3 | 8.6 | 4.0 | 5.7 | 11.2 | - | - | - | - | - | - | - | - | - | - | - | - |
| -2, 300 mM NaCl | 0 | 2.2 | - | - | 7.2 | 3.3 | 10.9 | 7.1 | - | - | - | - | - | - | - | - | - | - | - | - |
| -2, 100 mM NaCl | 1.7 | 3.0 | - | - | 0.5 | 0 | 5.5 | 6.8 | - | - | - | - | - | - | - | - | - | - | - | - |
| -2, 20 mM NaCl | 0 | 3.2 | 11.5 | 6.8 | 5.8 | 3.3 | 9.6 | 8.1 | - | - | - | - | - | - | - | - | - | - | - | - |
| -2, 20 mM NaCl, 0 mM MgCl<sub>2</sub> | - | - | 9.9 | 10.1 | 0 | 0 | 9.1 | 7.8 | - | - | 18.8 | 19.0 | 19.0 | 13.2 | - | - | - | - | - | - |

The activation energies are determined from $G_{ij}^\ddagger = -k_B T \ln(k_{ij}/k_{max})$ where $k_{max}$ is the fastest rate constant determined from the analysis. Energies are listed in units of $k_B T$ .

**Table S8.**
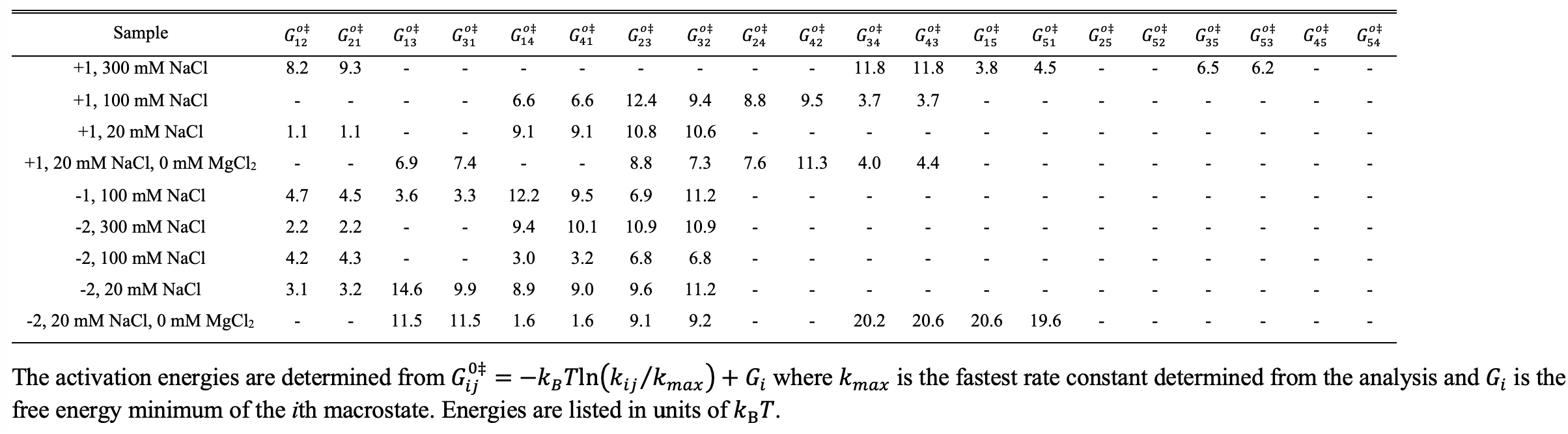
Free energies of activation corresponding to the elementary reactive steps.

## Salt concentration dependence of the local conformational macrostates of the +1 and -2 (iCy3)_2_ dimer-labeled ss-dsDNA fork constructs

In Fig. S1 we present our salt-concentration-dependent results for the +1 (iCy3)_2_ dimer-labeled ss-dsDNA construct. At the highest salt concentrations that we studied (300 mM NaCl, 6 mM MgCl_2_; Fig. S1*A*), nearly all the population resides in macrostate S_3_ (LHU,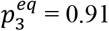), and the remaining populations in macrostates S_1_ (PO), S_2_ (WC), S_4_ (RHU) and S_5_ (to be assigned). The two-point TCF exhibits a somewhat faster overall decay [with mean relaxation time 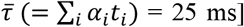 compared to the ‘physiological’ salt sample (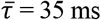, Fig. S1*B*). However, the three-point TCF exhibits significantly greater initial amplitude at the resolution of *T*_*w*_ = 10 ms, indicating the presence of highly correlated three-point transition pathways at elevated salt concentrations. Indeed, Table S4 of the SI shows that the largest ‘pathway terms’ at *T*_*w*_ = 10 ms involve recurrent measurements of macrostates S_5_ and S_4_ (i.e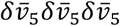 and 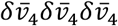 and transitions between macrostates S_1_ and S_5_ (e.g., 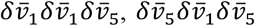 etc.). The CPs (bottom row) show that macrostate S_5_ is a transient intermediate species; the transition probability *p*_15_ increases in the sub-millisecond time scales, reaches its maximum value at ∼20 ms, and decays to its equilibrium value 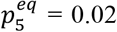, with a time scale of ∼100 ms. The optimized kinetic network scheme shown in Fig. S2*A* indicates that macrostate S_5_ is directly connected to macrostates S_1_ and S_3_, and the free energy minimum of S_5_ has an equal value to that of S_1_ (see Table S6 of the SI).

At ‘physiological’ salt concentrations (100 mM NaCl, 6 mM MgCl_2_, Fig. S1*B*), the major component is macrostate S_2_ (WC,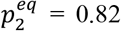) and the second most populated component is macrostate S_3_ (LHU,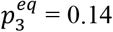). When the salt concentration is decreased below ‘physiological’ (20 mM NaCl and 6 mM MgCl_2_, Fig. S1*C*), the major component is macrostate S_2_ (WC,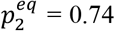), and the second most populated component is S_1_ (PO,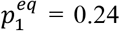). At this reduced salt concentration, the S_3_ macrostate exists as a trace population 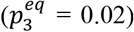.The two-point TCF exhibits faster overall dynamics at these low salt concentrations 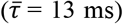,which is primarily due to significant lowering of the transition barriers between the two most thermodynamically favored macrostates, S_2_ (WC) and S_1_ (PO) (Fig. S2*E* and S2*F*). Table S4 shows that the largest pathway terms contributing to the initial amplitude of the three-point TCF involve recurrent measurements of macrostates S_1_, S_3_ and S_4_ and transitions between macrostates S_1_ and S_2_.

Finally, elimination of divalent magnesium ions at low monovalent sodium ion concentration (20 mM NaCl, 0 mM MgCl_2_, Fig. S1*D*) leads to a reversal in the relative stabilities of macrostates S_1_ and S_3_ (see Figs. S2*G* and S2*H*), resulting in a pattern that significantly resembles that seen under ‘physiological’ salt conditions. This comparison, with and without magnesium ions, permits us to isolate the effects of magnesium near the +1 position. When magnesium is eliminated at low sodium concentration, the major component is macrostate S_2_ (WC,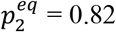) and the second most populated component is macrostate S_3_ (LHU,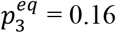). In the absence of magnesium, macrostate S_1_ (PO) exists as a trace population 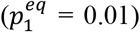.The two-point TCF exhibits faster overall dynamics in comparison to the physiological sample 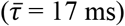,although somewhat slower than in the presence of magnesium. The very small initial amplitude of the three-point TCF (Fig. S1*D*) indicates a near cancellation between positive and negative three-point pathway terms (see Table S3). Finally, the CPs at physiological and zero magnesium concentrations (Figs. S1*B* and S1*D*, respectively) show that macrostate S_3_ (LHU) is a transient intermediate under these conditions.

**Figure S1.**
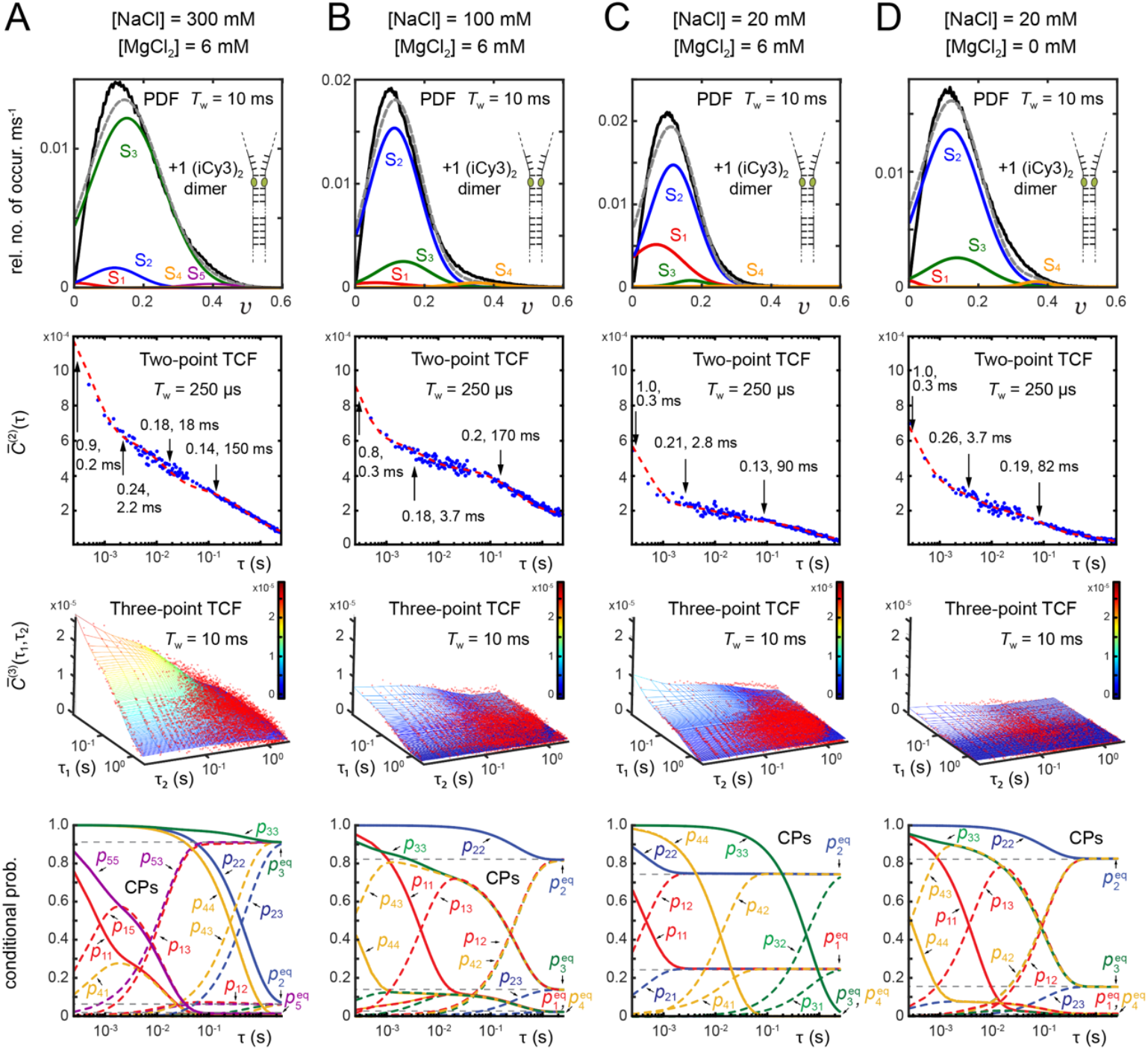
Results of kinetic network model analyses applied to PS-SMF measurements of the +1 (iCy3)_2_ dimer-labeled ss-dsDNA construct in 10 mM Tris at pH 8.0 and (***A***) 300 mM NaCl, 6 mM MgCl_2_; (***B***) 100 mM NaCl, 6 mM MgCl_2_; (***C***) 20 mM NaCl, 6 mM MgCl_2_; (***D***) 20 mM NaCl, 0 mM MgCl_2_. Formatting and color schemes are the same as described in Fig. 6 of the main text. Optimized values of the Gaussian parameters are listed in Table S1 and Table S2.

**Figure S2.**
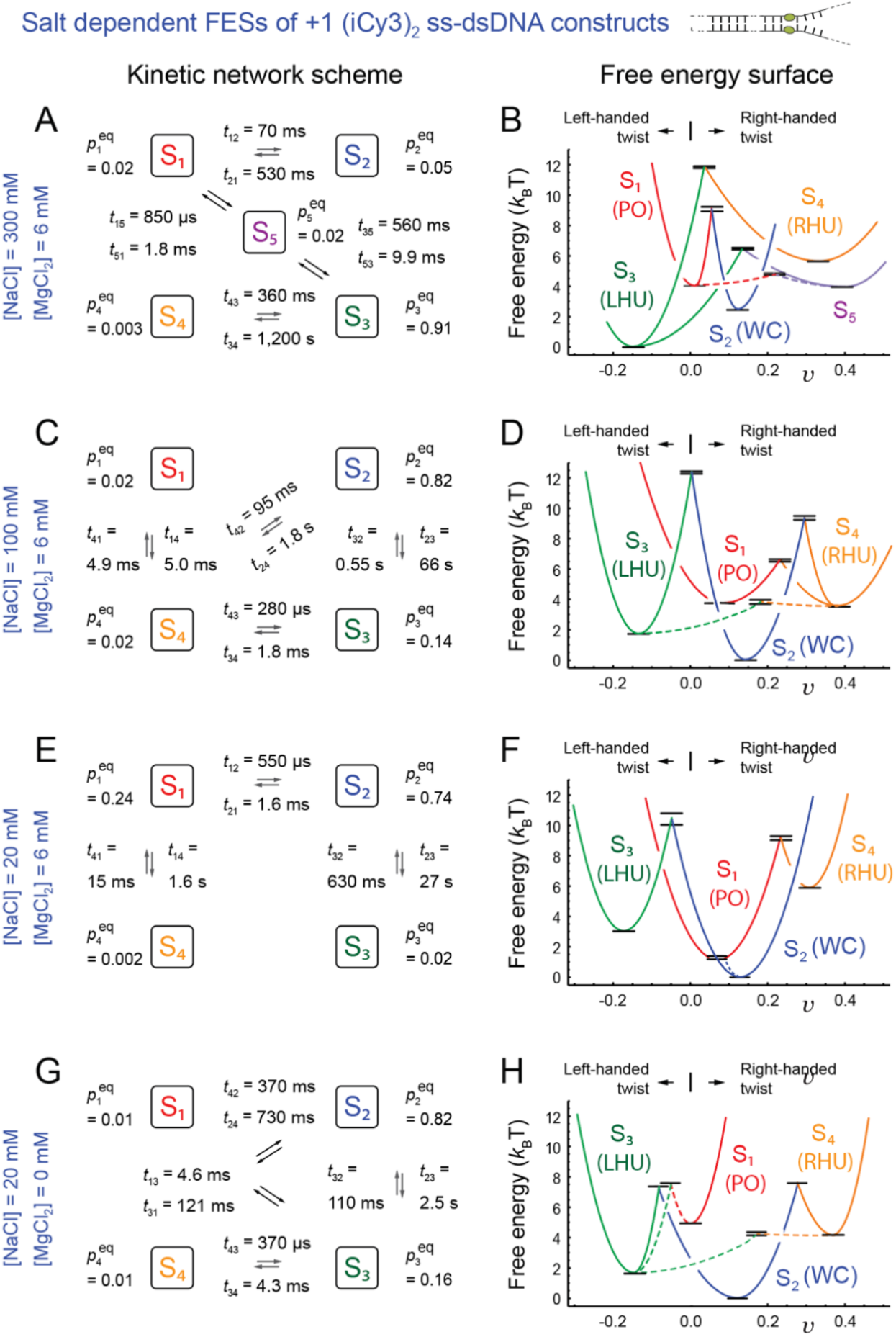
Kinetic network schemes (***A, C, E, G***) and free energy surfaces (FESs) (***B, D, F, H***) from the analyses of the +1 (iCy3)_2_ dimer-labeled ss-dsDNA fork construct in 10 mM Tris at pH 8.0 and (***A, B***) 300 mM NaCl, 6 mM MgCl_2_; (***C, D***) 100 mM NaCl, 6 mM MgCl_2_; (***E, F***) 20 mM NaCl, 6 mM MgCl_2_; (***G, H***) 20 mM NaCl, 0 mM MgCl_2_. Formatting and color schemes are the same as described in Fig. 7 of the main text. Optimized values of the time constants are listed in Table S5, free energy minima in Table S6, and transition energies in Table S7.

In Fig. S3, we present our salt concentration-dependent results of the -2 (iCy3)_2_ dimer-labeled ss-dsDNA fork construct. The optimized kinetic network schemes for each set of salt concentrations and the associated FESs are presented in Fig. S4. As discussed in Sect. 3.1, at the ‘physiological’ salt concentrations (100 mM NaCl, 6 mM MgCl_2_, Fig. S3*B*) the major component is macrostate S_3_ (LHU,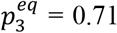) and the second most populated component is macrostate S_2_ (WC,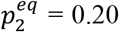), which is opposite to the population order observed for the +1 construct. The two-point TCF exhibits a relatively fast overall decay 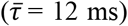,indicating that the transition barriers are relatively low in comparison to the +1 construct. Moreover, the three-point TCF exhibits a weak and negative initial amplitude at resolution *T*_*w*_ = 10 ms, which is consistent with relatively low transition barriers between macrostates. The effects of increasing or decreasing salt concentrations on the -2 construct are opposite to those we observed with the +1 construct. At the highest salt concentrations (300 mM NaCl, 6 mM MgCl_2_; Fig. S3*A*), the major component is macrostate S_2_ (WC,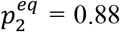) and the second component is macrostate S_1_ (PO, 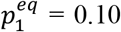). Macrostates S_3_ (LHU) and S_4_ (RHU) are populated at trace levels at these high salt concentrations. The two-point TCF exhibits a slower overall decay 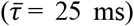 compared to ‘physiological’ concentrations, indicating that at least some transition barriers are elevated.

Decreasing the monovalent sodium ion concentration from the ‘physiological’ value to 20 mM NaCl and 6 mM MgCl_2_, (Fig. S3*C*), resulted in the transfer of nearly all the population to macrostate S_2 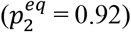,_with only trace populations occurring in macrostates S_1_, S_3_ and S_4_. The two-point TCF at the lower salt concentrations decays relatively rapidly 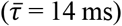,albeit slightly slower than under ‘physiological’ conditions. Elimination of divalent magnesium ions at the low sodium ion concentration significantly alters the equilibrium distribution of macrostates. At these lowest salt concentrations (20 mM NaCl, 0 mM MgCl_2_, Fig. S3*D*), the major component is macrostate S_2_ (WC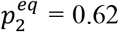) and the remaining population is shared nearly equally between macrostates S_1_ (PO,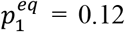), S_3_ (LHU,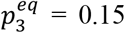) and S_4_ (RHU,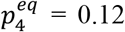). A trace population is also present in macrostate S_5_. The two-point TCF at the lowest salt concentrations exhibits a significantly slower overall decay 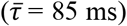 than for any of the other constructs and salt conditions that we studied. The slow decay of the two-point TCF indicates that the transition barriers are much higher than for the other samples, which is corroborated by the relatively large and positive initial amplitude of the three-point TCF at resolution *T*_*w*_ = 10 ms. Table S2 of the SI shows that the three-point pathway terms that dominate on the shortest time scales involve transitions between macrostates S_1_ and S_4_, and the optimized network scheme shown in Fig. S4*G* indicates that nearly all five of the macrostates may directly interconvert between one another.

**Figure S3.**
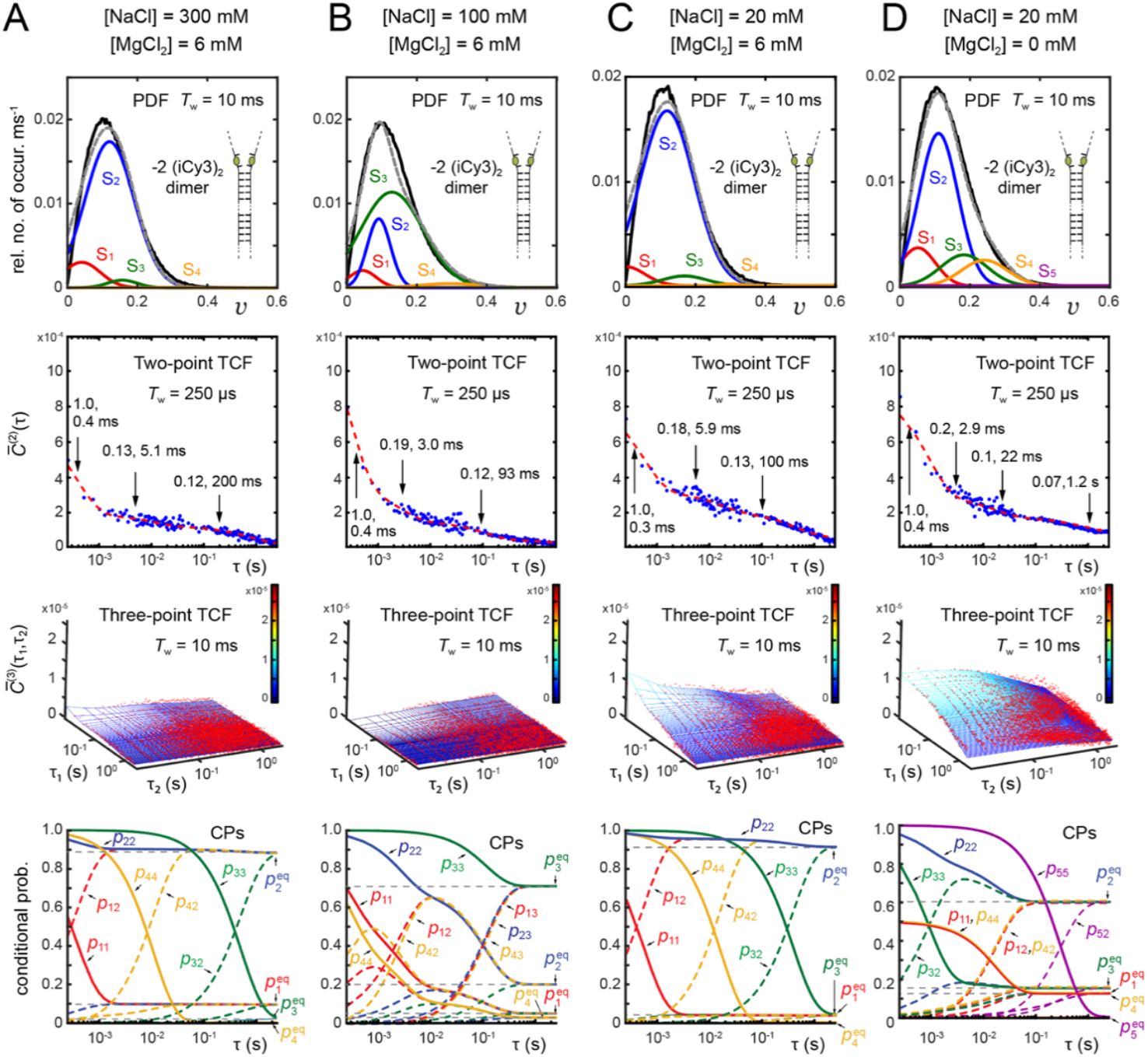
Results of kinetic network model analyses applied to PS-SMF measurements of the -2 (iCy3)_2_ dimer-labeled ss-dsDNA construct in 10 mM Tris at pH 8.0 and (***A***) 300 mM NaCl, 6 mM MgCl_2_; (***B***) 100 mM NaCl, 6 mM MgCl_2_; (***C***) 20 mM NaCl, 6 mM MgCl_2_; (***D***) 20 mM NaCl, 0 mM MgCl_2_. Formatting and color schemes are the same as those in Fig. 6 of the main text.

**Figure S4.**
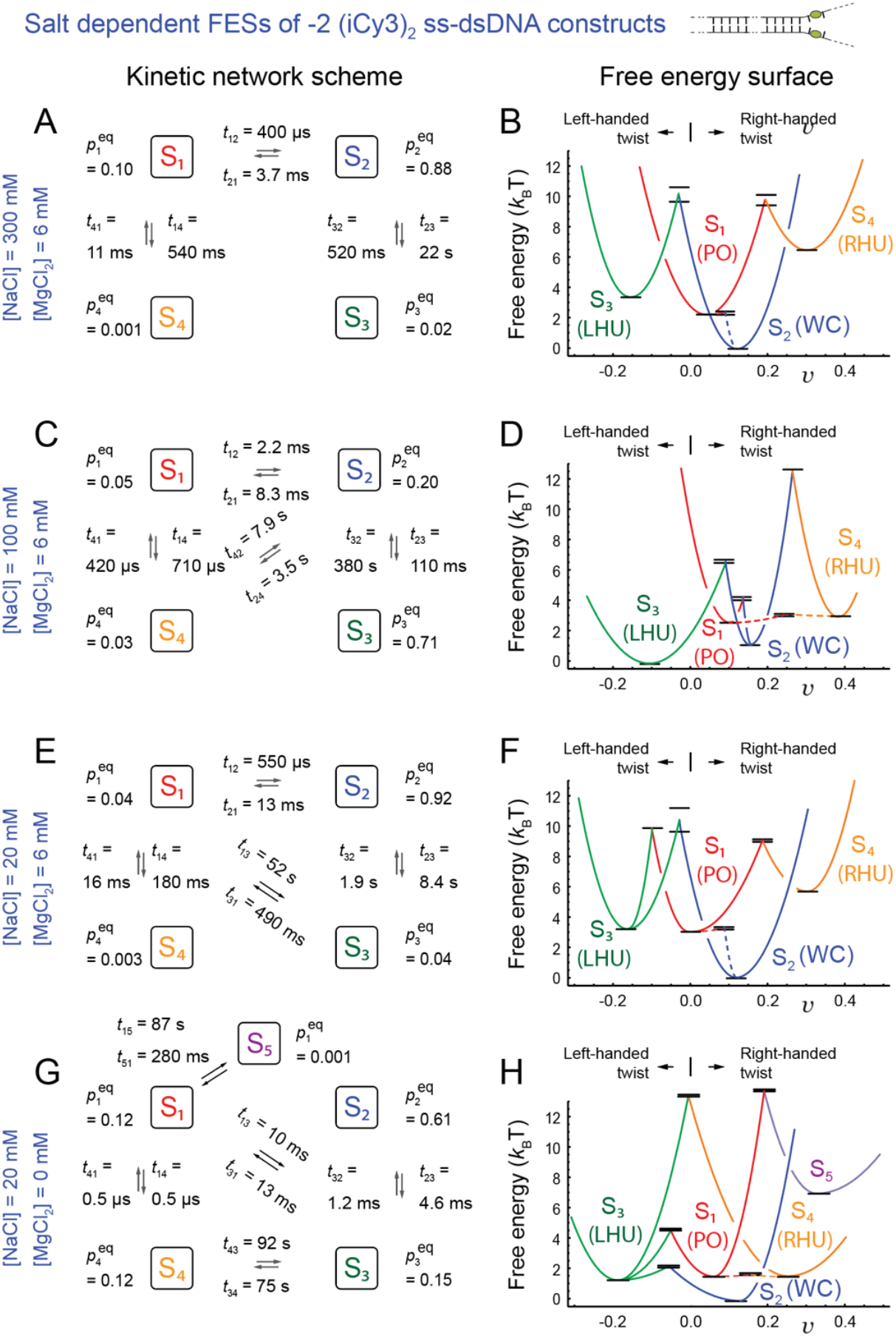
Kinetic network schemes (***A, C, E, G***) and free energy surfaces (FESs) (***B, D, F, H***) from the analyses of the -2 (iCy3)_2_ dimer-labeled ss-dsDNA fork construct in 10 mM Tris at pH 8.0 and (***A, B***) 300 mM NaCl, 6 mM MgCl_2_; (***C, D***) 100 mM NaCl, 6 mM MgCl_2_; (***E, F***) 20 mM NaCl, 6 mM MgCl_2_; (***G, H***) 20 mM NaCl, 0 mM MgCl_2_. Formatting and color schemes are the same as those in Fig. 7 of the main text.

## Iterative multiparameter optimization procedure

We performed an iterative multi-parameter optimization procedure, like the one described by Phelps *et al*. (1,2) and Israels *et al*. (3), to find the set of time constants and equilibrium probabilities that best match the experimentally-derived PDF [*P*(*v*)], two-point TCF 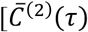C2 function] and three-point TCF 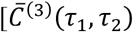, C3 function]. To assess the quality of the agreement between the solutions to the master equation and the statistical functions constructed from our experimental data, we introduced the global least squares error function 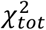,as described in Sect. 2.5 of the main text. For each of the three statistical functions, *q* [= PDF, C2, C3], we define the component error function

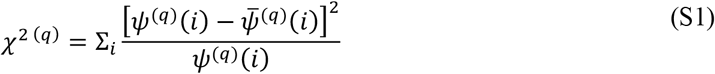

where *ψ*^(*q*)^(*i*) and 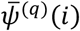 are the experimental and simulated functions, respectively, and the index *i* enumerates the data points within the coordinate space of the *q*th function.

The total error function, 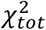.is the sum of weighted contributions from the three statistical functions.

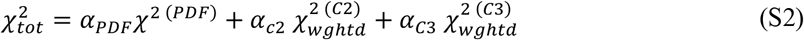

In Eq. (S2), the parameter *α*_*q*_ is a scaling factor that balances the relative contribution from each statistical function. Typical magnitudes (i.e., maxima) of the PDF, C2 and C3 functions are ∼10^-1^, ∼10^-3^ and ∼10^-5^, respectively. The factors *α*_*q*_ were thus adjusted to ensure that each of the terms in Eq. (S2) contributed equally to the globally minimized value of 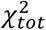, which reflects optimal agreement between all three simulated and experimental functions.

In addition to the above considerations, it was necessary to account for the time-dependent variations in the data point densities, signal amplitudes and noise of the C2 and C3 functions, which each span multiple decades in their respective time variables. To optimize the agreement between simulated and experimental functions over the full range of time scales, we used Eq. (S2) to calculate the ‘weighted’ functions 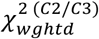 in which we assigned larger weights to regions of data with relatively sparse point density, high signal amplitude and low noise.

We first describe our procedure for determining 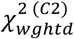 for the C2 function, which is a one-dimensional function of the time delay, τ. The C2 function contains *N*_*data*_ (= ∼200) data points and spans *N*_*dec*_ (= ∼4) decades in time. We divided each decade into *N*_*seg*_ (= ∼10) ‘segments,’ which we enumerated using the index *j*. In Eq. (S3), we define the *j*-dependent ‘inverse weight function’ 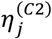 which sums over the *N*_*j*_ data points contained within the *j*th segment.

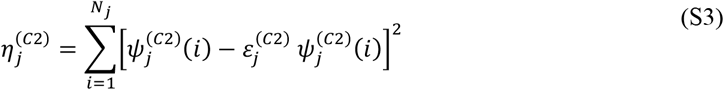

Here 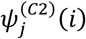 is the *i*th data point contained within the *j*th segment, and 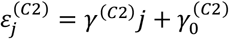 is a linear function of *j* with constant slope *γ*^(*C*2)^ and offset 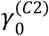 The slope *γ*^(*C*2)^ was chosen to account for the rate of increasing point density, decreasing amplitude, and increasing noise across segments within a given decade.

Finally, we expanded the dimension of the *j*-dependent function 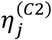 (with dimension *N*_*seg*_ · *N*_*dec*_) to include values corresponding to each of the *N*_*data*_ data points. We thus calculated the weighted error function used in Eq. (S2) according to

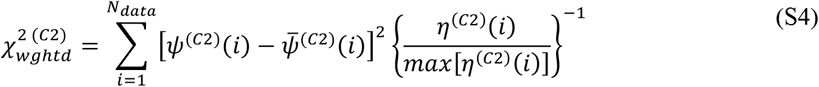

where the index *i* spans each of the data points contained within the C2 function. In Eq. (S4), the factor {*η*^(*C*2)^(*i*)/*max*4*η*^(*C*2)^(*i*)]} ^−1^ assigns a greater weight to regions of comparatively low point density, higher signal amplitude and low noise.

To determine 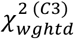 for the C3 function we performed a calculation like that for 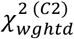.

However, we generalized the procedure to account for the dependence of the C3 function on the two time variables, τ_1_ and τ_2_. The number of data points contained by the C3 function is 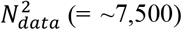. Each time variable spans *N*_*dec*_ (= ∼2.5) decades, and each decade was divided into *N*_*seg*_ (∼10) segments. The two-dimensional function was thus partitioned into 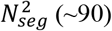 segments, each of which are enumerated using the indices *j* and *j*^′.^ The two-dimensional inverse weight function is thus defined

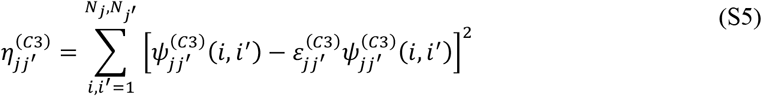

In Eq. (S5), 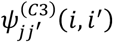 is the *ii*^′^th data point contained within the *jj*^′^ th segment, and 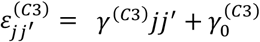 is a linear joint function of *j* and *j*^′^ with constant slope *γ*^(*C*3)^ and constant offset 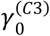 like that implemented for the C2 function.

We expanded the dimension of the *jj*^′^-dependent function 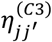 (with dimension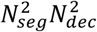) to include values corresponding to each of the 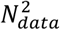 data points. We thus calculated the weighted error function used in Eq. (S2) according to

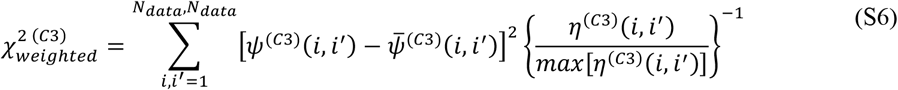

We determined optimal values for the input parameters 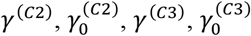 and *N*_*seg*_ for the weight functions 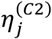 and 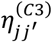 given by Eq. (S3) and Eq. (S5), respectively, and for the scaling factors *α*_*q*_ given by Eq. (S2) as an initialization phase of our optimization calculations. The sensitivity of these calculations to the most significant regions of the data was thus optimized.

We carried out our multiparameter optimization calculation in two stages. In the first stage, we used a custom-designed genetic algorithm (GA) to broadly search the parameter space and to obtain a set of parameters that produced qualitatively good agreement between simulated and experimental functions. In the second stage of the calculation, the output solutions obtained from the GA were fed into a commercial multivariable search function (*patternsearch*, MATLAB, The MathWorks, USA), which refined the solutions by further reducing the value of 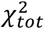.Further details of this procedure are given below.

The GA that we implemented is very similar to the one used by Israels *et al*. (3). However, to fit the C2 and C3 functions of the current work, which span several decades in time, some modifications were introduced. The GA is an optimization procedure that is inspired by biological evolution. A schematic workflow diagram for the GA is shown in Fig. S5. The calculation proceeds through a set of successive hierarchical ‘iterations’ (indexed by *N*). Within each iteration are associated multiple ‘generations’ (indexed by *n*), and within each generation are associated a population of initial ‘guesses’ for the set of input parameters (represented by differently colored circles in Fig. S5*A*). The input parameters are: (*i*) the mean visibilities of the macrostates, 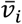 (*ii*) the Gaussian widths of the macrostates, σ_*i*_; and (*iii*) the inverse rate constants of transitions between macrostates, 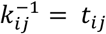 where *i, j* ∈ {1,2,3,4}.

Within each iteration, the calculation proceeds through multiple sequential generations, which are ‘evolved’ according to the workflow outlined in Fig. S5*B* and Fig. 5S*C*. The first generation, *G*_1_, typically contains ∼2,000 initial guesses of the parameters 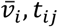,and σ_*i*_, which are each assigned values using a random number generator (see Fig. S5*B*). Because the parameters, *t*_*ij*_, span multiple decades, their values are assigned by sampling logarithmically the full range of time scales. Each individual guess within the first generation is ranked according to its agreement with the experimental functions, which is quantified using the value of 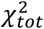 [Eq. (S2)]. From this relatively large initial generation (*G*_1_) is selected the top 10% (∼200) of individuals exhibiting the most favorable agreement between simulated and experimental functions.

**Figure S5.**
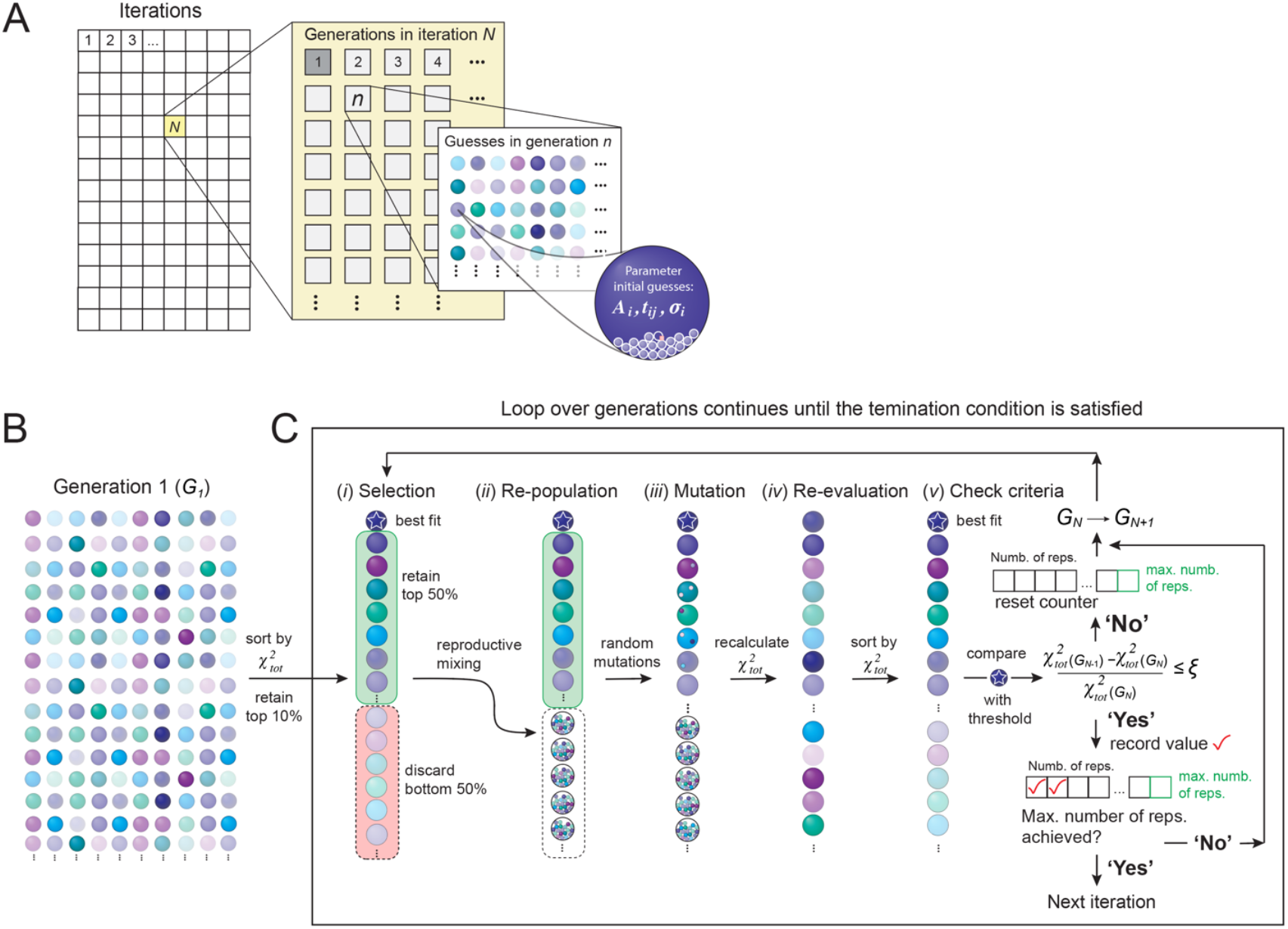
Workflow diagram of the multiparameter optimization procedure used in this work. See SI text for further details. Adapted from ref. (3).

The population of ∼200 individuals is next refined through a series of ‘evolutionary steps,’ as illustrated in Fig. S5*C*. In the ‘selection’ step (*i*), the top 50% of individuals are retained and the bottom 50% of individuals are discarded. In the ‘reproductive mixing’ step (*ii*), the top ranking individuals from step (*i*) are used to create ‘progeny,’ which involves the random exchange (or ‘mixing’) of parameters between individuals. In the ‘mutation’ step (*iii*), the parameters of all but the top two individuals are randomly varied to generate ‘diversity.’ In the re-evaluation step (*iv*), the refined population is ranked by updating the values of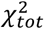 . In the final step (*v*), the value of 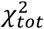 for the top individual (marked with a blue star) is compared to the highest ranked value obtained from the previous generation. When the value of the top individual has improved by an amount that meets or exceeds the set threshold value *ξ* (= ∼0.01), the updated 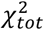 value is recorded, and the refined generation is used as input to repeat the next cycle. Generational cycles that do not satisfy the threshold criterion are tallied and their refined populations are also used as input for the next cycle, but their 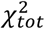 values are discarded. After ∼10 successive generational cycles occur for which the threshold criterion is satisfied, the final solution is recorded, and the next iteration is begun.

The GA is typically carried for a duration of 100 – 500 iterations. A subset of these solutions exhibit qualitatively good agreement between all three simulated and experimental functions. These output solutions obtained from the GA are then fed into the multivariable problem solver, *patternsearch* (MATLAB, The MathWorks, USA), to arrive at a final optimized solution.

**Figure S6.**
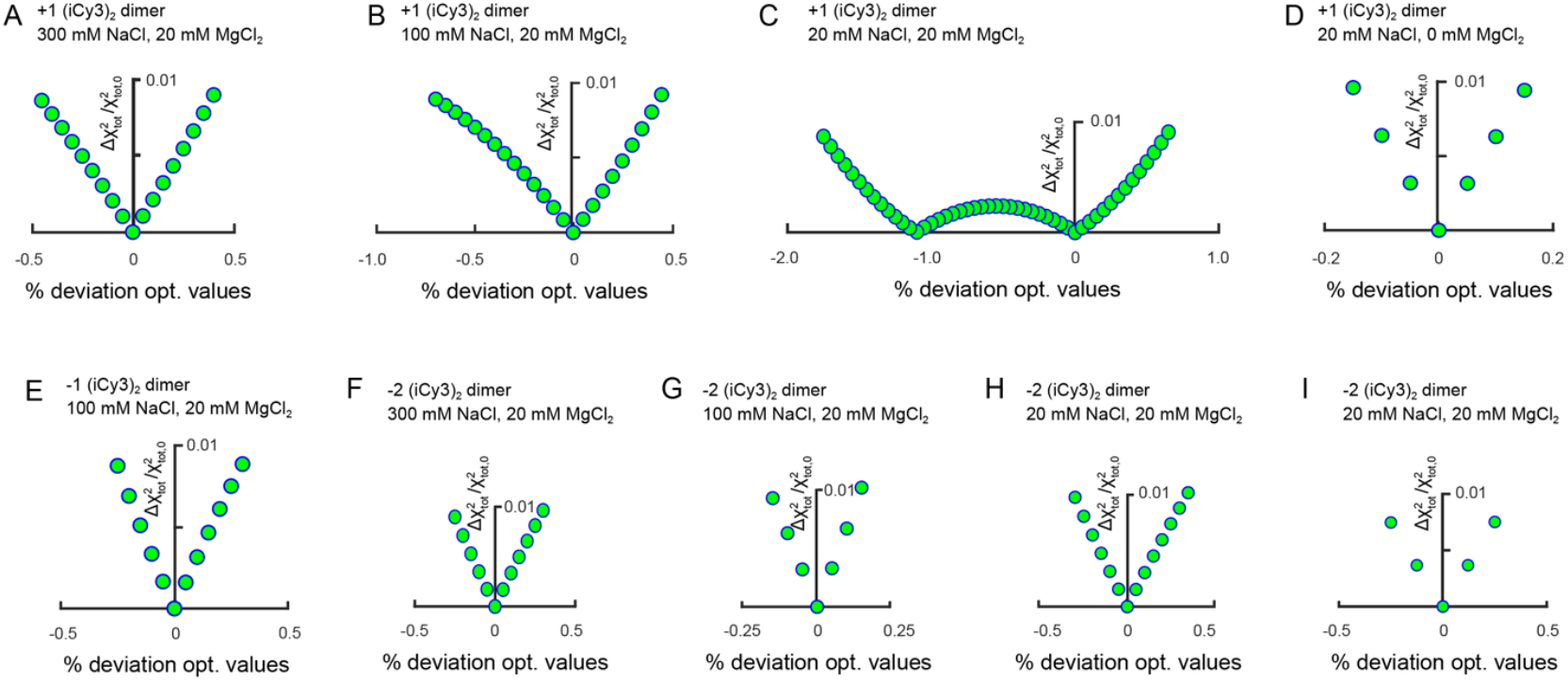
Relative deviation of the error function 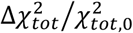 from the optimized value 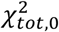,as a function of the time constants and the mean visibility parameter uncertainties for each of (iCy3)_2_ dimer-labeled ss-dsDNA constructs and salt concentrations studied in this work. (***A***) +1 (iCy3)_2_ dimer at 300 mM NaCl and 6 mM MgCl_2_. (***B***) +1 (iCy3)_2_ dimer at 100 mM NaCl and 6 mM MgCl_2_. (***C***) +1 (iCy3)_2_ dimer at 20 mM NaCl and 6 mM MgCl_2_. (***D***) +1 (iCy3)_2_ dimer at 20 mM NaCl and 0 mM MgCl_2_. (***E***) -1 (iCy3)_2_ dimer at 100 mM NaCl and 6 mM MgCl_2_. (***F***) -2 (iCy3)_2_ dimer at 300 mM NaCl and 6 mM MgCl_2_. (***G***) -2 (iCy3)_2_ dimer at 100 mM NaCl and 6 mM MgCl_2_. (***H***) -2 (iCy3)_2_ dimer at 20 mM NaCl and 6 mM MgCl_2_. (***I***) -2 (iCy3)_2_ dimer at 20 mM NaCl and 0 mM MgCl_2_. We note that the calculation for the sample shown in panel (*C*) exhibits two nearly stable minima within the error bar of ∼±1%.

## Statistical Uncertainty Analysis

We performed an analysis of the statistical uncertainty for the free parameters used in our kinetic network model for each of the (iCy3)_2_ dimer-labeled ss-dsDNA fork constructs at various salt conditions. This analysis is based on calculating the relative deviation of the global error function, 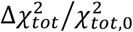 , from the optimized value, 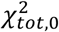, as a function of the time constants and the mean visibility parameter uncertainties. The global error function dependence on the parameter uncertainties for the various constructs and salt conditions, shown in Fig. S6, demonstrate that the optimized values we obtained each correspond to a stable minimum. We assign upper bounds to the statistical uncertainties of these values to a 1% deviation of the total error function relative to the corresponding optimized value. From this analysis we obtain upper bounds of ∼ ±1% for the error bars of the free parameters listed in Table S1 and Table S5.

## Footnotes

1 The ‘physiological’ salt concentrations referred to in this paper are typically [NaCl] = 100 mM and [MgCl_2_] = 6 mM for *in vitro* studies. These are not the actual intra-cellular salt concentrations, which vary substantially with changing environmental conditions. Rather these values correspond to a set of salt concentrations that result in the same protein-DNA binding constants in *in vitro* model systems as those measured in living *E. coli* cells

